# Green Fluorescent Carbon Dots as Targeting Probes for LED-Dependent Bacterial Killing

**DOI:** 10.1101/2021.03.26.437036

**Authors:** Jenny Samphire, Yuiko Takebayashi, Stephen A. Hill, Nicholas Hill, Kate J. Heesom, Philip A. Lewis, Dominic Alibhai, Eilis C. Bragginton, Josephine Dorh, Neciah Dorh, James Spencer, M. Carmen Galan

## Abstract

The emergence of antimicrobial resistance represents a significant health and economic challenge worldwide. The slow pace of antibacterial discovery necessitates strategies for optimal use of existing agents, including effective diagnostics able to drive informed prescribing; and development of alternative therapeutic strategies that go beyond traditional small-molecule approaches. Thus, the development of novel probes able to target bacteria for detection and killing, and that can pave the way to effective theranostic strategies, is of great importance. Here we demonstrate that metal-free green-emitting fluorescent carbon dots (FCDs) synthesized from glucosamine HCl and *m*-phenylenediamine, and featuring 2,5-deoxyfructosazine on a robust amorphous core, can label both Gram-positive (*Staphylococcus aureus*) and Gram-negative (*Escherichia coli, Klebsiella pneumoniae, Pseudomonas aeruginosa*) bacterial pathogens within 10 minutes of exposure. Moreover, effective killing of Gram-positive and -negative bacteria can be induced by combining FCD treatment with irradiation by LED light in the visible range. Cell-based, electron microscopy and Tandem Mass Tag (TMT) proteomic experiments indicate that FCD administration in combination with LED exposure gives rise to local heating, ROS production, and membrane- and DNA-damage, suggesting multiple routes to FCD-mediated bacterial killing. Our data identify FCDs as materials that combine facile synthesis from low-cost precursors with labelling and light-dependent killing of clinically important bacterial species, and that thus warrant further exploration as the potential bases for novel theranostics.

## 1. Introduction

Bacterial infections affect most people at some point in their lives and, while antibiotic treatments exist for most common pathogens, their effectiveness is threatened by growing antimicrobial resistance (AMR). This problem is exacerbated by the rising overuse of broad spectrum antibiotics combined with the slowing development of new antibiotics.^[1]^ In an effort to control and prevent growing antibiotic resistance, rapid diagnostic tools that can be used to detect the presence of bacteria are key to avoid antibiotic treatments being unnecessarily prescribed.^1b^

Fluorescent labelling has been previously implemented to investigate antibiotic susceptibility, intracellular pathogenesis and detection of bacterial cell wall proteins.^[2]^ However, molecular dyes are usually expensive and predisposed to photobleaching. Fluorescent nanoprobes on the other hand can be designed to exhibit high stability, sensitivity and specificity for their desired target without the limitations of organic fluorophores and fluorescent proteins and thus these nanomaterials have found many applications in the areas of bioimaging, drug delivery and diagnostics.^[3]^ Furthermore, the fluorescence properties of molecular fluorophores can be affected upon binding to the target, while intrinsically fluorescent nanoparticles are rarely affected and can be more robust bioimaging tools.^[4]^

One particular class of fluorescent nanoparticles, carbon dots (CDs), have emerged as promising bioimaging probes due to their many advantages over molecular fluorophores and other fluorescent nanoparticles such as the analogous heavy-metal containing quantum dots.^[5]^ The synthesis of these water-soluble carbon-based nanodots is often practical and low-cost,^[6]^ while CDs tend to be chemically inert, photo-stable and generally non-toxic, making them ideal for biological applications.^[6f, 6h, 6i, 7]^ Examples of blue-emitting functionalized CDs have been reported to label bacteria.^[8]^, ^[9]^ However, the autofluorescence properties of many microorganisms^[10]^ overlap with the blue emission of the CDs used, meaning that the sensitivity of such probes is not high enough for practical use. In addition, parameters such as nanoparticle type, size, shape and surface functionalization have been shown to have significant effects on targeting ability, intracellular uptake and localization.^[11]^

Stemming from our interest in the synthesis of carbon-based water soluble fluorescent probes for bioimaging applications,^[6f, 6g, 6i]^ we recently developed a new class of green-fluorescent CDs (FCDs) derived from glucosamine and *m*-phenylenediamine (Scheme 1) that could selectively label and kill human cancer cells upon activation by illumination with visible LEDs.^[12]^ The unique targeting ability of this nanomaterial was attributed to the presence of 2,5-deoxyfructosazine on the FCD surface. 2,5-deoxyfructosazine is the product of 1,2-aminoaldose self-dimerisation during FCD synthesis,^[13]^ and analogous fructosazine, is itself a versatile molecule with anti-diabetic and anti-inflammatory properties.^[14]^

Based on these findings, we thus proposed that these green-emitting FCDs could be promising candidates to target live bacteria. Herein, we demonstrate that the unique surface functionality of these green-emitting FCDs can also be used to label both Gram-positive and Gram-negative bacteria. Furthermore, we also show the combination of FCD exposure with LED-activation to exert a bactericidal effect, and investigate the mechanism of bacterial killing action.

**Scheme 1:**
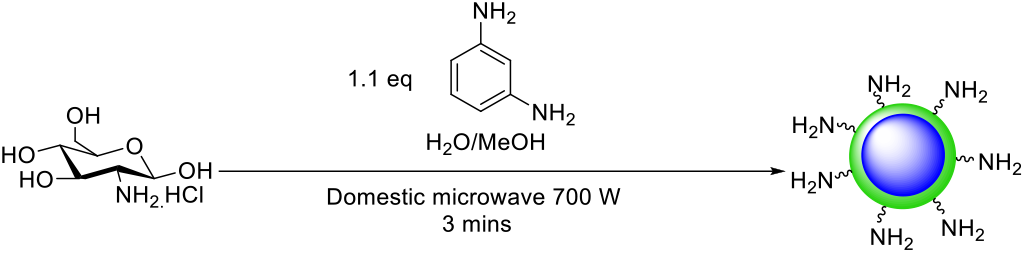
General synthetic approach to access green-emitting FCDs.

## 2. Results and Discussion

Green-fluorescent carbon dots (FCDs) surface functionalized with 2,5-deoxyfructosazine were obtained by a three-minute microwave-assisted one-pot reaction of glucosamine and *m*-phenylenediamine following a previously reported synthesis.^[12]^ To evaluate the ability of the FCD to label bacteria, different initial FCD concentrations (50-200 μg/mL) were incubated with four different bacterial species to assess the optimal labeling conditons over a range of time exposures (30-60 min). To that end, *Escherichia coli* (BW25113 strain), a Gramnegative bacterium; and *Staphylococcus aureus* (Newman strain), a Gram-positive bacterium; and further two Gram-negative species, *Pseudomonas aeruginosa* (PA01 strain) and *Klebsiella pneumoniae* (NCTC 5055 strain) were chosen as relevant microorganisms. Labelling of the bacteria was assessed using confocal microscopy. To our delight, all bacterial species evaluated showed labeling after as little as 10 min incubation with green FCDs (Figure 2) and at FCD concentrations as low as 25 μg/mL. As expected, fluorescence labeling increased with the amount of FCDs used. In general, optimal labelling was achieved with a 1 x10^8^ cfu/mL suspension of bacteria in phosphate buffered saline (PBS) incubated with 200 μg/mL of FCDs at room temperature for 30 min (see ESI Figure S1). To quantify bacterial labelling, fluorescent measurements of FCD-labeled bacterial suspensions were compared to a FCD calibration curve. Uniformly, FCDs were found to show increased labelling of Gram-negative bacteria when compared to the Gram-positive species *S. aureus*, which had the lowest levels of labelling (Figure 1).

**Figure 1:**
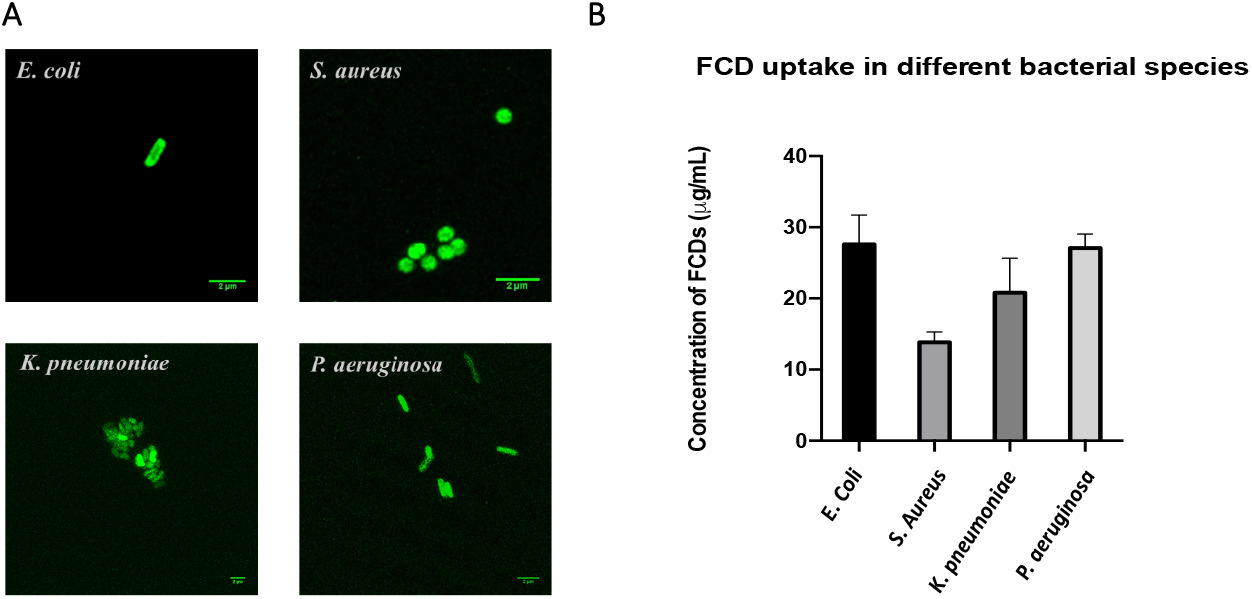
FCDs label Gram-positive and -negative Bacteria. A) Hyvolution confocal microscopy z-stack max-projected images of FCD labelled bacteria; *E. coli, S. aureus, P. aeruginosa* and *K. pneumoniae* and B) quantification of FCD labelling for each bacteria species after 10 min incubation with 200 μg/mL of FCDs at room temperature.

**Figure 2:**
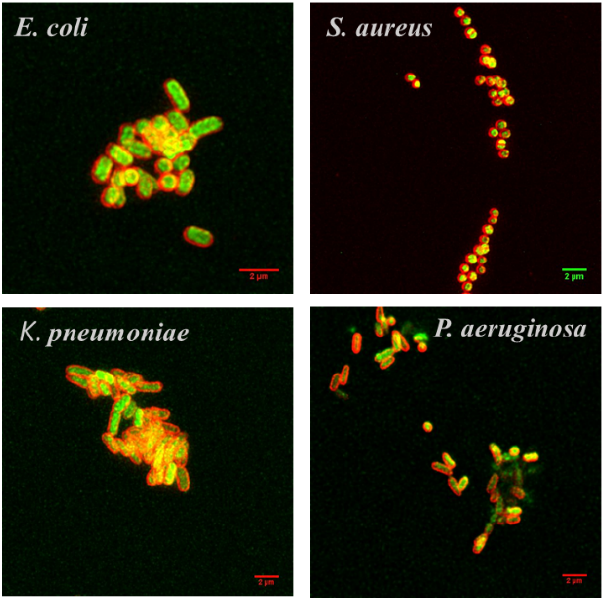
FCDs are Internalised by Gram-positive and -negative bacteria. Confocal image of different bacterial species incubated with membrane dye FM 4-64 (red) and FCDs (green).

Previous investigations of the chemical structure of the green-emitting FCDs^[12]^ revealed that the carbon-based probe contains a stable amorphous core, decorated with 2,5-deoxyfructosazine as the major surface component. It is however important to note that neither previously reported amine coated blue-emitting FCDs^[6i]^ or 2,5-deoxyfructosazine (itself non-fluorescent) isolated from green FCDs or from a commercial source, were able to fluorescently label the bacteria, demonstrating that the observed effects can be attributed to the properties of the green-emitting FCDs as a whole (see figures S1 and S2 in ESI).

To establish whether the FCDs are internalized within the bacteria, or remain associated with the cell surface, FCD-treated bacteria were then labelled with the membrane dye FM™ 4-64FX (Figure 2). FM™ 4-64FX is a lipophilic probe which has low fluorescence in aqueous media, but upon binding to plasma membranes fluoresces intensely in the infrared, allowing visualisation of the bacterial membrane and hence localisation of the green-emitting FCDs with respect to this.^[15]^ Confocal microscopy images of bacteria labeled with both FCDs (green) and FM™ 4-64FX (red, Figure 2) identified distinct patterns of staining for the two labels, confirmed that the FCDs are internalised within the bacteria and hence demonstrating their ability to penetrate the bacterial envelope.

Having confirmed that FCDs are internalized by bacteria, the toxicity of the green FCDs towards both the Gram-negative (*E. coli, K. pneumoniae, P. aeruginosa*) and Gram-positive (*S. aureus*) was next explored. Overall, little antibacterial activity was detected upon treatment with FCDs, even at high FCD concentrations (MIC > 1024 μg/mL), with the most notable effect being an increase in the duration of the lag phase (see ESI).

Previously, we have shown photothermal activation of green-emitting FCDs using low cost blue-light-emitting diodes (LEDs, λ_em_ = 460 nm).^[12]^ Irradiation by blue LEDs in the absence of FCDs did not induce toxic effects on bacteria, however, when LED irradiation was combined with FCD exposure, bactericidal activity was detected towards all four species tested (*E. coli, S. aureus, K. pneumoniae* and *P. aeruginosa* (ESI, figure S4)). Complete killing was reproducibly observed after treatment with 200 μg/mL FCDs with 4 hours of irradiation, and significant killing (>95%) could be observed after just 90 minute LED irradiation.

For two strains of *E. coli*, the antimicrobial effects of UV-B irradiation have been shown to be potentiated by fructosazine.^[16]^ To demonstrate that the bactericidal effects observed here were linked to the combination of FCD/LED treatment, and not due to either the presence of 2,5-deoxyfructosazine/LED, or 2,5-deoxyfructosazine alone, *E. coli* was treated with 2,5-deoxyfructosazine obtained from a commercial source at the same concentrations as those found on the FCDs, as estimated by NMR, and also exposed to LED irradition under the same conditions as previously used (See ESI figure S5 for details). Viable counts of treated cells showed that significant antibacterial effect could only be detected on exposure to the FCD/LED combination (*t*-test, *P* < 0.0001), with no effects observed, compared to controls, for cells treated with 2,5-deoxyfructosazine, either with or without LED illumination (*t*-test, *P* = 0.4254 and *P* = 0.3916 respectively) (Figure 3A).

**Figure 3:**
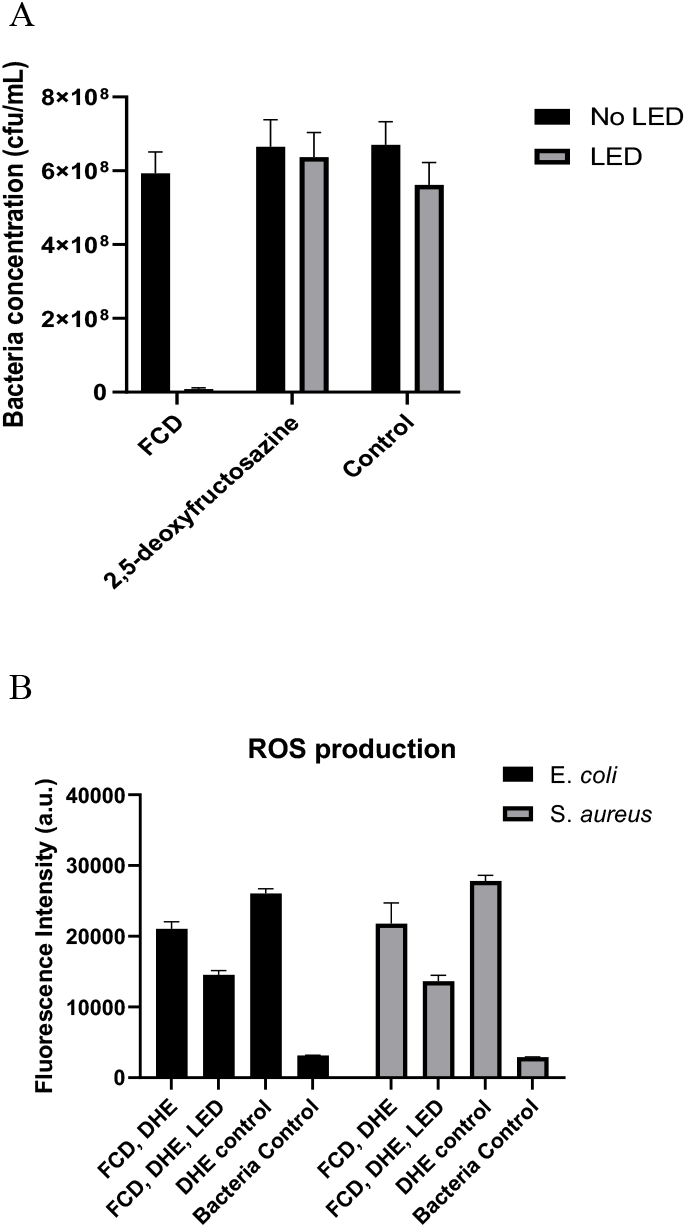
A) Antimicrobial Effects of Combined FCD Treatment and LED Illumination. Viable count shown as cfu/mL of *E. coli* cells treated with green FCDs (200 μg/mL), commercial 2,5-deoxyfructosazine (30 μg/mL) and controls with and without 90 min LED irradiation. Counts were averaged and shown with SD error bars (n=3). **B) FCD Treatment and LED Illumination elicits ROS production in Gram-positive and -negative bacteria.** Fluorescence intensity measurements show decrease in fluorescence at 460 nm (DHE emission) for *E. coli* and *S. aureus* incubated with 200 μg/mL FCDs and illuminated with LEDs (FCD, DHE, LED) or kept in the dark (FCD, DHE) for 90 min. Fluorescence intensities were averaged and shown with SD. Error bars (n=6).

Photothermal therapy (PTT) is a promising non-invasive therapeutic strategy, in which nanoparticles embedded within a pathogenic target generate heat, typically in response to exogenously applied light, for thermal ablation. Our previous study of green FCD interactions with cultured human cells^[12]^ demonstrated killing to be predominantly mediated by photothermal effects. Accordingly, the temperatures of bacterial suspensions incubated with 200 μg/mL and 800 μg/mL FCDs and exposed to LED irradiated for 60 min were measured, and found to be 3 °C higher (32 °C and 33 °C respectively) than those of cultures exposed to LED irradiation only, i.e. in the absence of FCDs.^[17]^ These data suggest that, consistent with our earlier investigation, photothermal activation of FCDs internalized within the bacteria might be responsible for at least some portion of the observed antimicrobial activity. However, we also sought to identify whether other mechanisms might contribute to the effects of FCD exposure upon bacteria. Reactive oxygen species (ROS) production is involved in the mechanism of action of multiple physical and chemical antibacterial agents and treatments,^[18]^ thus the superoxide indicator dihydroethidium (DHE, for which decrease in fluorescence on oxidation indicate ROS production) was used to monitor ROS formation in LED and FCD treated bacteria. For both *E. coli* and *S. aureus* the DHE’s fluorescence intensity for FCD-incubated (200 μg/mL) bacteria after 90 minutes of LED illumination was significantly lower by 46% (compared to DHE controls) than that of cultures treated with FCDs but without LEDs (*t*-test *P* = 0.0007 and *P* = 0.0008 respectively) (Figure 3B). This suggests that FCDs, when combined with LED irradiation, are eliciting ROS generation. Taken together, these data indicate that stress factors, such as ROS and temperature changes, are likely pertinent to the observed toxicity engendered by the LED/FCD combination.

Confocal microscopy (Figure 2) helped us demonstrate the ability of FCDs to penetrate the envelopes of both Gram-positive and -negative bacteria. To establish whether FCD exposure leads to membrane damage, and investigate the possibility that this may also contribute to the observed effects upon bacterial growth, scanning electron microscopy (SEM) was used to study the effects of LED and FCD exposure on *E. coli* and *S. aureus*. Bacterial cells exposed to 512 μg/mL FCDs, either with or without LED irradiation, indeed showed signs of envelope stress, evident in increased membrane blebbing and lysis. Importantly, LED irradiation alone did not induce these phenotypes, with these samples comparable to the control cells (Figure 4).

**Figure 4.**
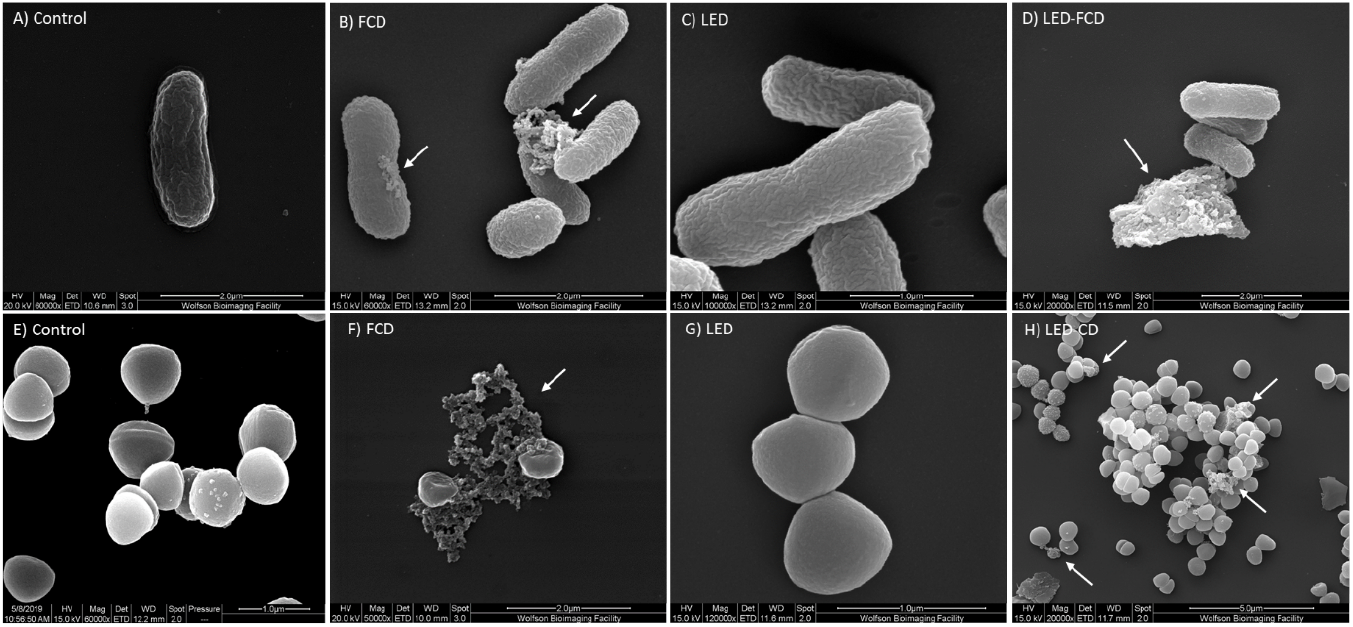
Representative SEM images of *E. coli* (A-D) and *S. aureus* (E-F) with FCD, LED and LED-FCD treatments. Signs of envelope stress/damage highlighted by arrows.

To further probe the cellular stress responses activated by FCD exposure, and the effects upon this of LED irradiation, *E. coli* and *S. aureus* cultures were grown in M9 minimal media in the presence or absence of 512 μg/mL FCDs and with and without LED irradiation, and analysed by Tandem Mass Tag (TMT) proteomics. Approximately 2,600 individual protein hits were obtained in each of the bacterial cultures analysed. From each sample (LED irradiated, FCD-treated or FCD-treated and LED irradiated), the total protein hits were compared against the control untreated sample. Changes in protein expression levels are summarised in Figure 5. The functions of proteins of interest were searched using PantherDB^44^.

**Figure 5.**
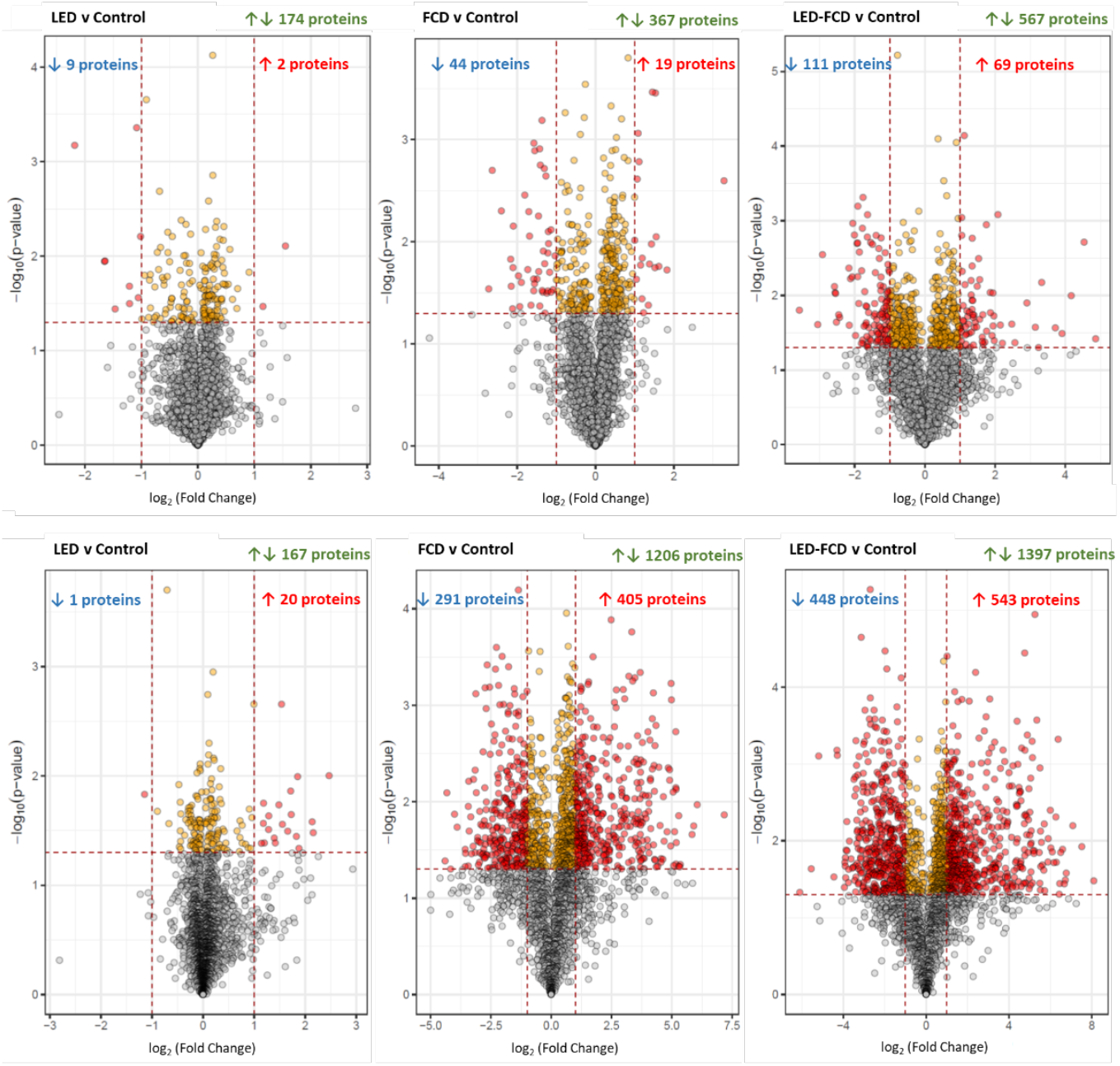
Volcano plots of (Top) *E. coli* and (Bottom) *S. aureus* treated with LED, FCD and LED-FCD. Overall number of proteins with changes in abundance levels indicated in green font. Grey circles represent proteins with differences in abundance levels (compared to control) that are not statistically significant, yellow circles represent those with statistical significance (p<0.05) and red circles represent those that are ≥ 2-fold different in abundance.

**Figure 6.**
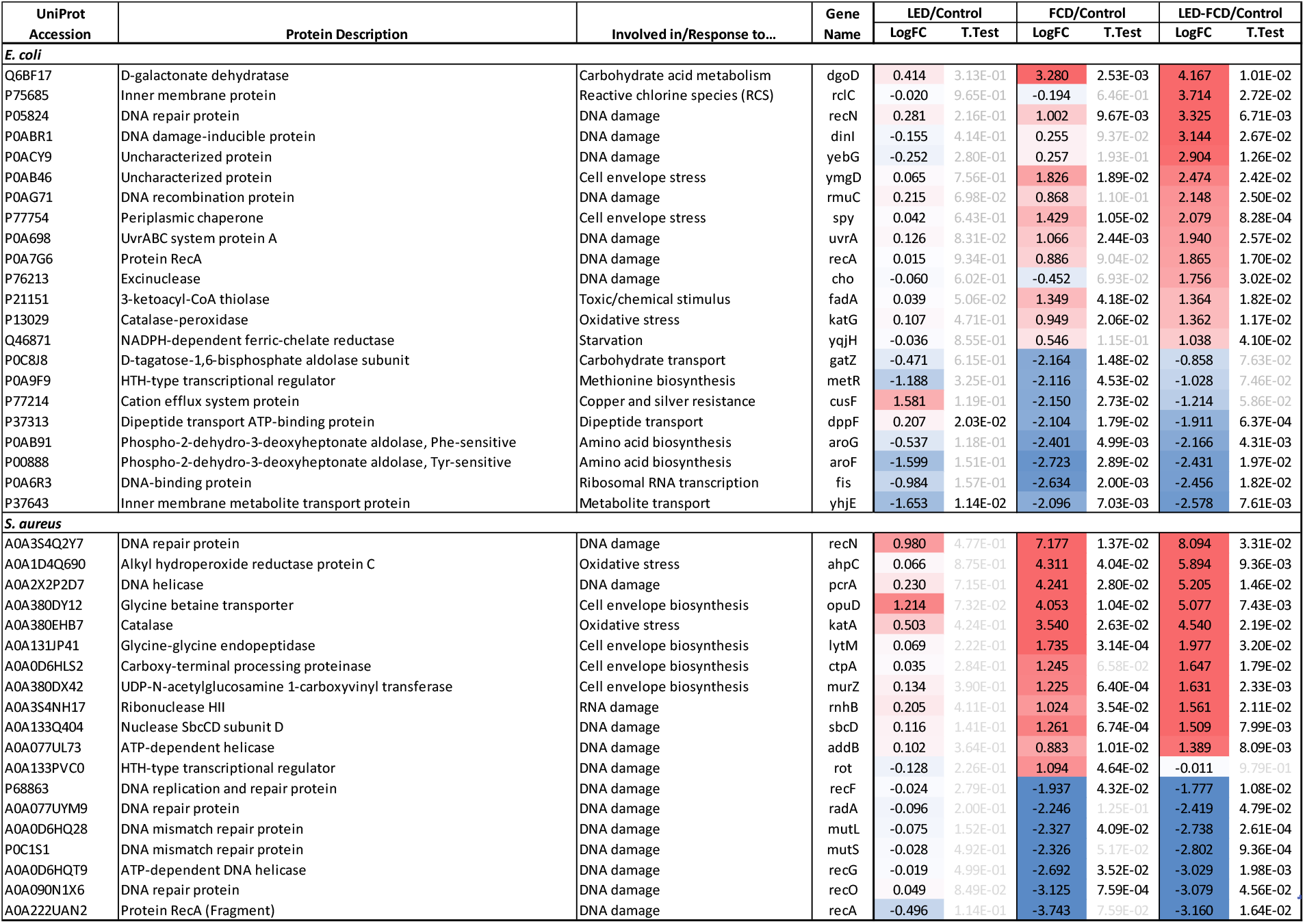
Summary of proteins of interest from proteomics analysis. LogFC, average log2 fold change (n=2). Heat map applied to LogFC values - shading in red indicate upregulation and blue indicate downregulation of protein abundance levels compared to control samples. Statistically significant values shown in black font (p<0.05) and insignificant values (p>0.05) in grey font as determined by *t*-test.

In both bacterial species, of the tested conditions LED irradiation had the least effect on protein expression levels (174 *E. coli* and 167 *S. aureus* proteins were identified as up- or down-regulated, p<0.05). Only 11 and 21 proteins in the two species, respectively, showed over 2-fold changes in expression level compared to the control. In *E. coli*, upregulated proteins included cation efflux protein (CusF), acetolactate synthetase (IlvG) and RNA helicase (DeaD), while various protein and amino acid transport proteins, such as the inner membrane metabolite transport protein YhjE and the Sec-independent protein translocase protein TatE, were downregulated. In *S. aureus*, 20 proteins were upregulated, including the amino acyl tRNA synthetases ValS and PheT, and only 1 protein, the elongation factor Tu fragment Tuf was downregulated by over 2-fold compared to the control. All proteins of interest are summarised in Figure 7 and the complete list of proteins with ≥2-fold changes in expression level is provided in the ESI.

Upon FCD exposure, the number of protein expression changes doubled in *E. coli* and increased by 7-fold in *S. aureus*. In *E. coli*, the protein showing the highest increase in expression level was D-galactonate dehydratase (DgoD), followed by a range of proteins involved in cellular and metabolic processes, some of which were indicative of DNA damage (UvrA, RecN), cell envelope stress (YmgD, Spy) and toxic/chemical stimulus (FadA). Downregulation of cellular, metabolic and localisation proteins was also observed, with at least two-log fold decreases in expression of enzymes involved in biosynthesis of important metabolic precursors such as amino acids (AroF, AroG, MetR), in carbohydrate metabolism (GatZ), ribosomal RNA transcription (Fis) and transport systems (CusF, DppF, YhjE). In *S. aureus*, the DNA repair protein RecN was the most upregulated protein, followed by hundreds of proteins involved in a vast range of cellular and metabolic pathways. Notable groups of upregulated proteins included those involved in responses to ROS exposure (KatA, AhpC), DNA damage (PcrA, SbcD, Rot) and RNA repair (RnhB). In terms of proteins involved in the response to cell envelope stress, various genome-wide transcriptional profiling studies of *S. aureus* treated with cell-wall-active antibiotics highlighted the upregulation of genes involved in cell envelope biogenesis^[19]^. Some of these corresponding proteins were also noted to be upregulated in the proteomics data, such as the glycine betaine transporter (OpuD), glycine-glycine endopeptidase (LytM), carboxyl-terminal processing proteinase (CtpA) and UDP-*N*-acetylglucosamine 1-caryboxylvinyl transferase (MurZ). Similarly, expression of over 200 enzymes was downregulated, with notable changes in translational, metabolite interconversion and nucleic acid metabolism proteins, including those involved in DNA repair (RecO, RecG, RecF and MutL).

With FCD treatment and LED irradiation, expression levels of a further ~200 proteins were affected, compared to the FCD-treated samples, for both bacterial species. In *E. coli*, multiple enzymes involved in DNA damage were upregulated compared to the FCD-only sample. Some of these (RecN, UvrA) were significantly upregulated on FCD exposure and further upregulated with LED irradiation; for others (RmuC, RecA) there was apparent upregulation on FCD exposure but the difference was not statistically significant without LED irradiation; and a third group (DinI, YebG, Cho) were little changed in the FCD-treated sample but showed significant upregulation with LED irradiation. Other proteins upregulated included those involved in the cellular response to reactive oxygen species (ROS, RclC), toxic/chemical stimuli (FadA), oxidative stress (KatG) and starvation (YqjH). In *S. aureus*, all of the proteins listed above (with the exception of Rot) were further upregulated, along with additional proteins involved in the cellular response to DNA damage (RecA, AddB). The majority of other upregulated enzymes were those involved in metabolite interconversion, followed by translation, transport and nucleic acid binding. Downregulation of the proteins involved in DNA repair listed above was maintained, and there was downregulation of additional proteins involved in DNA repair, such as MutS and RadA. In summary, the proteomic results suggest that exposure of *E. coli* to FCDs alone is associated with induction of responses to both membrane and DNA damage. In the case of *S. aureus*, the effect of FCD exposure upon the proteome is more profound, but also includes proteins involved in the response to ROS exposure. However, when FCD treatment is augmented with LED irradiation, in both organisms proteomic signatures are consistent with a response to multiple stressors that include both membrane and DNA damage, as well as elevated ROS-levels.

## 3. Conclusions

In summary, we have shown that green-emitting FCDs can be used to label both Gram-positive and Gram-negative bacteria within minutes of exposure at room temperature. Moreover, combining FCD treatment with LED-activation leads to effective bacteria cell killing. There are very few examples of FCD photothermal activation in antimicrobial photothermal therapy (PTT) ^[20]^ and most materials contain transition metals which can be activated with NIR light or other sensitizers for photodynamic therapy applications or do not achieve complete bacteria killing.^[21]^ We showed that the targeting properties exhibited by the FCDs are dependent upon the surface functionality on the nanomaterial with the presence of 2,5-deoxyfructosazine on the FCD surface being key and that the FCDs are internalized within the bacterial cell. Our investigations of the mechanism of bactericidal action suggest that upon FCD exposure, both *E. coli* and *S. aureus* experience membrane and DNA damage, as evidenced by both SEM and proteomic experiments, while combination of the FCD exposure with mild LED irradiation enhances DNA damage and induces additional effects including ROS production and local heating. In summary, our data show that these low cost metal-free bifunctional nanoprobes display unique physico-chemical and targeting properties that can be exploited as bioimaging tools, as well as in antimicrobial PTT and hence show great potential for further development as effective theranostic antimicrobial agents.

## 4. Materials and Methods

### Bacterial strains, media, growth conditions, suspensions and blue-LED irradiation

*Escherichia coli* BW 25113, *Klebsiella pneumoniae* NCTC 5055, *Pseudomonas aeruginosa* PA01 and *Staphylococcus aureus* Newman were used in this study. All four strains were grown on Nutrient agar/broth (Neogen) at 37°C overnight (18-20 h) unless otherwise stated. Bacterial suspensions were prepared from overnight plates in PBS to an OD600 of 0.8-1.0 (~1 x 10^9^ cfu/mL) unless otherwise stated. LED-irradiated samples were exposed to blue-LED strip lights (24 W, 12 V, λem= 460 nm) for durations specified below.

### FCD preparation

FCDs were prepared as previously described.^[12]^ Briefly, glucosamine.HCl and *m*-phenylenediamine were heated for 3 min in a domestic microwave (800W), re-suspended in water and filtered through a MWCO 10 kDa centrifugal concentrator. The solution was then reduced *in vacuo* and hydrolysed in water prior to use.

### Sample preparation for confocal imaging

10 μl of 5 mg/mL FCDs were added to 240 μl bacterial suspensions to a final concentration of 200 μg/mL. The mixtures were rotated for 30 min at room temperature before centrifuging at 5,000 xg for 5 min. The supernatants were removed and the cell pellets re-suspended in 25 μl 2% paraformaldehyde (PFA). The cells were left to fix for 1 h at room temperature before transferring 5 μl into 15 μl ProLongTM Gold Antifade mountant (ThermoFisher). For cell membrane staining, FCDs were added to bacterial suspensions to a final concentration of 40 μg/mL and processed as above. Prior to the addition of PFA, the cell pellets were washed in PBS solution, resuspended in 25 μL of 100 μg/mL of FM™ 4-64FX (ThermoFisher) and shaken at 37 °C for 30 min. 10 μL of 4% PFA was then added and processed as above. The samples were mounted on glass slides with coverslips and left to set at room temperature for at least 12 h.

### HyVolution Confocal microscopy

Images were acquired on a Leica TCS SP8 system attached to a Leica DMi8 inverted microscope (Leica Microsystems) using a 100x HC PL APO CS2 oil immersion objective. The FCD-treated samples were excited using a 120mW 405 nm diode laser. Fluorescence of the FCDs were detected using a hybrid detector operating over an emission range of 480-550 nm and images acquired at 512 x 512 pixels.. FM™ 4-64FX (ThermoFisher)- treated samples were excited using a white light laser tuned to 561 nm and fluorescence detected over an emission range of 700-770 nm.

### Growth curves

5 mL overnight bacterial cultures were centrifuged at 2,900 xg for 10 min at room temperature and washed twice in PBS. Serial dilutions of FCDs in water were prepared in flat-bottom 96-well plates (Corning), inoculated with 10 μL of 1×10^8^ cfu/ml of washed cells and continuously shaken on a rotating platform with or without blue-LED exposure for up to 4 h. The samples were then further diluted 10-fold in Mueller Hinton broth 2 (Sigma-Aldrich) and placed in a plate reader (BMG Omega) for absorbance readings at 600 nm every 10 min for 16.5 h at 37°C.

### Viable cell counting

Sample preparation was as described for confocal imaging above. After 30 min exposure to FCDs, samples were irradiated with LED lights for 30, 60 or 90 min. The samples were then centrifuged at 5,000 xg for 10 min and the cell pellet resuspended in 1 mL of PBS. 25 μL of this suspension was taken and diluted in ten-fold serial dilutions up to 10^-7^. 100 μL of the last three dilutions were plated on to agar plates and incubated overnight. Cell colonies were then counted to determine viable cell counts.

### Temperature measurements

In a flat-bottom 96-well plate (Corning), 240 μL of bacterial suspensions were mixed with either 10 μL (800 μg/mL final concentration) or 2 μL (200 μg/mL) of 20 mg/mL FCDs. Control wells contained PBS only. Wells were either irradiated with LEDs or kept in the dark, temperature was recorded at regular 30 min intervals. A Thermocouple (type K from Fisher Scientific) was used to record temperature to 1 decimal place.

### Dihydroethidium ROS determination

192 μL of bacterial suspensions were added to wells of 2 flat-bottom 96 well plates. 8 μL of either FCDs (10 mg/mL) or sterile water was incubated for 1 hour. One plate was then irradiated with LED lights for 90 min whilst the other was kept in the dark. 10 μL dihydroethidium (DHE) (1 mg/mL) was then added to each well and fluorescence measured at λ_ex_ = 350 nm and λ_em_ = 460 nm (BMG PolarStar Omega plate reader).

### Scanning Electron Microscopy (SEM)

1 mL bacterial overnight cultures were centrifuged at 13,000 xg for 5 min and the pellets washed once in PBS. FCDs were added to yield a final concentration of 512 μg/mL and incubated for 30 min with rotation. After 30 min, the blue-LED board was placed in front of the rotator and samples not requiring LED-treatment were covered with foil. The samples were rotated for another 30 min before centrifuging at 8,000 xg for 5 min and washing twice in PBS. On the last wash, 50 μl of sample was removed, centrifuged and the pellet re-suspended in 50 μl of PBS. Samples were fixed on poly-L-lysine (0.1% w/v in water) treated glass coverslips in 2.5% glutaraldehyde overnight at 4°C. The samples were dehydrated in a series of ethanol solutions from 20, 50, 70, 90 and 100% and chemically dried with hexamethyldisilizane (HMDS) before being mounted on metal stubs and gold sputter coated (Emitech). The samples were imaged at magnifications of 20,000-120,000x on the FEI Quanta 200 FEG-SEM with a working distance of 10-13 mm, chamber pressure of <10^-5^ Pa in high vacuum mode and an accelerating voltage of 15-10 kV.

### Proteomics Sample Preparation and Analysis

#### Tandem Mass Tag (TMT) Labelling and High pH reversed-phase chromatography

Whole cell lysates were obtained from *E. coli* and *S. aureus* cultures grown in M9 minimal media in duplicate for 6 and 3 h respectively, with or without 512 μg/mL CD and continuous LED irradiation. Aliquots of 50 μg of each sample were digested with trypsin (2.5 μg trypsin per 100 μg protein; 37°C, overnight), labelled with Tandem Mass Tag (TMT) ten plex reagents according to the manufacturer’s protocol (Thermo Fisher Scientific) and the labelled samples pooled.

The pooled sample was evaporated to dryness, resuspended in 5% formic acid and then desalted using a SepPak cartridge according to the manufacturer’s instructions (Waters, Milford, Massachusetts, USA). Eluate from the SepPak cartridge was again evaporated to dryness and resuspended in buffer A (20 mM ammonium hydroxide, pH 10) prior to fractionation by high pH reversed-phase chromatography using an Ultimate 3000 liquid chromatography system (Thermo Scientific). In brief, the sample was loaded onto an XBridge BEH C18 Column (130Å, 3.5 μm, 2.1 mm X 150 mm, Waters, UK) in buffer A and peptides eluted with an increasing gradient of buffer B (20 mM Ammonium Hydroxide in acetonitrile, pH 10) from 0-95% over 60 minutes. The resulting fractions were evaporated to dryness and resuspended in 1% formic acid prior to analysis by nano-LC MSMS using an Orbitrap Fusion Lumos mass spectrometer (Thermo Scientific).

### Nano-LC Mass Spectrometry

High pH RP fractions were further fractionated using an Ultimate 3000 nano-LC system in line with an Orbitrap Fusion Lumos mass spectrometer (Thermo Scientific). In brief, peptides in 1% (vol/vol) formic acid were injected onto an Acclaim PepMap C18 nano-trap column (Thermo Scientific). After washing with 0.5% (vol/vol) acetonitrile 0.1% (vol/vol) formic acid peptides were resolved on a 250 mm × 75 μm Acclaim PepMap C18 reverse phase analytical column (Thermo Scientific) over a 150 min acetonitrile gradient, divided into 7 gradient segments (1-6% solvent B over 1 min., 6 - 15% B over 58 min., 15 - 32% B over 58 min., 32 - 40% B over 5 min., 40 - 90% B over 1 min., held at 90% B for 6 min and then reduced to 1% B over 1 min.) with a flow rate of 300 nl min^-1^. Solvent A was 0.1% formic acid and Solvent B was aqueous 80% acetonitrile in 0.1% formic acid. Peptides were ionized by nano-electrospray ionization at 2.0kV using a stainless steel emitter with an internal diameter of 30 μm (Thermo Scientific) and a capillary temperature of 275°C.

All spectra were acquired using an Orbitrap Fusion Lumos mass spectrometer controlled by Xcalibur 4.1 software (Thermo Scientific) and operated in data-dependent acquisition mode using an SPS-MS3 workflow. FTMS1 spectra were collected at a resolution of 120 000, with an automatic gain control (AGC) target of 200 000 and a max injection time of 50ms. Precursors were filtered with an intensity threshold of 5000, according to charge state (to include charge states 2-7) and with monoisotopic peak determination set to Peptide. Previously interrogated precursors were excluded using a dynamic window (60 s +/- 10 ppm). The MS2 precursors were isolated with a quadrupole isolation window of 0.7m/z. ITMS2 spectra were collected with an AGC target of 10 000, max injection time of 70ms and CID collision energy of 35%.

For FTMS3 analysis, the Orbitrap was operated at 50 000 resolution with an AGC target of 50 000 and a max injection time of 105 ms. Precursors were fragmented by high energy collision dissociation (HCD) at a normalised collision energy of 60% to ensure maximal TMT reporter ion yield. Synchronous Precursor Selection (SPS) was enabled to include up to 5 MS2 fragment ions in the FTMS3 scan.

### Data Analysis

The raw data files were processed and quantified using Proteome Discoverer software v2.1 (Thermo Scientific) and searched against the UniProt Human database (downloaded September 2018: 152927 entries) using the SEQUEST algorithm.^[22]^ Peptide precursor mass tolerance was set at 1ppm, and MS/MS tolerance was set at 0.Da. Search criteria included oxidation of methionine (+15.9949) as a variable modification and carbamidomethylation of cysteine (+57.0214) and the addition of the TMT mass tag (+229.163) to peptide N-termini and lysine as fixed modifications. Searches were performed with full tryptic digestion and a maximum of 2 missed cleavages were allowed. The reverse database search option was enabled and all data were filtered to satisfy a false discovery rate (FDR) of 5%.

Volcano plots were plotted to visualise statistical significance (p-value) versus fold change using R studio^[23]^. PantherDB was used for geneontology analysis^[24]^.

## Supporting information

ESI

## Acknowledgements

This research was supported by ERC-COG: 648239 (MCG), EPSRC EP/G036764/1 (JS) and EPSRC EP/S026215/1 (MCG, JS).We would like to thank the following members of the Wolfson Bioimaging Facility for their assistance: Chris Neal and Judith Mantell for the SEM work and Alan Leard for the confocal microscopy work and BrisSynBio, a BBSRC/EPSRC-funded Synthetic Biology Research Centre (grant number: L01386X) for funding the Hyvolution system. This publication has risen from discussion at COST Action GLYCONanoPROBES (CA18132), supported by COST (European Cooperation in Science and Technology).

## References

[1] R. Aminov, Biochem Pharmacol 2017, 133, 4–19.

[2]a) B. L. Roth, M. Poot, S. T. Yue, P. J. Millard, Appl Environ Microbiol 1997, 63, 2421–2431;

b) D. A. Drevets, A. M. Elliott, J Immunol Methods 1995, 187, 69–79;

c) C. Anaya, N. Church, J. P. Lewis, Proteomics 2007, 7, 215–219.

[3]a) H. Zhu, J. L. Fan, J. J. Du, X. J. Peng, Accounts Chem Res 2016, 49, 2115–2126;

b) Y. H. Chan, P. J. Wu, Part Part Syst Char 2015, 32, 11–28;

c) S. M. Janib, A. S. Moses, J. A. MacKay, Adv Drug Deliv Rev 2010, 62, 1052–1063;

d) J. Xie, S. Lee, X. Chen, Adv Drug Deliv Rev 2010, 62, 1064–1079;

e) M. A. Hahn, A. K. Singh, P. Sharma, S. C. Brown, B. M. Moudgil, Anal Bioanal Chem 2011, 399, 3–27.

[4] O. S. Wolfbeis, Chem Soc Rev 2015, 44, 4743–4768.

[5]a) Z. Zhu, Q. Li, P. Li, X. Xun, L. Zheng, D. Ning, M. Su, PLoS One 2019, 14, e0216230;

b) H. Zhu, J. Fan, J. Du, X. Peng, Acc Chem Res 2016, 49, 2115–2126;

c) S. Guo, Y. Sun, X. Geng, R. Yang, L. Xiao, L. Qu, Z. Li, J Mater Chem B 2020, 8, 736–742.

[6]a) W. Liu, C. Li, Y. Ren, X. Sun, W. Pan, Y. Li, J. Wang, W. Wang, J. Mat. Chem. B 2016, 4, 5772–5788;

b) H. Li, Z. Kang, Y. Liu, S.-T. Lee, J. Mat. Chem. 2012, 22, 24230–24253;

c) S. Y. Lim, W. Shen, Z. Gao, Chem. Soc. Rev. 2015, 44, 362–381;

d) S. N. Baker, G. A. Baker, Angew. Chem. Int. Ed. 2010, 49, 6726–6744;

e) X. Xu, R. Ray, Y. Gu, H. J. Ploehn, L. Gearheart, K. Raker, W. A. Scrivens, J. Am. Chem. Soc. 2004, 126, 12736–12737;

f) T. A. Swift, M. Duchi, S. A. Hill, D. Benito-Alifonso, R. L. Harniman, S. Sheikh, S. A. Davis, A. M. Seddon, H. M. Whitney, M. C. Galan, T. A. A. Oliver, Nanoscale 2018, 10, 13908–13912;

g) S. A. Hill, D. Benito-Alifonso, S. A. Davis, D. J. Morgan, M. Berry, M. C. Galan, Sci Rep 2018, 8;

h) S. Hill, M. C. Galan, Beilstein J Org Chem 2017, 13, 675–693;

i) S. A. Hill, D. Benito-Alifonso, D. J. Morgan, S. A. Davis, M. Berry, M. C. Galan, Nanoscale 2016, 8, 18630–18634.

[7]a) S. Y. Lim, W. Shen, Z. Q. Gao, Chem Soc Rev 2015, 44, 362–381;

b) P. Miao, K. Han, Y. G. Tang, B. D. Wang, T. Lin, W. B. Cheng, Nanoscale 2015, 7, 1586–1595;

c) S. N. Baker, G. A. Baker, Angew Chem Int Edit 2010, 49, 6726–6744;

d) N. L. Teradal, R. Jelinek, Adv Healthc Mater 2017, 6;

e) P. Miao, K. Han, Y. Tang, B. Wang, T. Lin, W. Cheng, Nanoscale 2015, 7, 1586–1595.

[8] D. Zhong, Y. Zhuo, Y. Feng, X. Yang, Biosens Bioelectron 2015, 74, 546–553.

[9]a) F. Lin, C. Li, Z. Chen, Front Microbiol 2018, 9, 2697;

b) C. I. Weng, H. T. Chang, C. H. Lin, Y. W. Shen, B. Unnikrishnan, Y. J. Li, C. C. Huang, Biosens Bioelectron 2015, 68, 1–6;

c) F. M. Lin, Y. W. Bao, F. G. Wu, C-J Carbon Res 2019, 5.

[10] H. Zhu, W. Lao, Q. Chen, Q. Zhang, H. Chen, Int J Clin Exp Med 2015, 8, 3651–3661.

[11]a) D. Benito-Alifonso, B. Richichi, V. Baldoneschi, M. Berry, M. Fragai, G. Salerno, M. C. Galan, C. Nativi, ACS Omega 2018, 3, 9822–9826;

b) D. Benito-Alifonso, S. Tremel, B. Hou, H. Lockyear, J. Mantell, D. J. Fermin, P. Verkade, M. Berry, M. C. Galan, Angew Chem Int Ed 2014, 53, 810–814;

c) R. Levy, U. Shaheen, Y. Cesbron, V. See, Nano Rev 2010, 1;

d) M. Lundqvist, J. Stigler, G. Elia, I. Lynch, T. Cedervall, K. A. Dawson, Proc Natl Acad Sci U S A 2008, 105, 14265–14270;

e) D. Bartczak, O. L. Muskens, S. Nitti, T. Sanchez-Elsner, T. M. Millar, A. G. Kanaras, Small 2012, 8, 122–130;

f) H. Yuan, A. M. Fales, T. Vo-Dinh, J Am Chem Soc 2012, 134, 11358–11361.

[12] S. A. Hill, S. Sheikh, Q. Y. Zhang, L. S. Ballesteros, A. Herman, S. A. Davis, D. J. Morgan, M. Berry, D. Benito-Alifonsoa, M. C. Galan, Nanoscale Advances 2019, 1, 2840–2846.

[13]a) A. Zhu, J. B. Huang, A. Clark, R. Romero, H. R. Petty, Carbohyd Res 2007, 342, 2745–2749;

b) A. Bhattacherjee, Y. Hrynets, M. Betti, J Agr Food Chem 2017, 65, 4642–4650.

[14] Y. Hrynets, A. Bhattacherjee, M. Ndagijimana, D. J. Hincapie Martinez, M. Betti, J Agric Food Chem 2016, 64, 3266–3275.

[15] Q. Sun, W. Margolin, J Bacteriol 1998, 180, 2050–2056.

[16] A. Bhattacherjee, Y. Hrynets, M. Betti, Food Chem 2019, 271, 354–361.

[17] Note that the internal bacteria temperature could not be determined and is likely to be higher than the temperature measured in the bacterial suspension.

[18] F. C. Fang, mBio 2011, 2.

[19]a) M. Kuroda, H. Kuroda, T. Oshima, F. Takeuchi, H. Mori, K. Hiramatsu, Mol Microbiol 2003, 49, 807–821;

b) S. Utaida, P. M. Dunman, D. Macapagal, E. Murphy, S. J. Projan, V. K. Singh, R. K. Jayaswal, B. J. Wilkinson, Microbiology (Reading) 2003, 149, 2719–2732;

c) F. McAleese, S. W. Wu, K. Sieradzki, P. Dunman, E. Murphy, S. Projan, A. Tomasz, J Bacteriol 2006, 188, 1120–1133;

d) A. Muthaiyan, J. A. Silverman, R. K. Jayaswal, B. J. Wilkinson, Antimicrob Agents Chemother 2008, 52, 980–990.

[20]a) M. J. Meziani, X. Dong, L. Zhu, L. P. Jones, G. E. LeCroy, F. Yang, S. Wang, P. Wang, Y. Zhao, L. Yang, R. A. Tripp, Y. P. Sun, ACS Appl Mater Interfaces 2016, 8, 10761–10766;

b) R. Jijie, A. Barras, J. Bouckaert, N. Dumitrascu, S. Szunerits, R. Boukherroub, Colloids Surf B Biointerf 2018, 170, 347–354.

[21]a) N. Sattarahmady, M. Rezaie-Yazdi, G. H. Tondro, N. Akbari, J Photochem Photobiol B 2017, 166, 323–332;

b) R. Knoblauch, C. D. Geddes, Materials 2020, 13.

[22] J. K. Eng, A. L. McCormack, J. R. Yates, J Am Soc Mass Spectrom 1994, 5, 976–989.

[23] RStudio Team (2020). RStudio: Integrated Development for R. RStudio, PBC, Boston, MA URL http://www.rstudio.com/

[24] PANTHER version 16: a revised family classification, tree-based classification tool, enhancer regions and extensive API. H. Mi, D. Ebert, A. Muruganujan, C. Mills, L.-P. Albou, T. Mushayamaha and P. D. Thomas. Nucl. Acids Res. (2020) doi: 10.1093/nar/gkaa1106s.

