## Supplementary material for "Green Fluorescent Carbon Dots as Targeting Probes for LED-Dependent Bacterial Killing": ESI

[1]School of Chemistry, Cantock's Close, BS8 1TS, University of Bristol; [2]School of Cellular & Molecular Medicine, Biomedical Sciences Building, BS8 1QU, University of Bristol; [3]Proteomics Facility, Biomedical Sciences Building, BS8 1TD, University of Bristol; [4]Wolfson Bioimaging Facility, Biomedical Sciences Building, BS8 1QU, University of Bristol; [5]FluoretiQ Limited, Unit DX, St Philips Central, Albert Road, BS2 0XJ, Bristol, UK

Contents: Page number:

| | |
|---|---|
| FCD labelling of different bacterial species | S2 |
| Comparison of core-FCDs and FCDs | S2 |
| Bacterial growth curves | S3 |
| Viable count at different LED irradiation durations | S4 |
| 2,5-deoxyfructosazine concentration estimation on FCDs | S5 |
| Photothermal effect | S5 |
| TMT proteomics | S6 |

#### FCD labelling of different bacterial species

Labelling of *E. coli, S. aureus, K. pneumoniae and P. aeruginosa* with green carbon dots at different concentrations of FCD

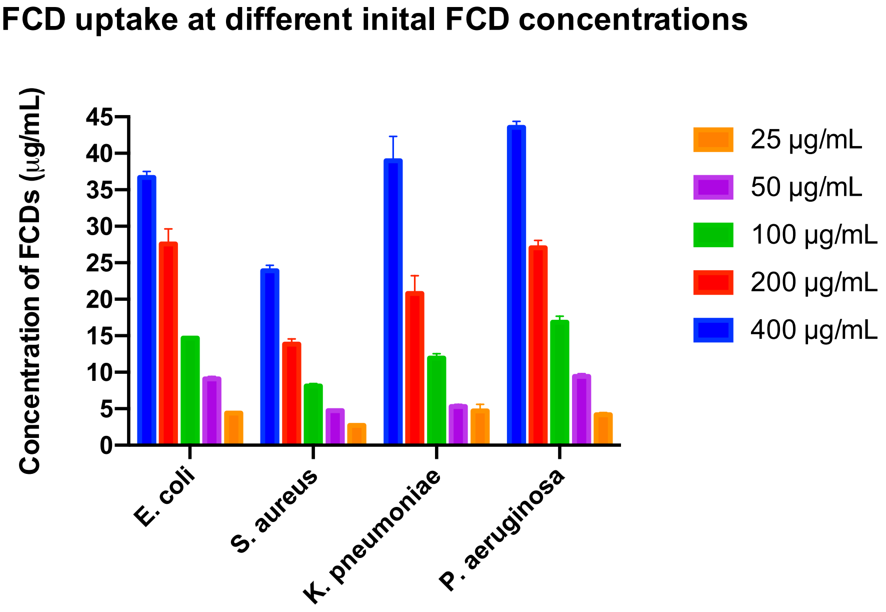


Figure S1. Labelling concentration of FCD with different species of bacteria at five different starting FCD concentrations

#### Comparison of core-FCDs and FCDs

Labelling of FCDs without further purification and dialysed FCDs, the majority of 2,5-deoxyfructosazine stripped from the surface. Fluorescence intensity calculated from confocal images.

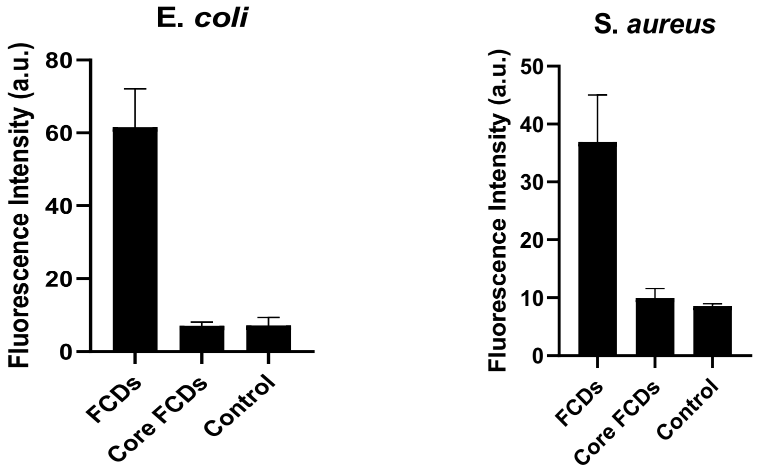


Figure S2. Fluorescent labelling of *E. coli* and *S. aureus* by FCDs and core FCDs (stripped of 2,5-deoxyfructosazine). Bacterial control represents autofluorescence level. Fluorescence intensities were averaged from confocal images and shown with SD error bars (n=5).

#### Bacterial growth curves

Bacterial suspensions treated to different concentrations of FCDs before irradiation with LEDs. Optical density ($OD_{600}$) measured every ten minutes over a 16-hour period to monitor growth.

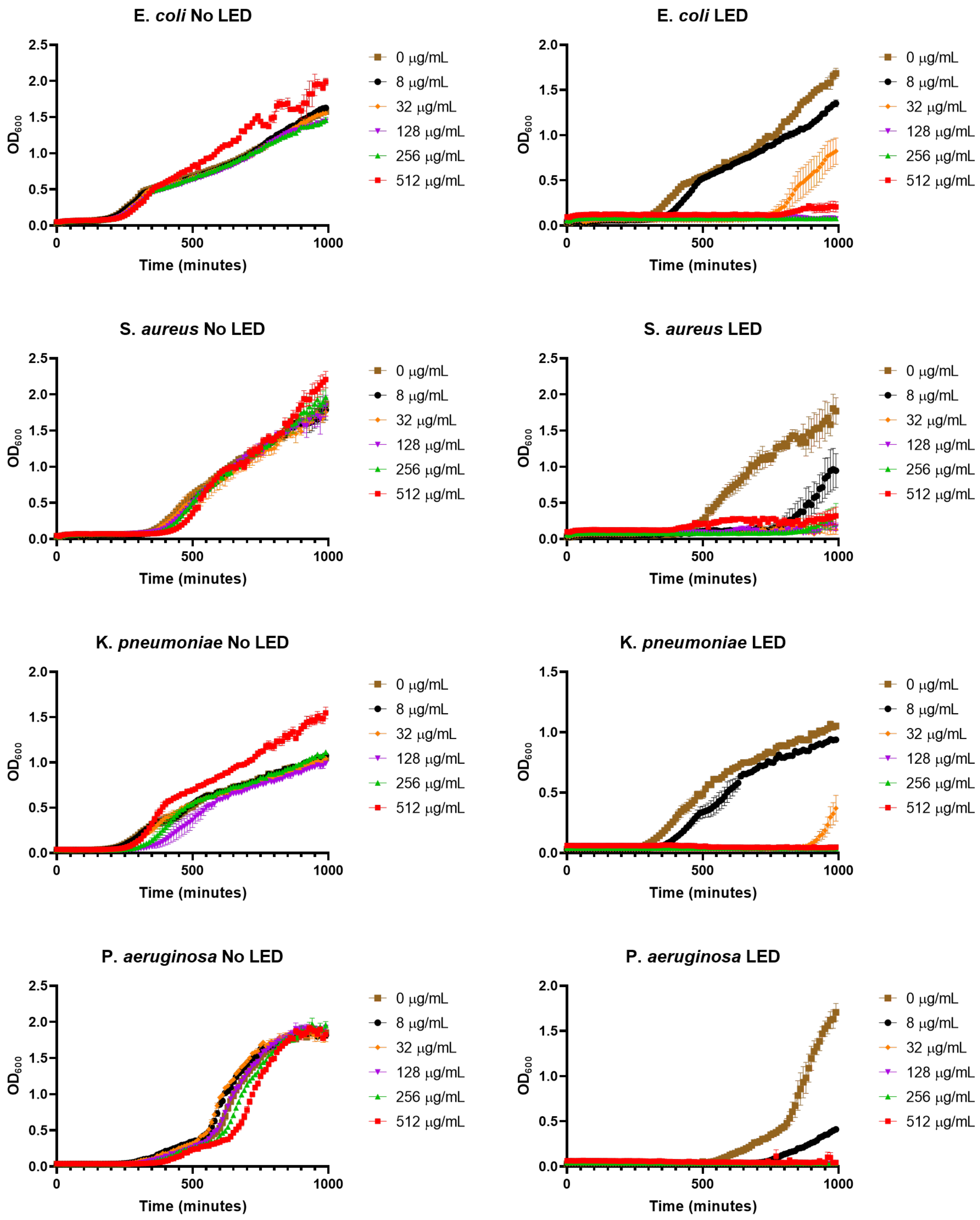


Figure S3. Growth curves of *E. coli, S. aureus, K. pneumoniae* and *P. aeruginosa* for the duration of 16 hours after treatment with varying concentrations of FCDs and 4 hours of LED irradiation

#### Viable count at different LED irradiation durations

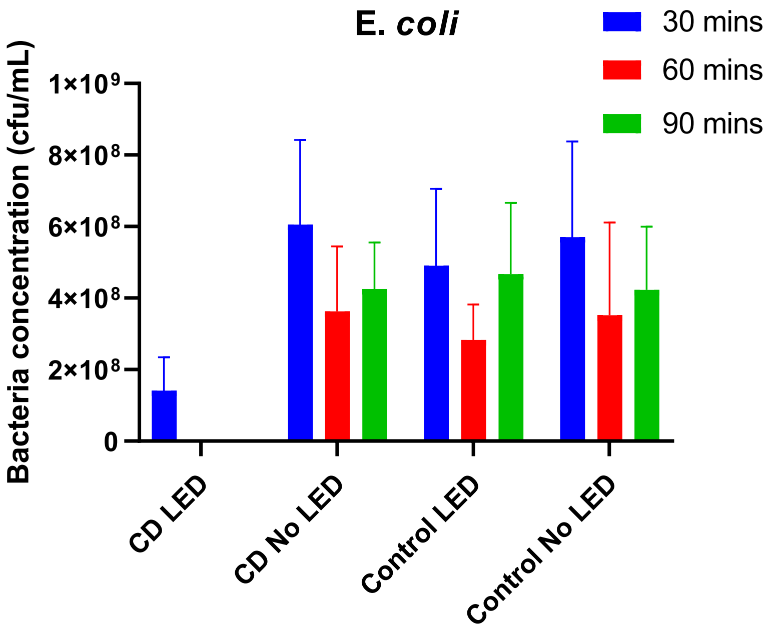


Figure S4. Viable count of *E. coli*. Bacteria inoculated onto agar plates after four different conditions; incubated with 200 µg/mL of FCDs for 30 mins and then irradiated with LEDs, incubated with FCDs for 30 mins with no LED irradiation, no FCD incubation with LED irradiation and no FCD incubation and no LED irradiation as control. Samples irradiation with LED for either 30, 60 and 90 mins. Samples were serially diluted to $10^6$ before inoculation. Counts were averaged and shown with SD error bars (n=6).

#### 2,5-deoxyfructosazine concentration estimation on FCDs

The distinctive pyrazine proton peaks can easily be identified within the FCD $^{1}$HNMR. The peak that is located between 8.70- 8.740 ppm was used as a reference peak to compare 2,5-deoxyfrutosazine (2,5-DOFR) concentration in the FCD samples.

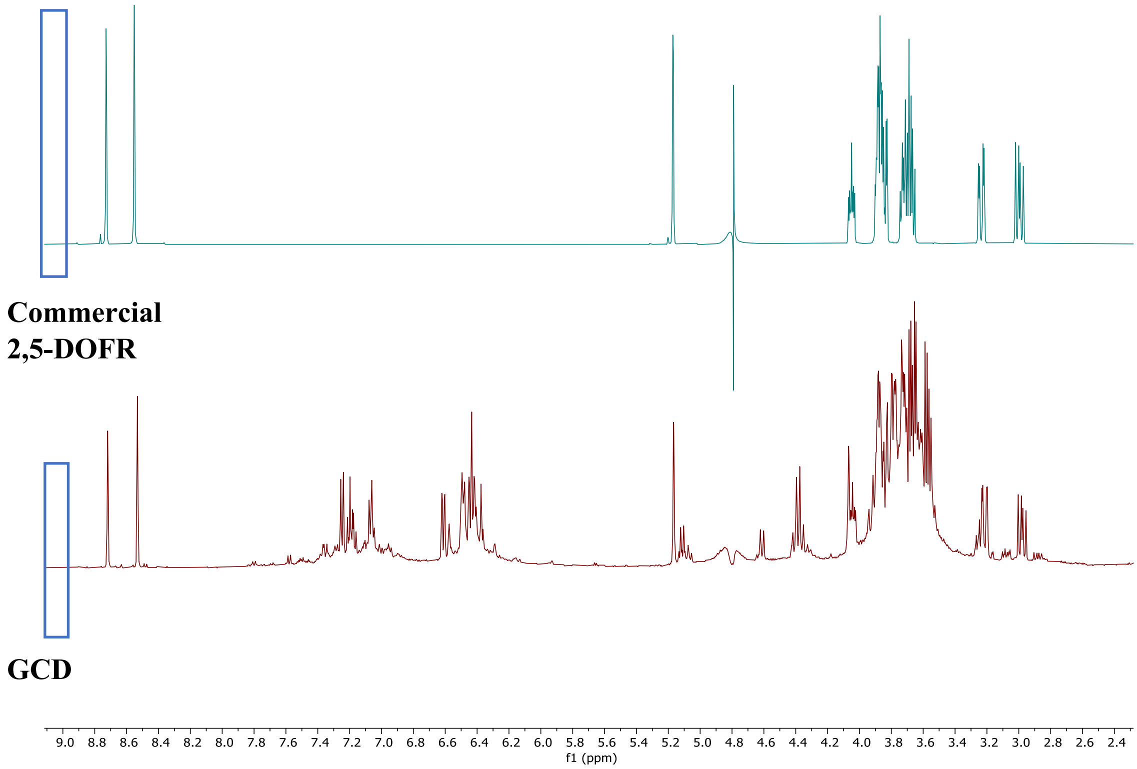


Figure S5. Stacked $^{1}$HNMR of commercially purchased 2,5-deoxyfructosazine and FCDs. Pyrazine peak highlighted

#### Photothermal effect

Temperature monitored after 30 minutes and 60 minutes LED irradiation times. Samples containing FCDs recorded higher temperatures. Two FCD concentrations tested, 200 µg/mL and 800 µg/mL.

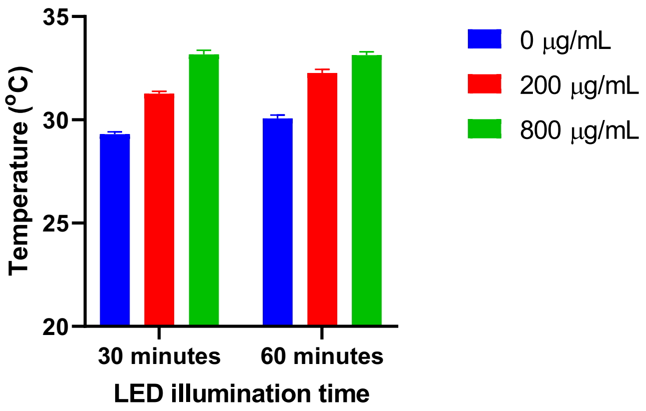


Figure S6. Temperature of FCD incubated bacterial suspensions after LED irradiation

### TMT proteomics

Table S1. *E. coli* proteins with ≥ 2-fold abundance changes compared to untreated control.

| UniProt Accession | Protein Description | Gene Name | LED/Control | | FCD/Control | | LED-FCD/Control | |
|---|---|---|---|---|---|---|---|---|
| | | | LogFC | T.Test | LogFC | T.Test | LogFC | T.Test |
| P77455 | Bifunctional protein | paaZ | 0.105 | 7.52E-01 | | | 4.863 | 3.82E-02 |
| P42619 | Inner membrane protein | yqjF | 0.049 | 8.72E-01 | 2.470 | 6.89E-02 | 4.533 | 1.94E-03 |
| Q6BF17 | D-galactonate dehydratase | dgoD | 0.414 | 3.13E-01 | 3.280 | 2.53E-03 | 4.167 | 1.01E-02 |
| P75975 | Prophage transcriptional regulatory protein | croE | 0.452 | 1.34E-01 | 1.046 | 2.33E-02 | 3.905 | 3.24E-02 |
| P75685 | Inner membrane protein | rclC | -0.020 | 9.65E-01 | -0.194 | 6.46E-01 | 3.714 | 2.72E-02 |
| P05824 | DNA repair protein | recN | 0.281 | 2.16E-01 | 1.002 | 9.67E-03 | 3.325 | 6.71E-03 |
| P0ABR1 | DNA damage-inducible protein | dinI | -0.155 | 4.14E-01 | 0.255 | 9.37E-02 | 3.144 | 2.67E-02 |
| P0ACY9 | Uncharacterized protein | yebG | -0.252 | 2.80E-01 | 0.257 | 1.93E-01 | 2.904 | 1.26E-02 |
| P19318 | Respiratory nitrate reductase 2 | narY | 0.195 | 1.66E-01 | -0.093 | 7.67E-01 | 2.563 | 4.30E-02 |
| P0AB46 | Uncharacterized protein | ymgD | 0.065 | 7.56E-01 | 1.826 | 1.89E-02 | 2.474 | 2.42E-02 |
| P76562 | tRNA(Met) cytidine acetyltransferase | tmcA | 0.209 | 2.58E-01 | 0.103 | 4.64E-01 | 2.206 | 4.37E-02 |
| P0AG71 | DNA recombination protein | rmuC | 0.215 | 6.98E-02 | 0.868 | 1.10E-01 | 2.148 | 2.50E-02 |
| P77754 | Periplasmic chaperone | spy | 0.042 | 6.43E-01 | 1.429 | 1.05E-02 | 2.079 | 8.28E-04 |
| P60932 | Undecaprenyl-diphosphatase | uppP | 0.497 | 2.04E-02 | -0.036 | 7.90E-01 | 2.031 | 5.03E-02 |
| P39377 | Isoaspartyl dipeptidase | iadA | 0.381 | 4.08E-01 | 1.173 | 1.53E-01 | 1.974 | 4.05E-02 |
| P76373 | UDP-glucose 6-dehydrogenase | ugd | -0.008 | 9.61E-01 | 0.046 | 7.67E-01 | 1.962 | 1.88E-02 |
| P0A698 | UvrABC system protein A | uvrA | 0.126 | 8.31E-02 | 1.066 | 2.44E-03 | 1.940 | 2.57E-02 |
| P07026 | Regulatory protein | sdiA | 0.419 | 2.98E-02 | 0.676 | 3.03E-01 | 1.938 | 9.29E-03 |
| P30744 | L-serine dehydratase 2 | sdaB | 0.757 | 1.90E-01 | 0.263 | 7.26E-01 | 1.912 | 3.46E-02 |
| P0AF03 | Molybdopterin adenylyltransferase | mog | 0.162 | 2.61E-02 | 0.099 | 1.88E-01 | 1.870 | 9.81E-02 |
| P0A7G6 | Protein RecA | recA | 0.015 | 9.34E-01 | 0.886 | 9.04E-02 | 1.865 | 1.70E-02 |
| P76559 | Uncharacterized protein YpfG | ypfG | 0.403 | 3.91E-01 | 1.378 | 1.65E-02 | 1.859 | 9.15E-03 |
| P76213 | Excinuclease | cho | -0.060 | 6.02E-01 | -0.452 | 6.93E-02 | 1.756 | 3.02E-02 |
| P77258 | N-ethylmaleimide reductase | nemA | -0.477 | 3.08E-02 | 0.709 | 5.94E-02 | 1.749 | 1.13E-03 |
| P0A809 | Holliday junction ATP-dependent DNA helicase | ruvA | -0.002 | 9.96E-01 | 0.451 | 3.24E-01 | 1.708 | 4.24E-02 |
| Q46807 | Carbamate kinase-like protein | yqeA | 0.297 | 3.90E-01 | 0.463 | 2.13E-01 | 1.655 | 4.46E-02 |
| P37353 | 2-succinylbenzoate--CoA ligase | menE | 0.199 | 3.35E-01 | 0.199 | 3.37E-01 | 1.649 | 1.82E-02 |
| P75992 | Probable two-component-system connector protein | ymgA | 0.908 | 1.48E-02 | -0.159 | 1.81E-01 | 1.635 | 1.61E-01 |
| Q7DFV3 | UPF0757 protein | ymgG | 0.042 | 8.74E-01 | 1.216 | 9.37E-02 | 1.633 | 4.15E-02 |
| P0A6L7 | Altronate oxidoreductase | uxaB | 0.141 | 2.51E-01 | 0.335 | 8.98E-02 | 1.632 | 3.62E-02 |
| P46482 | p-hydroxybenzoic acid efflux pump subunit | aaeA | -0.054 | 6.25E-01 | 0.399 | 7.08E-02 | 1.559 | 6.16E-03 |
| P33218 | Inner membrane protein | yebE | 0.380 | 4.75E-01 | 0.817 | 4.77E-02 | 1.558 | 2.09E-02 |
| P0A898 | Endoribonuclease | ybeY | 0.180 | 1.61E-01 | 0.211 | 1.64E-01 | 1.530 | 3.21E-03 |
| P0ABZ4 | 3-deoxy-D-manno-octulosonate 8-phosphate phosphatase | kdsC | 0.584 | 6.45E-02 | 0.192 | 4.89E-01 | 1.488 | 7.55E-03 |
| P69739 | Hydrogenase-1 small chain | hyaA | 0.182 | 4.58E-01 | 0.353 | 2.79E-01 | 1.474 | 2.65E-02 |
| P71298 | Prophage integrase | intF | -0.300 | 1.82E-01 | -0.145 | 5.03E-01 | 1.434 | 2.29E-02 |
| P77596 | D-xylonate dehydratase | yagF | -0.082 | 7.71E-01 | -0.098 | 6.31E-01 | 1.406 | 2.71E-02 |
| P09148 | Galactose-1-phosphate uridylyltransferase | galT | 0.134 | 2.62E-02 | 0.662 | 2.96E-02 | 1.396 | 1.83E-02 |
| P0AGG2 | Acyl-CoA thioesterase 2 | tesB | 0.107 | 7.57E-02 | 0.695 | 3.50E-03 | 1.384 | 7.46E-02 |
| P0AFU2 | Uncharacterized transporter | yfbS | 0.095 | 6.24E-01 | 0.091 | 1.58E-01 | 1.380 | 4.61E-02 |
| P27859 | 3'-5' ssDNA/RNA exonuclease | tatD | 0.090 | 3.08E-01 | 0.203 | 1.34E-01 | 1.374 | 1.97E-02 |
| P21151 | 3-ketoacyl-CoA thiolase | fadA | 0.039 | 5.06E-02 | 1.349 | 4.18E-02 | 1.364 | 1.82E-02 |
| P13029 | Catalase-peroxidase | katG | 0.107 | 4.71E-01 | 0.949 | 2.06E-02 | 1.362 | 1.17E-02 |
| P13033 | Anaerobic glycerol-3-phosphate dehydrogenase subunit B | glpB | 0.291 | 1.37E-02 | 0.508 | 1.19E-01 | 1.361 | 1.69E-01 |

| | | | | | | | | |
|---|---|---|---|---|---|---|---|---|
| P37351 | Ribose-5-phosphate isomerase B | rpiB | 0.014 | 5.22E-01 | 1.453 | 3.43E-04 | 1.350 | 6.53E-02 |
| P42624 | Pirin-like protein YhaK | yhaK | 0.117 | 1.56E-01 | 0.192 | 8.22E-02 | 1.347 | 1.72E-03 |
| Q46814 | Probable hypoxanthine oxidase | xdhD | 0.224 | 4.53E-02 | 0.509 | 1.51E-02 | 1.344 | 4.04E-03 |
| P41407 | FMN-dependent NADH-azoreductase | azoR | -0.070 | 6.08E-01 | 0.727 | 7.41E-02 | 1.343 | 1.54E-02 |
| P0A8F8 | UvrABC system protein B | uvrB | 0.158 | 4.22E-02 | 0.444 | 1.61E-02 | 1.341 | 1.16E-02 |
| P0C0V0 | Periplasmic serine endoprotease | degP | 0.129 | 2.13E-01 | 1.531 | 3.50E-04 | 1.324 | 3.11E-02 |
| P0ACR2 | Uncharacterized HTH-type transcriptional regulator | ydhB | 0.078 | 4.78E-01 | 0.418 | 5.10E-02 | 1.317 | 9.87E-03 |
| P21177 | Fatty acid oxidation complex subunit alpha | fadB | -0.001 | 9.88E-01 | 1.549 | 8.94E-03 | 1.314 | 6.33E-03 |
| P0ABK9 | Cytochrome c-552 | nrfA | 0.266 | 1.40E-03 | 0.236 | 3.29E-01 | 1.296 | 6.66E-02 |
| P76399 | Multidrug resistance protein | mdtC | -0.003 | 9.52E-01 | 0.732 | 1.41E-02 | 1.296 | 1.05E-01 |
| P0AAE2 | Proline-specific permease | proY | -0.074 | 7.90E-01 | -0.157 | 6.44E-01 | 1.288 | 4.96E-02 |
| P28904 | Trehalose-6-phosphate hydrolase | treC | 0.130 | 4.63E-01 | 1.147 | 6.39E-02 | 1.267 | 1.68E-02 |
| P0ADV9 | Lipopolysaccharide export system protein | lptC | -0.077 | 6.32E-01 | -0.108 | 5.28E-01 | 1.256 | 3.24E-02 |
| P27896 | Nitrate/nitrite sensor protein | narQ | 0.146 | 1.46E-01 | 0.442 | 1.34E-02 | 1.255 | 1.27E-01 |
| Q47537 | Taurine-binding periplasmic protein | tauA | -0.096 | 2.78E-01 | 1.113 | 1.65E-03 | 1.241 | 9.61E-02 |
| P17411 | 6-phospho-beta-glucosidase | chbF | -0.020 | 8.62E-01 | 0.621 | 6.57E-02 | 1.238 | 2.41E-02 |
| P16384 | tRNA dimethylallyltransferase | miaA | -0.075 | 3.94E-01 | 0.591 | 1.83E-02 | 1.220 | 1.92E-02 |
| P0ACA3 | Stringent starvation protein A | sspA | 0.021 | 8.85E-01 | 0.447 | 1.45E-01 | 1.195 | 1.62E-02 |
| P28629 | Biodegradative arginine decarboxylase | adiA | 0.096 | 6.50E-01 | 0.553 | 1.46E-01 | 1.179 | 3.28E-02 |
| P24197 | 4,5-DOPA dioxygenase extradiol | ygiD | 0.200 | 4.05E-01 | 0.850 | 1.12E-02 | 1.175 | 6.26E-02 |
| P60716 | Lipoyl synthase | lipA | 0.446 | 2.42E-01 | 0.225 | 4.11E-01 | 1.165 | 4.51E-02 |
| P0AGJ7 | tRNA (cytidine(34)-2'-O)-methyltransferase | trmL | -0.077 | 5.12E-01 | 0.228 | 4.54E-01 | 1.155 | 2.64E-02 |
| P37610 | Alpha-ketoglutarate-dependent taurine dioxygenase | tauD | -0.217 | 1.83E-01 | 1.151 | 1.69E-02 | 1.153 | 3.41E-02 |
| P15028 | Fe(3+) dicitrate-binding periplasmic protein | fecB | -0.018 | 8.69E-01 | 0.821 | 2.64E-02 | 1.147 | 1.02E-01 |
| P37648 | Protein YhjJ | yhjJ | -0.085 | 2.73E-01 | 0.998 | 3.67E-03 | 1.140 | 7.18E-03 |
| P0AD59 | Inhibitor of vertebrate lysozyme | ivy | 0.076 | 4.25E-01 | 0.579 | 1.22E-01 | 1.121 | 7.23E-05 |
| P0ADP7 | Ubiquinone biosynthesis protein | ubiJ | 0.157 | 2.91E-01 | 0.323 | 1.13E-01 | 1.103 | 2.23E-02 |
| P0A6C5 | Amino-acid acetyltransferase | argA | 0.042 | 1.26E-01 | 0.665 | 6.27E-04 | 1.096 | 1.71E-01 |
| P36672 | PTS system trehalose-specific EIIBC component | treB | 0.135 | 8.47E-02 | 0.858 | 3.24E-01 | 1.089 | 1.37E-02 |
| P17444 | Oxygen-dependent choline dehydrogenase | betA | 0.032 | 8.27E-01 | 0.642 | 8.61E-02 | 1.087 | 2.17E-02 |
| P0A9X1 | Zinc import ATP-binding protein | znuC | 0.006 | 9.65E-01 | -0.803 | 3.88E-02 | 1.085 | 2.10E-01 |
| P0AE60 | Cell division activator | cedA | 0.096 | 3.08E-01 | 0.186 | 1.74E-01 | 1.062 | 4.77E-03 |
| P31828 | Probable zinc protease | pqqL | -0.211 | 4.61E-03 | 1.090 | 8.67E-04 | 1.049 | 1.57E-03 |
| P25396 | Tellurite resistance protein | tehA | 0.293 | 3.82E-02 | 0.270 | 2.48E-01 | 1.047 | 9.06E-04 |
| P18392 | Sensor protein | rstB | 0.046 | 7.67E-01 | 0.564 | 1.08E-01 | 1.042 | 3.08E-02 |
| Q46871 | NADPH-dependent ferric-chelate reductase | yqjH | -0.036 | 8.55E-01 | 0.546 | 1.15E-01 | 1.038 | 4.10E-02 |
| P0A7N1 | 50S ribosomal protein L31 type B | ykgM | 0.230 | 1.33E-01 | 1.225 | 4.95E-02 | 1.029 | 1.02E-01 |
| P30015 | Probable ATP-dependent helicase | lhr | 0.063 | 6.90E-01 | 0.253 | 1.97E-01 | 1.022 | 2.70E-02 |
| P76537 | Uncharacterized protein YfeY | yfeY | 0.045 | 5.04E-01 | 1.193 | 1.45E-02 | 1.009 | 4.89E-02 |
| P13035 | Aerobic glycerol-3-phosphate dehydrogenase | glpD | 0.303 | 5.38E-02 | 0.503 | 5.74E-02 | 1.009 | 2.78E-02 |
| P0A9P6 | ATP-dependent RNA helicase | deaD | 1.153 | 3.41E-02 | 0.073 | 5.43E-01 | 0.900 | 2.04E-02 |
| P0DP90 | Acetolactate synthase isozyme 2 large subunit | ilvG | 1.551 | 7.82E-03 | 1.618 | 1.74E-02 | 0.791 | 7.12E-02 |
| P33940 | Malate:quinone oxidoreductase | mqo | -0.316 | 9.66E-02 | 1.517 | 1.80E-02 | 0.788 | 1.49E-02 |
| P69506 | Iron-sulfur cluster repair protein | ytfE | 0.343 | 5.29E-02 | 1.012 | 4.30E-01 | 0.678 | 4.74E-03 |
| P0ADT8 | Uncharacterized protein YgiM | ygiM | -0.005 | 9.72E-01 | 1.187 | 3.62E-02 | 0.436 | 1.13E-01 |
| P64426 | UPF0748 lipoprotein | yddW | -0.499 | 1.15E-02 | -1.264 | 2.27E-03 | 0.428 | 2.90E-01 |
| P14294 | DNA topoisomerase 3 | topB | -0.633 | 6.02E-02 | -1.217 | 3.03E-02 | 0.194 | 6.33E-01 |
| P0ADX7 | Uncharacterized protein YhhA | yhhA | 0.107 | 3.67E-01 | -1.577 | 2.32E-02 | 0.079 | 8.59E-01 |
| P0A8C8 | Putative membrane protein insertion efficiency factor | yidD | -0.479 | 8.12E-02 | -1.203 | 7.96E-03 | -0.301 | 9.73E-02 |

| | | | | | | | | |
|---|---|---|---|---|---|---|---|---|
| P0AAY6 | Uncharacterized protein YbjN | ybjN | -0.990 | 4.63E-02 | -1.342 | 3.12E-02 | -0.327 | 2.57E-01 |
| P77611 | Ion-translocating oxidoreductase complex subunit C | rsxC | -0.951 | 1.63E-01 | -1.214 | 1.29E-02 | -0.370 | 2.98E-02 |
| P0AF90 | Regulator of ribonuclease activity B | rraB | -0.943 | 1.60E-02 | -1.051 | 6.35E-02 | -0.408 | 9.44E-02 |
| P0AEX7 | High-affinity branched-chain amino acid transport system permease protein | livH | -1.083 | 4.39E-04 | -1.491 | 2.32E-02 | -0.459 | 2.84E-02 |
| P10384 | Long-chain fatty acid transport protein | fadL | 0.469 | 1.52E-02 | -1.147 | 3.11E-01 | -0.569 | 4.68E-02 |
| P11289 | Uncharacterized protein YfiL | yfiL | 0.078 | 6.73E-01 | -1.932 | 4.32E-02 | -0.724 | 9.04E-02 |
| P0ACU5 | HTH-type transcriptional repressor | fabR | -0.215 | 5.27E-01 | -1.135 | 7.64E-03 | -0.827 | 1.07E-02 |
| P0C8J8 | D-tagatose-1,6-bisphosphate aldolase subunit | gatZ | -0.471 | 6.15E-01 | -2.164 | 1.48E-02 | -0.858 | 7.63E-02 |
| P0A910 | Outer membrane protein A | ompA | 0.088 | 2.31E-01 | -1.346 | 2.06E-01 | -0.882 | 1.75E-02 |
| P0AEX9 | Maltose/maltodextrin-binding periplasmic protein | malE | 0.412 | 9.04E-02 | -1.197 | 1.04E-02 | -0.884 | 1.54E-01 |
| P68679 | 30S ribosomal protein S21 | rpsU | -0.185 | 1.80E-01 | -1.132 | 1.07E-01 | -0.957 | 1.00E-02 |
| P0AFC7 | NADH-quinone oxidoreductase subunit B | nuoB | 0.178 | 1.57E-01 | -0.192 | 1.84E-01 | -1.004 | 2.62E-02 |
| P06610 | Thioredoxin/glutathione peroxidase | btuE | -0.034 | 7.41E-01 | -0.209 | 1.77E-01 | -1.009 | 1.12E-02 |
| P0AAA1 | Inner membrane protein | yagU | -0.241 | 2.40E-02 | -0.767 | 1.73E-01 | -1.012 | 4.85E-03 |
| P0A9L8 | Pyrroline-5-carboxylate reductase | proC | 0.084 | 5.56E-01 | -0.271 | 3.83E-02 | -1.013 | 8.52E-03 |
| P0AGH1 | Inner membrane transport permease | yhhJ | 0.332 | 6.74E-02 | 0.069 | 4.43E-01 | -1.014 | 1.33E-02 |
| P76270 | Free methionine-R-sulfoxide reductase | msrC | -0.003 | 9.85E-01 | -0.444 | 1.26E-01 | -1.018 | 4.03E-02 |
| P0A9F9 | HTH-type transcriptional regulator | metR | -1.188 | 3.25E-01 | -2.116 | 4.53E-02 | -1.028 | 7.46E-02 |
| P37349 | PEP-dependent dihydroxyacetone kinase, phosphoryl donor subunit | dhaM | 0.014 | 9.20E-01 | -0.178 | 3.46E-01 | -1.028 | 4.60E-03 |
| P32173 | Molybdenum cofactor guanylyltransferase | mobA | 0.084 | 5.39E-01 | -0.007 | 9.45E-01 | -1.029 | 2.32E-02 |
| P30131 | Carbamoyltransferase | hypF | -0.006 | 9.46E-01 | -0.207 | 1.32E-01 | -1.038 | 3.82E-02 |
| P29013 | Uncharacterized protein YcgB | ycgB | 0.026 | 8.68E-01 | -0.138 | 4.73E-01 | -1.047 | 1.87E-02 |
| P06996 | Outer membrane protein C | ompC | -0.355 | 2.73E-01 | -1.073 | 7.12E-02 | -1.050 | 3.96E-02 |
| P46144 | Uncharacterized protein YedJ | yedJ | 0.163 | 1.21E-01 | -0.095 | 2.74E-01 | -1.053 | 3.85E-03 |
| P0A9D2 | Glutathione S-transferase | gstA | 0.049 | 6.40E-01 | -0.041 | 7.10E-01 | -1.055 | 1.08E-02 |
| P0A843 | Sec-independent protein translocase protein | tatE | -1.212 | 2.08E-02 | -0.982 | 1.66E-02 | -1.056 | 8.92E-03 |
| P0AFE4 | NADH-quinone oxidoreductase subunit K | nuoK | 0.471 | 1.71E-01 | -0.122 | 6.92E-01 | -1.057 | 4.41E-02 |
| P23857 | Thiosulfate sulfurtransferase | pspE | -0.426 | 8.56E-02 | -0.604 | 5.00E-02 | -1.059 | 5.71E-02 |
| P0AF93 | 2-iminobutanoate/2-iminopropanoate deaminase | ridA | -0.034 | 7.36E-01 | -0.180 | 2.43E-01 | -1.080 | 9.42E-03 |
| P33018 | S-formylglutathione hydrolase | yeiG | 0.157 | 2.18E-01 | -0.635 | 1.75E-02 | -1.090 | 1.20E-02 |
| P03024 | HTH-type transcriptional regulator | galR | 0.091 | 5.16E-01 | -0.236 | 2.13E-01 | -1.093 | 1.60E-02 |
| P09832 | Glutamate synthase [NADPH] small chain | gltD | 0.011 | 8.07E-01 | -0.710 | 5.08E-03 | -1.094 | 5.05E-02 |
| P64590 | Inner membrane protein | yhaH | 0.564 | 6.19E-02 | -0.297 | 6.05E-01 | -1.099 | 2.91E-02 |
| P0AES9 | Acid stress chaperone | hdeA | -0.010 | 9.29E-01 | -0.334 | 3.79E-01 | -1.099 | 1.40E-02 |
| P0C0S1 | Small-conductance mechanosensitive channel | mscS | 0.242 | 1.93E-01 | -0.518 | 9.07E-02 | -1.103 | 4.19E-02 |
| P0A814 | Crossover junction endodeoxyribonuclease | ruvC | -0.206 | 3.12E-01 | 0.026 | 8.93E-01 | -1.111 | 1.80E-02 |
| P13482 | Periplasmic trehalase | treA | 0.206 | 2.38E-01 | -0.195 | 2.22E-01 | -1.111 | 1.91E-02 |
| P0AF59 | Uncharacterized protein YjdI | yjdI | 0.274 | 1.31E-01 | -0.613 | 7.12E-02 | -1.113 | 1.25E-02 |
| P77581 | Succinylornithine transaminase | astC | 0.116 | 4.67E-01 | -0.476 | 1.10E-01 | -1.124 | 3.68E-02 |
| P64540 | Uncharacterized protein YfcL | yfcL | -0.510 | 1.00E-01 | -0.691 | 1.32E-01 | -1.126 | 3.93E-02 |
| P0AEJ2 | Isochorismate synthase | entC | -1.465 | 3.62E-02 | -1.556 | 2.58E-02 | -1.127 | 2.88E-02 |
| P07014 | Succinate dehydrogenase iron-sulfur subunit | sdhB | 0.287 | 9.58E-03 | -0.872 | 1.27E-02 | -1.127 | 5.50E-02 |
| P76507 | Uncharacterized protein YfdI | yfdI | -0.130 | 2.57E-01 | -0.465 | 4.81E-01 | -1.131 | 2.68E-03 |
| P0ACK5 | Deoxyribose operon repressor | deoR | -0.100 | 4.64E-01 | -0.206 | 2.19E-01 | -1.139 | 2.09E-02 |
| P0AB14 | Uncharacterized protein YccJ | yccJ | 0.173 | 1.39E-01 | -0.711 | 4.03E-02 | -1.145 | 2.57E-02 |
| P76142 | Autoinducer 2-binding protein | lsrB | -0.372 | 1.76E-01 | -0.614 | 1.73E-01 | -1.151 | 3.33E-02 |
| P00894 | Acetolactate synthase isozyme 3 small subunit | ilvH | -0.173 | 1.78E-01 | -0.841 | 1.02E-02 | -1.159 | 9.24E-02 |
| P06993 | HTH-type transcriptional regulator | malT | -0.636 | 2.65E-01 | -1.804 | 3.49E-03 | -1.167 | 1.48E-02 |

| | | | | | | | | |
|---|---|---|---|---|---|---|---|---|
| P15067 | Glycogen debranching enzyme | glgX | -0.199 | 7.20E-02 | -0.558 | 4.37E-02 | -1.187 | 7.64E-03 |
| P0A7Q6 | 50S ribosomal protein L36 | rpmJ | -0.254 | 9.16E-02 | -0.574 | 1.71E-01 | -1.200 | 4.71E-02 |
| P0AE37 | Arginine N-succinyltransferase | astA | 0.267 | 2.68E-01 | -0.288 | 2.47E-01 | -1.207 | 1.98E-02 |
| P21420 | Putative outer membrane porin protein | nmpC | -0.109 | 5.05E-01 | -1.305 | 1.92E-03 | -1.209 | 5.03E-02 |
| P77214 | Cation efflux system protein | cusF | 1.581 | 1.19E-01 | -2.150 | 2.73E-02 | -1.214 | 5.86E-02 |
| P08550 | Colicin V production protein | cvpA | -0.008 | 9.22E-01 | 0.167 | 2.67E-01 | -1.216 | 1.65E-02 |
| P63340 | Inner membrane transport protein | yqeG | 0.089 | 4.45E-01 | -0.938 | 1.23E-02 | -1.217 | 2.87E-02 |
| P0C0T5 | Penicillin-insensitive murein endopeptidase | mepA | -0.174 | 3.99E-01 | -0.489 | 1.05E-01 | -1.221 | 3.28E-02 |
| P23847 | Periplasmic dipeptide transport protein | dppA | 0.123 | 4.27E-02 | -1.094 | 5.30E-02 | -1.233 | 1.76E-02 |
| P39180 | Antigen 43 | flu | -0.190 | 3.00E-02 | -1.340 | 1.78E-01 | -1.235 | 1.12E-02 |
| P00893 | Acetolactate synthase isozyme 3 large subunit | ilvI | -0.274 | 3.52E-02 | -1.032 | 1.37E-02 | -1.248 | 1.70E-03 |
| P0A734 | Cell division topological specificity factor | minE | -0.120 | 1.51E-01 | -0.114 | 5.07E-01 | -1.251 | 5.68E-03 |
| P0AED0 | Universal stress protein A | uspA | 0.003 | 8.86E-01 | -0.170 | 3.90E-01 | -1.256 | 4.28E-02 |
| P31135 | Putrescine transport system permease protein | potH | 0.309 | 2.05E-01 | 0.531 | 9.10E-02 | -1.257 | 8.64E-03 |
| P0A9H7 | Cyclopropane-fatty-acyl-phospholipid synthase | cfa | -0.532 | 5.59E-03 | -0.447 | 7.64E-02 | -1.260 | 3.64E-02 |
| P77735 | 1-deoxyxylulose-5-phosphate synthase | yajO | -0.009 | 9.33E-01 | -0.596 | 4.07E-02 | -1.271 | 7.35E-03 |
| P0AEK0 | Exodeoxyribonuclease 10 | exoX | -0.204 | 1.99E-01 | -0.690 | 2.70E-01 | -1.272 | 2.67E-02 |
| P02931 | Outer membrane protein F | ompF | -0.108 | 5.88E-01 | -1.646 | 4.18E-02 | -1.284 | 1.82E-02 |
| P75839 | UPF0702 transmembrane protein | ycaP | 0.154 | 2.50E-01 | -0.325 | 1.11E-01 | -1.298 | 3.20E-03 |
| P62601 | Cytoplasmic trehalase | treF | 0.009 | 9.01E-01 | -0.365 | 1.85E-01 | -1.314 | 2.36E-03 |
| P0A769 | Divalent metal cation transporter | mntH | -0.470 | 4.90E-02 | -0.129 | 8.39E-01 | -1.314 | 2.55E-02 |
| P14176 | Glycine betaine/proline betaine transport system permease protein | proW | -0.254 | 1.60E-01 | -0.556 | 2.59E-01 | -1.323 | 3.20E-02 |
| P0AFW2 | Ribosome modulation factor | rmf | -0.210 | 6.87E-01 | -0.430 | 2.40E-01 | -1.336 | 3.88E-02 |
| P0ABQ4 | Dihydrofolate reductase | folA | 0.207 | 2.71E-01 | -0.826 | 1.78E-01 | -1.337 | 3.30E-02 |
| P0A786 | Aspartate carbamoyltransferase catalytic subunit | pyrB | -0.139 | 3.91E-01 | -1.098 | 5.98E-02 | -1.345 | 4.08E-02 |
| P00490 | Maltodextrin phosphorylase | malP | 0.322 | 9.71E-03 | -0.770 | 3.27E-03 | -1.373 | 2.12E-02 |
| P22729 | High-affinity branched-chain amino acid transport system permease protein | livM | -0.415 | 2.36E-01 | -1.104 | 5.94E-02 | -1.373 | 3.05E-02 |
| P0AFC3 | NADH-quinone oxidoreductase subunit A | nuoA | 0.217 | 4.03E-01 | -0.149 | 5.29E-01 | -1.383 | 4.19E-02 |
| P0ACX5 | Fumarase D | fumD | 0.324 | 4.88E-02 | -0.561 | 5.13E-02 | -1.386 | 8.18E-02 |
| P03841 | Maltose operon periplasmic protein | malM | 0.370 | 6.22E-02 | -1.682 | 8.80E-03 | -1.391 | 1.57E-03 |
| P0AD24 | UPF0352 protein | yejL | -0.506 | 1.20E-01 | -1.134 | 3.12E-02 | -1.399 | 5.19E-02 |
| P0A7F3 | Aspartate carbamoyltransferase regulatory chain | pyrI | -0.106 | 3.76E-01 | -1.168 | 1.99E-02 | -1.412 | 4.96E-03 |
| P0AFD4 | NADH-quinone oxidoreductase subunit H | nuoH | 0.442 | 1.32E-02 | -0.310 | 4.05E-01 | -1.427 | 1.77E-02 |
| P31475 | Uncharacterized HTH-type transcriptional regulator | yieP | 0.143 | 2.45E-01 | -0.540 | 2.91E-02 | -1.433 | 1.29E-02 |
| P0AFM4 | Phosphate starvation-inducible protein | psiF | 0.185 | 4.65E-01 | -0.664 | 9.92E-02 | -1.444 | 2.52E-02 |
| P0ADF8 | Acetolactate synthase isozyme 1 small subunit | ilvN | -0.563 | 1.29E-01 | -0.180 | 3.73E-01 | -1.449 | 1.88E-02 |
| P0AFU8 | Riboflavin synthase | ribC | -0.035 | 4.83E-01 | -0.250 | 3.10E-02 | -1.479 | 8.36E-02 |
| P0AET2 | Acid stress chaperone | hdeB | -0.107 | 3.80E-01 | -0.557 | 2.46E-01 | -1.494 | 2.07E-03 |
| P37327 | Inner membrane protein | yfdC | 0.048 | 6.17E-01 | -0.607 | 3.40E-02 | -1.517 | 3.21E-02 |
| P05793 | Ketol-acid reductoisomerase (NADP(+)) | ilvC | -1.643 | 1.13E-02 | -1.475 | 5.54E-02 | -1.539 | 4.02E-02 |
| P05804 | Beta-glucuronidase | uidA | -0.009 | 9.12E-01 | -0.139 | 1.38E-01 | -1.543 | 1.96E-03 |
| P04816 | Leucine-specific-binding protein | livK | -1.014 | 6.20E-03 | -1.423 | 1.23E-03 | -1.574 | 5.50E-02 |
| P45795 | Uncharacterized protein YrdB | yrdB | 0.129 | 6.83E-01 | -0.373 | 3.32E-01 | -1.575 | 3.85E-02 |
| P18956 | Glutathione hydrolase proenzyme | ggt | -0.044 | 8.12E-01 | -0.616 | 9.54E-02 | -1.600 | 2.70E-02 |
| P24188 | UPF0176 protein | yceA | -0.544 | 4.13E-01 | -1.448 | 1.96E-02 | -1.605 | 1.70E-02 |
| P0C960 | Endo-type membrane-bound lytic murein transglycosylase A | emtA | -0.329 | 3.50E-01 | -0.302 | 4.29E-01 | -1.618 | 4.44E-02 |
| P0A8L7 | UPF0263 protein | yciU | -0.293 | 2.34E-01 | -0.897 | 8.49E-02 | -1.620 | 4.97E-02 |
| P0A7Q1 | 50S ribosomal protein L35 | rpmI | 0.189 | 5.49E-01 | -1.419 | 4.26E-02 | -1.632 | 6.95E-02 |

| | | | | | | | | |
|---|---|---|---|---|---|---|---|---|
| P33650 | Fe(2+) transporter | feoB | -0.366 | 4.49E-01 | -1.364 | 6.48E-04 | -1.635 | 8.31E-04 |
| P0AEG1 | Dipeptide transport system permease protein | dppC | 0.286 | 6.22E-02 | -1.361 | 3.00E-02 | -1.641 | 1.65E-03 |
| P0ABS8 | DNA polymerase III subunit theta | holE | -0.712 | 5.27E-02 | -1.080 | 2.94E-02 | -1.647 | 4.83E-02 |
| P65292 | Uncharacterized lipoprotein YgdI | ygdI | 0.335 | 4.29E-02 | -1.453 | 5.60E-03 | -1.656 | 3.55E-02 |
| P77213 | Putative glutamate--cysteine ligase 2 | ybdK | 0.029 | 8.09E-01 | -0.623 | 7.60E-02 | -1.657 | 1.42E-02 |
| P64432 | Inner membrane protein | ypjD | 0.323 | 1.98E-01 | -0.524 | 4.65E-01 | -1.673 | 2.33E-02 |
| P39172 | High-affinity zinc uptake system protein | znuA | 0.047 | 7.97E-01 | -0.568 | 9.77E-02 | -1.680 | 1.38E-02 |
| P39411 | Inosine/xanthosine triphosphatase | yjjX | -0.146 | 3.73E-01 | -0.281 | 2.11E-01 | -1.683 | 2.48E-02 |
| P09151 | 2-isopropylmalate synthase | leuA | -0.739 | 8.99E-02 | -1.240 | 3.19E-02 | -1.715 | 8.90E-03 |
| P77302 | Diguanylate cyclase | dgcM | -1.209 | 3.18E-02 | -1.727 | 1.87E-02 | -1.733 | 2.59E-02 |
| P76227 | Uncharacterized protein YnjH | ynjH | 0.076 | 1.62E-01 | -0.268 | 2.66E-01 | -1.760 | 4.44E-02 |
| P37188 | PTS system galactitol-specific EIIB component | gatB | -0.013 | 8.09E-01 | -0.721 | 2.83E-03 | -1.764 | 4.87E-04 |
| P0A6E1 | Shikimate kinase 2 | aroL | -0.494 | 1.36E-01 | -0.448 | 1.44E-01 | -1.774 | 3.96E-02 |
| P0AEM4 | Negative regulator of flagellin synthesis | flgM | -0.106 | 2.61E-01 | -0.561 | 1.26E-01 | -1.781 | 1.26E-03 |
| P0ACZ4 | DNA-binding transcriptional activator | evgA | 0.056 | 6.42E-01 | -0.405 | 1.08E-01 | -1.803 | 4.68E-03 |
| P0AGL5 | Ribosome association toxin | ratA | -0.857 | 3.40E-01 | -1.575 | 3.02E-02 | -1.810 | 4.39E-02 |
| P0A9S7 | High-affinity branched-chain amino acid transport ATP-binding protein | livG | -1.054 | 2.76E-02 | -1.332 | 1.23E-02 | -1.867 | 2.63E-03 |
| P0AEL3 | Fe(2+) transport protein A | feoA | -0.065 | 8.61E-01 | -1.288 | 5.00E-02 | -1.885 | 1.28E-02 |
| P37313 | Dipeptide transport ATP-binding protein | dppF | 0.207 | 2.03E-02 | -2.104 | 1.79E-02 | -1.911 | 6.37E-04 |
| P64638 | Probable [Fe-S]-dependent transcriptional repressor | feoC | -0.206 | 5.73E-01 | -1.901 | 5.04E-02 | -1.915 | 5.62E-03 |
| P0AB65 | Acylphosphatase | yccX | 0.189 | 8.23E-02 | -0.703 | 4.61E-02 | -1.919 | 1.98E-03 |
| P08660 | Lysine-sensitive aspartokinase 3 | lysC | -2.180 | 6.73E-04 | -1.550 | 1.29E-03 | -1.932 | 2.02E-03 |
| P0AAG0 | Dipeptide transport ATP-binding protein | dppD | 0.175 | 3.90E-01 | -1.891 | 2.50E-02 | -1.977 | 3.45E-02 |
| P0A6A6 | 3-isopropylmalate dehydratase large subunit | leuC | -0.566 | 3.69E-02 | -1.700 | 5.08E-03 | -2.009 | 1.55E-03 |
| P0AD96 | Leu/Ile/Val-binding protein | livJ | -0.695 | 1.39E-02 | -1.296 | 1.64E-02 | -2.054 | 1.08E-03 |
| P0AEF8 | Dipeptide transport system permease protein | dppB | 0.395 | 8.96E-02 | -1.965 | 2.16E-02 | -2.074 | 5.65E-03 |
| P39386 | Multidrug resistance protein | mdtM | 0.705 | 2.21E-02 | -0.008 | 9.90E-01 | -2.120 | 5.99E-03 |
| P0AAV6 | Uncharacterized protein YbgS | ybgS | 0.105 | 6.18E-01 | -0.684 | 8.73E-02 | -2.156 | 2.51E-02 |
| P0AB91 | Phospho-2-dehydro-3-deoxyheptonate aldolase, Phe-sensitive | aroG | -0.537 | 1.18E-01 | -2.401 | 4.99E-03 | -2.166 | 4.31E-03 |
| P30126 | 3-isopropylmalate dehydratase small subunit | leuD | -0.460 | 4.93E-02 | -1.571 | 1.08E-03 | -2.267 | 6.37E-02 |
| P00888 | Phospho-2-dehydro-3-deoxyheptonate aldolase, Tyr-sensitive | aroF | -1.599 | 1.51E-01 | -2.723 | 2.89E-02 | -2.431 | 1.97E-02 |
| P0A6R3 | DNA-binding protein | fis | -0.984 | 1.57E-01 | -2.634 | 2.00E-03 | -2.456 | 1.82E-02 |
| P0AA57 | Protein YobA | yobA | 0.077 | 4.97E-01 | -0.599 | 4.10E-02 | -2.522 | 2.28E-02 |
| P07021 | Putative lipoprotein | yfiB | 0.062 | 7.51E-01 | -1.406 | 1.78E-03 | -2.547 | 9.42E-03 |
| P06611 | Vitamin B12 import ATP-binding protein | btuD | 0.025 | 9.35E-01 | -0.267 | 4.16E-01 | -2.559 | 9.18E-03 |
| P37643 | Inner membrane metabolite transport protein | yhjE | -1.653 | 1.14E-02 | -2.096 | 7.03E-03 | -2.578 | 7.61E-03 |
| P27300 | Tetraacyldisaccharide 4'-kinase | lpxK | -0.324 | 7.49E-02 | -0.854 | 1.23E-02 | -2.918 | 2.84E-03 |
| P0ACW4 | Uncharacterized protein YdcA | ydcA | -0.562 | 9.34E-02 | -1.245 | 2.06E-01 | -3.053 | 2.46E-02 |
| P0CE48 | Elongation factor Tu 2 | tufB | 0.293 | 4.56E-01 | -0.645 | 2.82E-01 | -3.577 | 1.58E-02 |

LogFC, average log2 fold change (n=2). Heat map applied to LogFC values - shading in red indicate upregulation and blue indicate downregulation of protein abundance levels compared to control samples. Statistically significant values shown in black font ($p < 0.05$) and insignificant values ($p > 0.05$) in grey font as determined by *t*-test.

Table S2. *S. aureus* proteins with ≥ 2-fold abundance changes compared to untreated control.

| UniProt Accession | Protein Description | Gene Names | LED/Control | | FCD/Control | | LED-FCD/Control | |
|---|---|---|---|---|---|---|---|---|
| | | | LogFC | T.test | LogFC | T.test | LogFC | T.test |
| A0A3S4Q2Y7 | DNA repair protein | recN | 0.980 | 4.77E-01 | 7.177 | 1.37E-02 | 8.094 | 3.31E-02 |
| A0A1D4V4A1 | tRNA-binding domain-containing protein | | 1.484 | 1.84E-02 | 6.054 | 1.07E-02 | 7.516 | 1.19E-02 |
| A0A1Q8DEZ2 | Glutamate-1-semialdehyde 2,1-aminomutase | hemL | | | 4.652 | 8.87E-03 | 7.083 | 6.35E-03 |
| B8XPY6 | Uridylate kinase (Fragment) | | 1.367 | 2.59E-01 | 5.331 | 4.73E-02 | 6.877 | 5.69E-02 |
| A0A2X2L4R8 | Glyceraldehyde-3-phosphate dehydrogenase | gapA | 0.572 | 2.97E-01 | 5.169 | 7.14E-03 | 6.766 | 2.63E-02 |
| A0A454UY90 | Uncharacterized protein | | 1.264 | 1.29E-01 | 5.841 | 2.18E-02 | 6.721 | 3.24E-02 |
| A0A1Q8DGI2 | PTS lactose transporter subunit IIB | | 2.133 | 1.22E-01 | 4.970 | 5.26E-02 | 6.674 | 3.64E-02 |
| A0A3S5DZF0 | ABC transporter ATP-binding protein | yheS | -0.544 | 7.10E-01 | 5.167 | 1.88E-03 | 6.589 | 1.29E-02 |
| A0A1Q8DEE7 | ATP-dependent helicase/deoxyribonuclease subunit B | | 1.974 | 8.81E-02 | 5.910 | 1.78E-02 | 6.498 | 1.42E-02 |
| A0A1Q8DFI5 | Thymidylate synthase | thyA | 0.781 | 2.97E-01 | 5.294 | 4.66E-02 | 6.451 | 3.60E-02 |
| A0A077UG64 | Zinc-type alcohol dehydrogenase-like protein | | | | 4.259 | 8.22E-02 | 6.426 | 4.96E-02 |
| G8GLL3 | Dihydrofolate reductase (Fragment) | dhfr | 0.477 | 6.76E-02 | 5.056 | 7.94E-03 | 6.374 | 4.78E-04 |
| A0A0U1MLG2 | Phenylalanine--tRNA ligase beta subunit | pheT | 0.906 | 2.01E-02 | 4.631 | 3.95E-02 | 6.287 | 1.90E-02 |
| A0A380ESP1 | Uncharacterized protein conserved in bacteria | | 1.828 | 3.57E-02 | 4.853 | 4.45E-03 | 6.265 | 4.90E-03 |
| A0A1Q8DCU0 | Uncharacterized protein | | 0.082 | 8.27E-01 | 3.793 | 8.46E-03 | 6.167 | 1.92E-02 |
| A0A077UK35 | DNA binding protein | | 1.414 | 4.10E-02 | 5.189 | 2.65E-02 | 6.090 | 1.39E-02 |
| A0A3S4LUN1 | Exonuclease | sbcC | 0.201 | 6.49E-01 | 4.955 | 2.88E-02 | 6.020 | 1.24E-02 |
| A0A447Z887 | Putative lipoprotein | | 1.428 | 8.80E-02 | 5.179 | 7.45E-03 | 5.996 | 4.00E-03 |
| A0A1D4Q690 | Alkyl hydroperoxide reductase protein C | ahpC | 0.066 | 8.75E-01 | 4.311 | 4.04E-02 | 5.894 | 9.36E-03 |
| A0A380EJI8 | Tripeptidase T | | 0.933 | 4.60E-02 | 4.265 | 4.20E-03 | 5.881 | 1.89E-02 |
| A0A1D5A2I9 | Pyridoxine biosynthesis glutamine amidotransferase, synthase subunit | pdxS | 1.168 | 2.14E-01 | 5.171 | 4.42E-02 | 5.727 | 4.46E-02 |
| A0A0Y9UAH8 | Transposon-related protein | | 0.611 | 1.54E-01 | 3.718 | 6.30E-03 | 5.703 | 6.03E-03 |
| A0A1D5AC31 | Thioredoxin | trxA | -0.034 | 9.66E-01 | 3.450 | 9.67E-02 | 5.682 | 2.41E-02 |
| A0A380ERY3 | DNA-directed RNA polymerase subunit beta' | rpoC | 0.816 | 1.39E-01 | 4.682 | 2.57E-02 | 5.615 | 2.06E-02 |
| A0A0B4N9S6 | DUF2977 domain-containing protein | | -0.007 | 9.90E-01 | 4.128 | 1.90E-02 | 5.567 | 1.05E-02 |
| A0A380E0K8 | Transketolase | tkt | 0.442 | 4.33E-01 | 3.488 | 1.49E-02 | 5.536 | 2.16E-02 |
| A0A380DFX1 | Enolase | eno | 0.557 | 1.31E-01 | 4.193 | 4.58E-03 | 5.526 | 7.50E-03 |
| A0A1Q8DGM2 | Pantothenate synthetase | panC | 0.445 | 4.76E-01 | 4.286 | 3.95E-02 | 5.517 | 4.36E-02 |
| A0A380DU17 | Hydroxymethylbilane synthase | hemC | 0.412 | 1.30E-01 | 4.987 | 8.78E-04 | 5.499 | 1.13E-03 |
| A0A2S6DE03 | Glycine--tRNA ligase (Fragment) | | 0.640 | 3.39E-01 | 4.553 | 4.24E-02 | 5.491 | 3.56E-02 |
| A0A133PWH8 | Uncharacterized protein | | 0.005 | 9.95E-01 | 3.782 | 2.43E-03 | 5.460 | 1.42E-02 |
| A0A380DGI6 | Carbamoyl-phosphate synthase (glutamine-hydrolyzing) | carB | -0.022 | 9.85E-01 | 4.912 | 5.98E-03 | 5.345 | 2.69E-04 |
| A0A299MHJ1 | Uncharacterized protein | | 1.215 | 2.23E-01 | 4.555 | 3.90E-02 | 5.306 | 5.80E-02 |
| A0A380E0V1 | Serine/threonine protein kinase | prkC | 0.127 | 4.66E-01 | 3.738 | 4.02E-02 | 5.253 | 1.14E-05 |
| A0A3S4Q4P8 | 3-methyl-2-oxobutanoate hydroxymethyltransferase | panB | 0.288 | 2.28E-01 | 4.039 | 1.26E-02 | 5.228 | 1.34E-02 |
| Q93IA5 | Uncharacterized protein OS=Staphylococcus aureus | | 0.074 | 9.12E-01 | 3.811 | 3.87E-02 | 5.216 | 5.91E-02 |
| A0A2X2P2D7 | DNA helicase OS=Staphylococcus aureus | pcrA | 0.230 | 7.15E-01 | 4.241 | 2.80E-02 | 5.205 | 1.46E-02 |
| A0A380DDN8 | Haloacid dehalogenase-like family hydrolase | | 1.126 | 2.03E-01 | 3.705 | 4.11E-02 | 5.170 | 1.35E-02 |
| A0A3S4M7N2 | Haloacid dehalogenase-like family hydrolase | hdl IVa | 0.315 | 6.15E-01 | 4.096 | 2.30E-03 | 5.137 | 8.28E-04 |
| A0A1D4R433 | Phi 11 orf40 OS=Staphylococcus aureus | | 0.472 | 6.32E-02 | 3.341 | 1.74E-04 | 5.135 | 3.53E-02 |
| A0A380DFZ1 | Nucleotide-binding protein NCTC7887_02050 | | 2.467 | 9.95E-03 | 5.356 | 4.47E-02 | 5.119 | 5.07E-04 |
| A0A380DY12 | Glycine betaine transporter | opuD | 1.214 | 7.32E-02 | 4.053 | 1.04E-02 | 5.077 | 7.43E-03 |
| A0A0U1MUW7 | Bacitracin ABC efflux transporter (ATP-binding protein) | bceA | 0.174 | 5.77E-01 | 4.973 | 5.93E-04 | 5.075 | 1.07E-02 |
| A0A0U1MRQ2 | Ribose-5-phosphate isomerase A | rpi | 1.327 | 1.62E-01 | 5.117 | 2.05E-02 | 5.063 | 2.54E-02 |
| A0A2S6DA54 | Elongation factor Ts (Fragment) | | 0.894 | 7.00E-02 | 4.346 | 1.62E-03 | 5.050 | 1.53E-03 |

| | | | | | | | | |
|---|---|---|---|---|---|---|---|---|
| A0A077UGS9 | Putative lipoprotein | | 0.165 | 7.82E-01 | 2.809 | 4.85E-02 | 5.042 | 3.78E-02 |
| A0A380DJK7 | Secreted von Willebrand factor-binding protein | vwbp | 0.709 | 3.72E-01 | 3.847 | 5.41E-02 | 5.040 | 1.12E-02 |
| A0A077UJG8 | Aspartokinase | lysC2 | 1.180 | 2.72E-02 | 3.714 | 3.68E-03 | 5.008 | 4.24E-03 |
| A8ILY9 | Protein Rep | rep | 1.951 | 5.08E-02 | 5.288 | 1.40E-02 | 5.006 | 8.44E-03 |
| A0A447ZCY2 | Cytosolic protein | | 1.344 | 1.01E-01 | 3.780 | 1.65E-02 | 4.980 | 3.07E-02 |
| A0A1D4NWD7 | Immunoglobulin G-binding protein A | spa | -0.252 | 1.17E-01 | 3.220 | 4.58E-03 | 4.977 | 4.21E-02 |
| A0A0U1MH75 | Uncharacterized protein | | 0.167 | 6.91E-01 | 3.683 | 9.33E-03 | 4.966 | 2.10E-03 |
| A0A0H2B9X7 | Phage tail protein | | 0.568 | 2.22E-01 | 3.697 | 3.42E-02 | 4.941 | 5.32E-03 |
| A0A380DW86 | Purine nucleosidase | | 0.264 | 3.85E-01 | 3.516 | 1.70E-02 | 4.925 | 1.42E-02 |
| A0A380ER49 | Protein RecA | recA | 0.147 | 6.17E-01 | 3.357 | 1.36E-02 | 4.888 | 4.21E-04 |
| E2G4H9 | von Willebrand factor-binding protein | vwb | 0.756 | 6.37E-02 | 3.827 | 3.88E-03 | 4.856 | 8.34E-03 |
| A0A2S6D2D4 | Formate acetyltransferase (Fragment) | | -1.026 | 1.42E-01 | 3.035 | 1.41E-02 | 4.849 | 4.82E-03 |
| A0A447ZDB0 | 50S ribosomal protein L7/L12 | rplL | 0.395 | 5.64E-01 | 4.045 | 8.65E-03 | 4.804 | 3.57E-02 |
| A0A168SSP8 | SLT orf 129-like protein | | 0.911 | 7.53E-02 | 3.623 | 3.20E-02 | 4.797 | 4.53E-03 |
| A0A0K2D7F6 | Acetyl coenzyme A acetyltransferase (Fragment) | yqiL | 0.615 | 2.24E-01 | 3.718 | 1.48E-02 | 4.777 | 3.51E-02 |
| A0A380DC15 | Lipoprotein | lpl2 | 1.200 | 4.10E-02 | 3.612 | 2.69E-02 | 4.771 | 6.63E-04 |
| A0A3S4M5J4 | ATP synthase gamma chain | atpG | 1.283 | 3.09E-02 | 4.464 | 4.38E-03 | 4.765 | 1.56E-02 |
| A0A380DHT4 | GMP synthase [glutamine-hydrolyzing] | guaA | 0.427 | 1.34E-01 | 3.614 | 1.26E-02 | 4.761 | 3.60E-05 |
| D2JA14 | Glucose-6-phosphate isomerase | pgi | -1.071 | 1.90E-01 | 3.704 | 4.94E-03 | 4.755 | 4.69E-03 |
| A0A380EIP6 | Valyl-tRNA synthetase | valS | 1.537 | 2.21E-03 | 4.806 | 2.05E-02 | 4.751 | 1.30E-02 |
| A0A380EN78 | N-acetylglucosamine-1-phosphate uridyltransferase eukaryotic | | 0.627 | 1.81E-01 | 4.114 | 5.75E-03 | 4.744 | 5.99E-03 |
| A0A380EUQ5 | Two component transcriptional regulator, AraC family | | 0.875 | 2.63E-01 | 3.856 | 4.93E-02 | 4.724 | 5.81E-02 |
| P22874 | Mercuric resistance operon regulatory protein | merR | 0.596 | 4.65E-01 | 5.080 | 4.59E-02 | 4.700 | 3.59E-02 |
| A0A380DIN8 | Pyridoxal 5'-phosphate synthase subunit | pdxS | 0.791 | 2.33E-01 | 3.718 | 1.62E-02 | 4.666 | 2.30E-02 |
| A0A0E7PD33 | ABC transporter permease | | 0.193 | 4.22E-01 | 4.194 | 1.38E-03 | 4.612 | 6.22E-04 |
| A0A2S6D3V2 | Iron-sulfur cluster carrier protein | apbC | 0.957 | 1.71E-01 | 4.416 | 4.66E-02 | 4.597 | 2.19E-02 |
| A0A098H1P9 | Phi 11 orf38 OS=Staphylococcus aureus | | 0.721 | 3.66E-02 | 3.048 | 8.57E-04 | 4.590 | 3.95E-02 |
| A0A3S4LCK0 | 30S ribosomal protein S17 | rpsQ | 0.802 | 1.05E-01 | 3.654 | 5.57E-02 | 4.582 | 3.01E-03 |
| A0A1D4R989 | Putative transcriptional regulator | | 0.433 | 3.50E-01 | 3.352 | 5.12E-02 | 4.581 | 1.84E-02 |
| A0A0E8HDD8 | DUF3310 domain-containing protein | | 0.457 | 2.81E-01 | 2.923 | 1.59E-02 | 4.560 | 1.99E-02 |
| A0A077V028 | Heptaprenyl diphosphate synthase (HEPPP synthase) subunit 1 family protein | | 1.878 | 4.60E-02 | 4.910 | 4.65E-03 | 4.553 | 2.30E-03 |
| A0A380EHB7 | Catalase | katA | 0.503 | 4.24E-01 | 3.540 | 2.63E-02 | 4.540 | 2.19E-02 |
| A0A380E4R0 | Phage protein | | -0.023 | 9.16E-01 | 2.608 | 2.25E-02 | 4.535 | 5.12E-02 |
| D7RHA4 | SpA IgG-binding domain protein (Fragment) | | 0.095 | 9.32E-01 | 4.338 | 8.66E-02 | 4.529 | 2.28E-03 |
| A0A2X2L1A1 | Polyribonucleotide nucleotidyltransferase | pnp | 1.235 | 1.92E-01 | 3.983 | 2.48E-02 | 4.503 | 5.05E-02 |
| A0A2S6DJB3 | Uridylate kinase | pyrH | 1.554 | 2.77E-02 | 4.236 | 7.44E-04 | 4.500 | 2.41E-03 |
| A0A2I7Y983 | Uncharacterized protein | | -0.053 | 7.94E-01 | 3.429 | 3.88E-03 | 4.485 | 2.02E-03 |
| A0A1L1J5M4 | Holin, phage phi LC3 family | | 0.956 | 2.40E-02 | 2.935 | 8.67E-03 | 4.464 | 3.38E-02 |
| A0A0B6XQS2 | Phage protein | | 0.006 | 9.75E-01 | 2.986 | 6.87E-04 | 4.440 | 5.24E-02 |
| Q8KRL7 | Guanylate kinase (Fragment) | gmk | 0.272 | 4.56E-01 | 3.631 | 7.29E-03 | 4.429 | 1.01E-03 |
| A0A1D5A4B3 | Aspartyl/glutamyl-tRNA amidotransferase subunit B | gatB | 0.473 | 4.79E-02 | 4.020 | 2.91E-02 | 4.410 | 3.49E-03 |
| A0A447Z788 | Type I restriction-modification system specificity protein | | 0.562 | 3.43E-01 | 3.892 | 1.34E-02 | 4.406 | 9.42E-03 |
| P0C1V1 | Delta-hemolysin | hld | 0.201 | 5.32E-01 | 3.472 | 5.18E-04 | 4.391 | 2.78E-04 |
| A0A0U1MV41 | Luciferase family protein | limB | 0.619 | 3.62E-01 | 5.036 | 5.36E-02 | 4.379 | 2.27E-02 |
| A0A2X2MDU5 | Phosphocarrier protein | ptsH | 0.917 | 5.91E-02 | 4.114 | 6.02E-03 | 4.368 | 3.57E-03 |
| J7FMN4 | Guanylate kinase (Fragment) | gmk | 0.102 | 7.68E-01 | 2.608 | 1.84E-02 | 4.365 | 1.80E-02 |
| A0A2S6CZZ3 | DUF2479 domain-containing protein | | 0.563 | 3.11E-01 | 3.207 | 4.27E-02 | 4.365 | 1.05E-02 |
| A0A2X2M2Y9 | Two component transcriptional regulator VraR, LuxR family | vraR | 0.407 | 5.72E-01 | 3.565 | 4.09E-03 | 4.360 | 3.37E-03 |

| | | | | | | | | |
|---|---|---|---|---|---|---|---|---|
| A0A0U1MR72 | Uncharacterized protein | | 0.055 | 8.50E-01 | 3.379 | 5.61E-04 | 4.353 | 3.99E-02 |
| A0A2W7VUR2 | Phage protein | | 0.259 | 2.38E-01 | 2.949 | 2.97E-02 | 4.320 | 5.09E-02 |
| A0A380DDY3 | Alanine--tRNA ligase | alaS | 0.912 | 6.46E-02 | 4.013 | 1.88E-03 | 4.317 | 1.90E-02 |
| A0A0U1MTQ7 | Secretory antigen ssaA2 | | -0.606 | 2.61E-01 | 3.711 | 1.75E-02 | 4.263 | 3.00E-02 |
| A0A2S6DJ04 | Redox-regulated ATPase YchF (Fragment) | | 0.233 | 3.94E-01 | 3.696 | 2.18E-02 | 4.247 | 2.44E-02 |
| A0A1D4SUC0 | Fructose-bisphosphate aldolase | fba | 0.759 | 2.56E-01 | 2.925 | 8.00E-03 | 4.194 | 1.06E-03 |
| A0A2X2JW40 | Lipoate-protein ligase A | lipM | 0.593 | 3.72E-01 | 2.355 | 1.46E-01 | 4.192 | 3.79E-02 |
| A0A2W2VJ18 | Phage protein | | 0.639 | 2.21E-01 | 2.810 | 1.84E-02 | 4.164 | 4.02E-02 |
| A0A447ZMA9 | Serine-aspartate repeat-containing protein E | sdrE | 0.268 | 5.81E-01 | 2.240 | 4.28E-02 | 4.153 | 3.57E-02 |
| A0A380EVE1 | Cytidine deaminase | cdd | 0.530 | 1.36E-01 | 3.220 | 5.00E-04 | 4.148 | 1.93E-03 |
| A0A2X2MY88 | Oligopeptide transporter putative substrate binding domain protein | opp-1A | -0.033 | 9.60E-01 | 1.680 | 3.33E-02 | 4.126 | 9.00E-03 |
| A0A166KHF6 | Phi 11 orf35 | | 0.452 | 1.59E-01 | 3.420 | 1.61E-02 | 4.056 | 3.61E-02 |
| A0A0D3QAF5 | IgG-binding protein SBI | sbi | 0.331 | 1.76E-01 | 4.304 | 2.62E-03 | 4.016 | 1.95E-03 |
| A0A3S4NGC9 | Formate--tetrahydrofolate ligase | fhs | 0.255 | 4.28E-01 | 2.679 | 2.35E-02 | 4.004 | 1.30E-03 |
| A0A454V147 | DUF4355 domain-containing protein | | 0.115 | 4.91E-01 | 2.702 | 6.92E-04 | 3.953 | 4.50E-02 |
| A0A2X2K5R6 | SceD-like transglycosylase, biomarker for vancomycin-intermediate strains | sceD | 0.327 | 3.30E-01 | 2.358 | 2.00E-02 | 3.950 | 5.62E-03 |
| A0A447Z955 | Glutamine--fructose-6-phosphate aminotransferase [isomerizing] | glmS | 1.035 | 2.57E-01 | 3.823 | 5.06E-02 | 3.942 | 2.89E-02 |
| A0A1Q8DEN7 | Orotate phosphoribosyltransferase OS=Staphylococcus aureus | pyrE | 0.821 | 7.71E-02 | 3.140 | 1.81E-02 | 3.925 | 3.02E-02 |
| A0A3S4MWG9 | ATP-dependent helicase/nuclease subunit A | rexA | 0.824 | 1.54E-01 | 3.619 | 1.70E-02 | 3.924 | 5.93E-03 |
| A0A1Q8DFC0 | Uncharacterized protein | | 0.874 | 9.77E-02 | 4.057 | 3.40E-02 | 3.915 | 2.65E-02 |
| A0A447ZBP7 | dITP/XTP pyrophosphatase | rdgB | 0.688 | 1.17E-01 | 3.125 | 9.22E-03 | 3.904 | 4.23E-03 |
| A0A380DPX8 | ATP-dependent endopeptidase hsl ATP-binding subunit | hslU | 0.370 | 2.33E-01 | 2.819 | 3.59E-02 | 3.898 | 6.45E-03 |
| A0A077UJM5 | Putative anthranilate synthase component I | trpE | 0.783 | 2.02E-01 | 3.349 | 4.07E-02 | 3.891 | 3.94E-02 |
| A0A380DFY2 | D-3-phosphoglycerate dehydrogenase | serA | 0.460 | 4.56E-01 | 3.858 | 3.87E-02 | 3.889 | 5.92E-02 |
| A0A0U1MTK5 | Dehydrosqualene desaturase | crtN | 1.716 | 1.37E-02 | 4.282 | 7.79E-03 | 3.885 | 7.78E-02 |
| A0A380DWF4 | Chorismate mutase I / 2-keto-3-deoxy-D-arabino-heptulosonate-7-phosphate synthase I beta, AroH/AroA I beta | aroA | 0.729 | 4.41E-01 | 3.787 | 3.11E-02 | 3.884 | 4.15E-02 |
| A0A380EGE7 | Protein translation elongation factor G (EF-G) | fusA | 0.543 | 4.00E-01 | 3.488 | 2.39E-02 | 3.833 | 2.27E-02 |
| A0A3G1QDW6 | Immunoglobulin G-binding protein A (Fragment) | | -0.387 | 3.38E-01 | 3.105 | 9.12E-03 | 3.827 | 2.96E-02 |
| A0A1Q8DGJ1 | Dihydrolipoyl dehydrogenase | | -0.021 | 9.60E-01 | 2.568 | 2.20E-02 | 3.825 | 1.40E-02 |
| A0A077UVU9 | Putative GTPases (G3E family) | cobW | 0.295 | 3.85E-01 | 3.651 | 3.75E-02 | 3.793 | 9.30E-04 |
| A0A2S6DA08 | Cell division protein FtsZ (Fragment) | | 0.884 | 4.55E-02 | 3.636 | 4.07E-03 | 3.758 | 5.44E-03 |
| A0A380EGL9 | Glucose-6-phosphate isomerase | pgi | 1.292 | 1.57E-01 | 4.355 | 4.74E-02 | 3.720 | 6.09E-02 |
| P0C1U8 | Glutamyl endopeptidase | sspA | 0.024 | 8.86E-01 | 3.312 | 2.48E-02 | 3.715 | 4.06E-03 |
| A0A077UN86 | Putative surface anchored protein | fmtB | 0.216 | 3.87E-01 | 3.559 | 6.34E-03 | 3.714 | 6.60E-04 |
| A0A1Q8DC47 | Cell division protein | ftsK | 0.917 | 2.31E-01 | 3.463 | 9.31E-03 | 3.711 | 3.51E-02 |
| A0A380DMU0 | Polyketide synthase | pksM | -0.256 | 5.76E-01 | 3.693 | 4.57E-04 | 3.708 | 6.63E-02 |
| A0A0U1MV09 | Putative N-acetylmannosamine-6-phosphate 2-epimerase | nanE | 0.379 | 4.87E-01 | 3.366 | 4.99E-02 | 3.688 | 5.14E-02 |
| A0A380EUG1 | Ribosome-binding factor A | rbfA | 2.106 | 8.59E-02 | 4.515 | 1.76E-02 | 3.687 | 3.55E-02 |
| A0A0U1N070 | ATP-dependent 6-phosphofructokinase | pfkA | 0.108 | 7.42E-01 | 2.323 | 2.90E-02 | 3.660 | 6.53E-03 |
| A0A077VMD2 | Trigger factor | tig | 0.845 | 8.19E-02 | 3.578 | 2.07E-03 | 3.594 | 2.72E-02 |
| A0A380DFT4 | Aspartyl/glutamyl-tRNA(Asn/Gln) amidotransferase subunit C | gatC | -0.429 | 6.12E-01 | 1.419 | 2.34E-01 | 3.591 | 1.14E-03 |
| A0A2X2KY14 | Putative rRNA methyltransferase | spoU | 1.327 | 1.62E-01 | 3.694 | 4.78E-02 | 3.586 | 1.58E-02 |
| A0A0C6EG82 | Phage head-tail adapter protein | | -0.126 | 6.14E-01 | 2.247 | 3.53E-02 | 3.563 | 4.97E-03 |
| A0A380EMM0 | Chaperonin GroEL | groEL | 0.297 | 1.07E-01 | 2.763 | 3.66E-03 | 3.562 | 3.18E-02 |
| A0A380DMP1 | Ornithine aminotransferase | rocD2 | 0.569 | 1.54E-01 | 2.466 | 1.49E-02 | 3.558 | 9.27E-03 |
| A0A077ULE8 | O-methyltransferase family protein | | 1.159 | 7.69E-02 | 3.911 | 8.66E-03 | 3.530 | 1.16E-02 |

| | | | | | | | | |
|---|---|---|---|---|---|---|---|---|
| A0A2X2MYH1 | Uncharacterized protein | | 1.726 | 2.25E-02 | 2.804 | 2.86E-02 | 3.530 | 1.69E-02 |
| A0A1D4RVK1 | Putative cell-division initiation protein | divIVA | -0.127 | 1.41E-01 | 2.666 | 3.45E-03 | 3.499 | 3.11E-03 |
| A0A1Q8DC44 | Pyridoxal 5'-phosphate synthase subunit | pdxS | 1.206 | 1.28E-01 | 3.525 | 3.60E-03 | 3.495 | 2.02E-02 |
| A0A380EGQ0 | Phi 11 orf36 | | 0.388 | 1.39E-01 | 2.289 | 5.44E-03 | 3.463 | 6.59E-03 |
| A0A3S4L8S9 | Holliday junction ATP-dependent DNA helicase | ruvA | -0.381 | 4.64E-01 | 3.009 | 5.63E-03 | 3.441 | 4.24E-02 |
| A0A0U1MGK3 | Phage protein | | 0.272 | 2.10E-01 | 2.869 | 1.26E-02 | 3.413 | 4.02E-03 |
| A0A0U1MTI2 | 3-hydroxy-3-methylglutaryl coenzyme A reductase | mvaA | -0.220 | 5.43E-01 | 3.037 | 2.15E-02 | 3.407 | 2.90E-03 |
| A0A1M4PZD7 | Transcriptional regulator, y4mF family | | -0.165 | 6.74E-01 | 1.967 | 1.51E-02 | 3.372 | 2.31E-02 |
| A0A2Z6FJG4 | Immunoglobulin G binding protein (Fragment) | spa | -0.032 | 6.70E-01 | 3.040 | 1.18E-02 | 3.354 | 2.26E-02 |
| A0A2S6DEP8 | ATP-dependent Clp protease proteolytic subunit | | 0.206 | 6.87E-03 | 2.487 | 1.30E-04 | 3.346 | 1.00E-02 |
| I7GSD0 | Ampicillin resistance protein | ear | 0.119 | 5.06E-01 | 2.264 | 9.50E-02 | 3.345 | 1.92E-02 |
| A0A077UIP3 | Isovaleryl-CoA dehydrogenase | | 0.196 | 2.21E-01 | 3.137 | 1.34E-02 | 3.343 | 2.20E-02 |
| A0A454H234 | Uncharacterized protein | | 0.269 | 4.91E-01 | 2.258 | 5.57E-02 | 3.341 | 4.19E-02 |
| A0A0H2HU80 | Acyl carrier protein | acpP | 0.520 | 3.12E-01 | 3.268 | 5.18E-02 | 3.340 | 3.04E-02 |
| A0A380DEM8 | Catalase | katA | 0.318 | 2.72E-01 | 2.624 | 2.46E-02 | 3.338 | 2.29E-02 |
| A0A1D5AC17 | Thioredoxin reductase | trxB | 0.265 | 1.56E-01 | 2.413 | 4.02E-02 | 3.311 | 5.20E-04 |
| A0A1D4R360 | Phage protein | | 0.223 | 2.42E-02 | 1.498 | 3.07E-02 | 3.306 | 8.03E-02 |
| Q7BWE0 | Vancomycin response regulator | vanR | 0.262 | 1.37E-01 | 2.517 | 7.87E-02 | 3.305 | 1.58E-03 |
| A0A1Q8DHB6 | Alpha/beta hydrolase | | -0.567 | 5.10E-01 | 3.789 | 2.54E-02 | 3.282 | 1.99E-02 |
| A0A0U1MGN7 | Putative major head protein | | 0.105 | 4.32E-01 | 1.886 | 1.31E-02 | 3.268 | 5.17E-02 |
| A0A380EC03 | Immunoglobulin G-binding protein A | spa | 0.047 | 3.57E-01 | 3.012 | 1.49E-02 | 3.268 | 2.79E-02 |
| A0A1Q8DD39 | UPF0342 protein BSZ10_08550 | | 0.572 | 3.38E-01 | 2.320 | 1.57E-01 | 3.261 | 1.27E-02 |
| A0A0C6EXD4 | Alanyl-tRNA synthetase | | 0.313 | 3.21E-01 | 1.894 | 2.46E-02 | 3.254 | 6.22E-02 |
| J7FMM4 | Triosephosphate isomerase (Fragment) | tpi | 0.102 | 8.23E-01 | 2.296 | 3.48E-02 | 3.252 | 8.15E-03 |
| A0A380DHQ1 | Immunoglobulin G binding protein A | spa | 0.105 | 2.97E-01 | 2.692 | 1.06E-03 | 3.223 | 2.07E-02 |
| A0A2X2N386 | Phage tail fiber protein | | 0.023 | 8.75E-01 | 1.755 | 1.67E-03 | 3.209 | 4.64E-02 |
| A0A1Q8DB13 | HAD family hydrolase | | 0.765 | 2.83E-02 | 2.035 | 6.80E-02 | 3.190 | 3.22E-02 |
| A0A1Q8DAY2 | SUF system NifU family Fe-S cluster assembly protein | | 1.394 | 3.70E-02 | 3.194 | 7.88E-02 | 3.189 | 5.27E-02 |
| A0A380DIP9 | DNA-directed RNA polymerase subunit | rpoC | 0.164 | 3.57E-01 | 1.642 | 1.28E-02 | 3.185 | 2.39E-02 |
| A0A224AX52 | Phi ETA orf 18-like protein | | 0.150 | 3.05E-01 | 2.043 | 2.87E-02 | 3.171 | 6.69E-03 |
| A0A1Q8DEY5 | Ferritin | | 0.079 | 1.01E-01 | 1.165 | 1.30E-01 | 3.165 | 6.14E-04 |
| D9I4C8 | Glyceraldehyde-3-phosphate dehydrogenase (Fragment) | gap | 0.214 | 4.67E-01 | 2.884 | 1.79E-02 | 3.163 | 8.74E-03 |
| A0A380ESC5 | Elongation factor G | fus | 0.159 | 4.19E-01 | 2.368 | 2.61E-02 | 3.162 | 2.23E-02 |
| A0A447ZC75 | Oligoendopeptidase F | pepF | 0.335 | 4.77E-01 | 2.687 | 2.66E-02 | 3.118 | 3.32E-02 |
| A0A133Q3F8 | Arginine repressor | argR | 0.137 | 1.60E-01 | 2.003 | 1.95E-02 | 3.074 | 1.43E-04 |
| A0A0B4NAP7 | Phage major tail protein, TP901-1 family phi 11 orf39 | | 0.379 | 1.08E-01 | 1.490 | 3.53E-02 | 3.074 | 2.27E-02 |
| A0A380ERY8 | Phosphoribosylaminoimidazole-succinocarboxamide synthase | purC | -0.183 | 6.33E-01 | 1.966 | 3.42E-02 | 3.064 | 2.56E-02 |
| A0A077ULL9 | Lipase family protein | | 2.157 | 3.31E-02 | 3.277 | 6.08E-02 | 3.062 | 9.03E-02 |
| A0A077UJ42 | Peptidase T | pepT | -0.354 | 1.15E-01 | 1.616 | 1.35E-01 | 3.056 | 4.65E-03 |
| A0A2X2KW44 | Probable transcriptional regulatory protein NCTC13131_03956 | | 0.792 | 2.39E-01 | 3.193 | 2.65E-03 | 3.053 | 1.70E-02 |
| A0A380DD11 | UPF0340 protein NCTC7887_00871 | | 0.622 | 8.39E-02 | 3.494 | 2.86E-02 | 3.032 | 7.27E-04 |
| A0A380EFQ6 | ABC transporter ATP-binding protein VraF | macB | 0.472 | 1.64E-01 | 1.575 | 2.91E-02 | 3.017 | 6.48E-03 |
| A0A090LUI7 | Biotin transporter | bioY | 0.412 | 1.36E-01 | 1.385 | 2.95E-02 | 2.993 | 2.68E-01 |
| A0A145MQ35 | Phage family protein | | 1.142 | 4.13E-02 | 2.145 | 1.32E-02 | 2.980 | 4.79E-03 |
| A0A1Q8DFE0 | Phosphate import ATP-binding protein | pstB | 0.389 | 3.06E-02 | 2.863 | 2.64E-03 | 2.976 | 9.39E-04 |
| A0A1Q8DAX8 | Thiol reductase thioredoxin OS=Staphylococcus aureus | | 0.434 | 1.42E-01 | 2.831 | 6.00E-03 | 2.967 | 5.20E-03 |
| A0A3S4M1K7 | Phage-related cell wall hydrolase | atl | 2.049 | 1.37E-01 | 2.626 | 3.00E-01 | 2.938 | 3.33E-02 |
| A0A077UMZ5 | Glutamyl-tRNA(Gln) amidotransferase subunit A | gatA | -0.568 | 5.87E-02 | 2.627 | 9.34E-04 | 2.932 | 3.89E-03 |

| | | | | | | | | |
|---|---|---|---|---|---|---|---|---|
| A0A380DF06 | CTP synthase | pyrG | 1.014 | 2.78E-01 | 2.444 | 2.18E-03 | 2.924 | 1.30E-03 |
| A0A1Q8DEG2 | Catabolite control protein A | | 0.649 | 2.56E-01 | 2.426 | 3.12E-02 | 2.912 | 1.32E-02 |
| A0A380DTM7 | 5'-methylthioadenosine/S-adenosylhomocysteine nucleosidase | mtnN | -0.043 | 9.39E-01 | 1.823 | 1.18E-02 | 2.896 | 2.44E-04 |
| A0A1D4W7N6 | Phage protein | | 0.042 | 8.93E-01 | 1.818 | 5.43E-02 | 2.889 | 4.03E-02 |
| B9V2C1 | Copper-transporting ATPase | copA | 0.273 | 1.28E-01 | 2.523 | 8.87E-03 | 2.887 | 2.01E-03 |
| A0A2S6D6R6 | Aconitate hydratase (Fragment) | | 1.053 | 1.66E-01 | 3.933 | 1.49E-02 | 2.851 | 2.42E-02 |
| A0A2X2KR97 | Mobile element-associated integrase, putative | int | 0.295 | 3.54E-01 | 2.923 | 7.35E-03 | 2.829 | 2.43E-02 |
| A0A167TXZ9 | N5-carboxyaminoimidazole ribonucleotide synthase | purK | 0.208 | 3.88E-01 | 1.718 | 1.73E-02 | 2.822 | 7.48E-03 |
| A0A484W4Z4 | Zinc uptake regulation protein | zur | 0.515 | 3.55E-01 | 2.617 | 1.11E-02 | 2.819 | 1.45E-02 |
| D2J7K3 | Uncharacterized protein | | -0.289 | 2.70E-02 | 2.479 | 8.64E-03 | 2.812 | 3.73E-04 |
| A0A1Q8DDM9 | DtxR family transcriptional regulator | | -0.447 | 3.18E-02 | 2.365 | 2.39E-03 | 2.808 | 5.10E-03 |
| A0A2S6DJ21 | Glutamate dehydrogenase (Fragment) | | 0.457 | 3.24E-01 | 2.312 | 5.00E-02 | 2.798 | 2.75E-02 |
| A0A0U1MU80 | Phosphotransferase enzyme family protein | | 0.304 | 4.44E-01 | 2.909 | 2.46E-02 | 2.780 | 2.76E-02 |
| A0A380DI34 | Delta-1-pyrroline-5-carboxylate dehydrogenase | pruA | 0.404 | 3.79E-01 | 2.725 | 5.17E-02 | 2.767 | 3.58E-02 |
| A0A0Y8ZK60 | Uncharacterized protein | | 0.354 | 1.90E-01 | 1.847 | 2.16E-02 | 2.753 | 4.25E-03 |
| A0A380EQJ6 | Ser-Asp rich fibrinogen-binding/bone sialoprotein-binding protein | bbp | 0.470 | 3.31E-01 | 2.115 | 3.75E-03 | 2.737 | 1.06E-02 |
| A0A077UMR5 | Oligopeptide transport system ATP-binding protein | oppF2 | 0.710 | 3.43E-01 | 2.391 | 1.10E-01 | 2.713 | 4.99E-02 |
| A0A1Q8DDC8 | Alkyl hydroperoxide reductase subunit F | | 0.275 | 8.70E-02 | 1.841 | 4.88E-02 | 2.667 | 9.76E-04 |
| A0A2S6D048 | Peptidase | | 1.002 | 9.19E-02 | 2.815 | 2.98E-02 | 2.643 | 1.04E-02 |
| A0A077V0E2 | Alpha-amylase | malA | 0.035 | 8.86E-01 | 1.927 | 1.31E-02 | 2.632 | 3.69E-03 |
| A0A0C6E126 | Putative phage head morphogenesis protein | | 0.102 | 3.88E-01 | 1.735 | 3.56E-03 | 2.615 | 1.15E-02 |
| D2J6J4 | Recombinase | | 0.013 | 9.80E-01 | 1.576 | 8.78E-02 | 2.584 | 4.04E-02 |
| A0A2X2JTI4 | 3-oxoacyl-[acyl-carrier-protein] synthase, KASIII | fabH | -0.491 | 1.41E-01 | 1.678 | 1.15E-01 | 2.571 | 2.74E-02 |
| A0A1Q8DDW4 | GTP-binding protein | | 0.012 | 9.48E-01 | 2.045 | 4.13E-02 | 2.568 | 9.79E-03 |
| E4Z841 | ABC transporter | vga(C) | 0.995 | 2.21E-03 | 3.278 | 1.80E-02 | 2.554 | 4.62E-04 |
| A0A2X3UJS1 | Protein serine/threonine phosphatase PrpC, regulation of stationary phase | prpC | 0.367 | 2.89E-01 | 1.532 | 8.69E-02 | 2.538 | 5.75E-03 |
| A0A1D4R7M1 | Prophage L54a, antirepressor | | 0.246 | 3.17E-01 | 2.079 | 1.40E-02 | 2.534 | 4.02E-02 |
| A0A380ENG5 | DNA-directed RNA polymerase alpha subunit | rpoA | -0.425 | 6.47E-02 | 1.531 | 1.59E-01 | 2.502 | 1.81E-02 |
| A0A0U1MVU7 | Flavohemoprotein | YHB | 0.320 | 4.55E-02 | 1.880 | 3.15E-02 | 2.501 | 3.24E-03 |
| A0A3S4NGE7 | Citrate synthase | gltA | 0.277 | 2.72E-01 | 1.850 | 1.15E-02 | 2.481 | 5.68E-03 |
| A0A2S6D971 | ATP-dependent Clp protease ATP-binding subunit ClpC (Fragment) | | 0.440 | 1.96E-01 | 2.170 | 3.60E-02 | 2.477 | 3.85E-02 |
| A0A3A5LB65 | Putative antitoxin YezG | yezG | 0.213 | 3.70E-01 | 1.384 | 3.83E-02 | 2.472 | 2.99E-02 |
| A0A0C6EXD0 | Phage protein | | 0.211 | 2.79E-01 | 1.618 | 3.89E-02 | 2.461 | 5.12E-02 |
| A0A0C6ERM9 | Head protein | | -0.005 | 9.64E-01 | 1.449 | 4.19E-03 | 2.448 | 2.84E-02 |
| A0A380ERT1 | Elongation factor | tsf | -0.011 | 9.50E-01 | 1.605 | 8.17E-02 | 2.439 | 1.75E-02 |
| A0A0C6ELE4 | Uncharacterized protein | | 0.409 | 3.08E-01 | 1.502 | 7.56E-02 | 2.437 | 4.85E-02 |
| A0A1D4X7D9 | Triosephosphate isomerase | tpiA | 0.926 | 1.87E-01 | 2.190 | 3.58E-02 | 2.437 | 3.80E-02 |
| A0A2S6DDM8 | Valine--tRNA ligase (Fragment) | valS | -0.147 | 3.81E-01 | 1.585 | 1.22E-02 | 2.433 | 2.57E-02 |
| A0A3M8Y7T5 | Iron transporter | | 0.505 | 2.66E-01 | 1.584 | 7.64E-02 | 2.417 | 3.86E-02 |
| A0A482QPD2 | Uncharacterized protein | | 0.765 | 4.12E-02 | 2.030 | 1.97E-02 | 2.416 | 9.06E-03 |
| A0A1S6AK53 | dUTPase | | -0.002 | 9.86E-01 | 1.022 | 2.97E-03 | 2.397 | 6.79E-02 |
| A0A380DCR5 | 30S ribosomal protein S8 | rpsH | 0.005 | 9.31E-01 | 1.597 | 4.37E-03 | 2.397 | 4.94E-03 |
| A0A380ER04 | GTPase Era | bex | 0.525 | 5.05E-02 | 1.959 | 9.78E-03 | 2.389 | 6.44E-05 |
| A0A1Q8DDW1 | GntR family transcriptional regulator | | 0.723 | 2.57E-02 | 1.597 | 4.37E-02 | 2.388 | 1.31E-02 |
| A0A447Z7D6 | Pimeloyl-CoA synthase | bioW | 0.744 | 5.21E-02 | 2.068 | 2.57E-02 | 2.377 | 1.57E-02 |
| A0A077VMV9 | 2,3-bisphosphoglycerate-independent phosphoglycerate mutase | pgm | 0.135 | 6.57E-01 | 1.684 | 3.75E-02 | 2.365 | 2.64E-02 |
| A0A3M9CPR8 | Uncharacterized protein | | 0.309 | 1.57E-01 | 1.359 | 8.59E-03 | 2.362 | 5.30E-02 |
| A0A0Z1RVA9 | Enterotoxin P, phage associated | sep | 0.209 | 3.08E-01 | 1.759 | 2.20E-03 | 2.361 | 7.71E-03 |

| | | | | | | | | |
|---|---|---|---|---|---|---|---|---|
| A0A380DV53 | Uncharacterized protein | | 0.088 | 7.44E-01 | 1.740 | 6.80E-02 | 2.341 | 1.68E-02 |
| W8UY38 | Thermonuclease | nucI | 0.116 | 4.44E-01 | 1.452 | 4.42E-02 | 2.337 | 3.29E-02 |
| A0A380DGB6 | Penicillin-binding protein 3 | pbp3 | 0.309 | 2.89E-01 | 1.572 | 1.62E-01 | 2.324 | 1.12E-02 |
| A0A0C6EXE5 | Phage protein | | 0.168 | 2.37E-02 | 1.742 | 2.80E-02 | 2.310 | 5.89E-02 |
| A0A380DH40 | Succinate--CoA ligase [ADP-forming] subunit alpha | sucD | 0.241 | 3.36E-01 | 1.178 | 2.54E-02 | 2.294 | 2.99E-02 |
| A0A0U1MMD3 | Dynamin family protein | | -0.111 | 6.16E-01 | 1.113 | 1.15E-02 | 2.274 | 1.14E-02 |
| A0A0U1N0B0 | Cell division protein | ftsL | 0.066 | 6.85E-01 | 1.928 | 1.80E-02 | 2.269 | 5.66E-03 |
| A0A077UFK0 | High-affinity iron permease | efeU | 0.097 | 5.69E-01 | 1.346 | 2.36E-02 | 2.261 | 3.15E-02 |
| A0A389LVH8 | Long chain fatty acid CoA ligase | vraA | -0.060 | 7.98E-01 | 2.900 | 1.36E-02 | 2.230 | 1.21E-02 |
| Q9LC45 | N-acetylglucosaminyldiphosphoundecaprenol N-acetyl-beta-D-mannosaminyltransferase | tarA | 0.557 | 2.83E-01 | 2.626 | 1.67E-03 | 2.225 | 2.44E-03 |
| A0A380E4W1 | Phage protein OS=Staphylococcus aureus | | 0.090 | 7.29E-01 | 1.302 | 4.76E-02 | 2.224 | 3.91E-02 |
| A0A1Q8DHD8 | DNA topoisomerase 3 | topB | 0.298 | 5.33E-02 | 2.156 | 2.15E-03 | 2.217 | 1.01E-03 |
| A0A3S5DZN1 | Transcription-repair-coupling factor | mfd | 0.228 | 4.16E-01 | 2.155 | 1.33E-02 | 2.211 | 2.27E-02 |
| A0A455WI62 | Uncharacterized protein | | 0.276 | 8.68E-02 | 1.424 | 1.46E-02 | 2.206 | 4.59E-02 |
| A0A0C6E690 | Uncharacterized protein | | 0.367 | 1.49E-01 | 1.374 | 3.96E-02 | 2.206 | 3.09E-02 |
| A0A2C9TT91 | Phage head-tail adapter protein | | 0.302 | 2.80E-01 | 1.248 | 1.67E-02 | 2.203 | 6.55E-02 |
| A0A1S6BG31 | Phage nucleotide-binding protein | | 1.856 | 5.30E-02 | 2.424 | 2.90E-01 | 2.200 | 2.08E-02 |
| A0A3S4LUC3 | Alanine--tRNA ligase | alaS | 0.011 | 9.54E-01 | 1.600 | 3.80E-02 | 2.177 | 9.27E-03 |
| A0A133PXS6 | TIGR00370 family protein | | 0.171 | 4.18E-01 | 1.393 | 2.93E-02 | 2.174 | 6.47E-03 |
| A0A077UJP9 | dUTPase family protein | | 0.044 | 7.10E-01 | 1.420 | 1.15E-03 | 2.167 | 3.70E-02 |
| A0A447ZB12 | Stage 0 sporulation protein M | spo0M | 0.032 | 6.26E-01 | 1.520 | 1.04E-01 | 2.153 | 2.11E-02 |
| A0A0U1MVX0 | Ribitol-5-phosphate cytidylyltransferase | ispD | -0.093 | 3.54E-01 | 1.191 | 1.04E-01 | 2.152 | 3.38E-03 |
| G3FM61 | Triosephosphate isomerase (Fragment) | tpi | 1.033 | 3.87E-01 | 1.538 | 1.22E-01 | 2.150 | 1.54E-02 |
| A0A2X2K0V2 | D-arabino-3-hexulose 6-phosphate formaldehyde lyase | | 0.271 | 3.11E-01 | 1.916 | 1.32E-02 | 2.147 | 1.52E-02 |
| A0A1Q8DCH5 | Peptidase M24 family protein | | 0.247 | 1.09E-01 | 1.696 | 3.57E-03 | 2.144 | 1.41E-02 |
| A0A1D5ACH9 | Ribosome recycling factor | frr | 0.572 | 4.92E-01 | 2.914 | 4.97E-02 | 2.142 | 6.68E-02 |
| A0A0U1MPZ4 | YolD-like family protein | | 0.163 | 5.96E-01 | 0.867 | 1.12E-01 | 2.139 | 2.42E-02 |
| Q200E2 | SigB (Fragment) | sigB | -0.550 | 1.25E-01 | 1.244 | 6.73E-02 | 2.124 | 1.48E-03 |
| A0A380ERI8 | RLX protein | | 0.080 | 9.87E-02 | 0.652 | 3.81E-02 | 2.109 | 7.47E-02 |
| A0A2X2K1Z9 | UDP-N-acetylmuramoylpentapeptide-glycine glycyltransferase | femA | 1.879 | 7.26E-02 | 3.579 | 1.52E-02 | 2.106 | 4.31E-02 |
| A0A3S4L7Z8 | Phosphoglucosamine mutase | glmM | 0.523 | 1.38E-01 | 2.417 | 3.18E-02 | 2.103 | 8.39E-03 |
| A0A0U1MQ05 | Putative membrane protein | | 0.291 | 3.14E-02 | 1.679 | 1.50E-01 | 2.098 | 6.81E-03 |
| A0A113RL25 | Phi 11 orf33 | | 0.173 | 4.72E-01 | 1.395 | 2.44E-02 | 2.089 | 2.66E-02 |
| A0A1D4UF14 | FMN-dependent NADH-azoreductase | azoR | 0.123 | 6.94E-01 | 1.184 | 7.73E-02 | 2.083 | 2.66E-02 |
| Q59811 | Glycerol ester hydrolase | lip | -0.119 | 9.38E-03 | 2.370 | 4.05E-02 | 2.083 | 1.53E-04 |
| A0A0G2LUJ9 | Cyclic pyranopterin monophosphate synthase | moaC | 0.100 | 5.04E-02 | 1.288 | 7.32E-02 | 2.079 | 2.35E-02 |
| A0A2S6DJ10 | Peroxiredoxin (Fragment) | | 0.150 | 6.82E-01 | 1.719 | 8.09E-02 | 2.060 | 4.34E-02 |
| A0A169E8V2 | Secretory antigen | ssaA | 0.029 | 8.71E-01 | 2.308 | 2.80E-02 | 2.047 | 1.21E-02 |
| A0A2S6DBX9 | 5-(Carboxyamino)imidazole ribonucleotide synthase (Fragment) | | 0.660 | 3.06E-01 | 2.421 | 3.33E-02 | 2.047 | 2.69E-02 |
| A0A077UJL1 | Putative peptidase | pepT | 0.063 | 3.34E-01 | 1.847 | 4.60E-03 | 2.041 | 6.52E-02 |
| A0A380E9M7 | Uncharacterized protein | | 0.206 | 1.73E-01 | 1.499 | 4.34E-02 | 2.040 | 4.74E-02 |
| A0A3S4Q2A6 | GTP cyclohydrolase | folE2 | -0.086 | 5.92E-01 | 1.123 | 3.08E-02 | 2.035 | 3.45E-02 |
| A0A389V6H7 | Uncharacterized protein | | 0.178 | 3.24E-01 | 0.992 | 6.75E-03 | 2.033 | 5.49E-02 |
| A0A3S4NHK6 | Signal peptidase I | sipB | 0.212 | 3.26E-01 | 1.604 | 6.83E-02 | 2.006 | 2.16E-03 |
| A0A484VSF4 | Alkyl hydroperoxide reductase protein F | ahpF | 0.578 | 2.20E-01 | 1.648 | 9.04E-02 | 2.003 | 3.70E-02 |
| A0A0U1MJJ4 | Iron-regulated surface determinant protein B | isdB | -0.261 | 4.34E-01 | 1.915 | 1.69E-03 | 1.996 | 3.88E-02 |
| A0A167XWM0 | Deoxyadenosine kinase / Deoxyguanosine kinase | dck | 0.415 | 1.14E-01 | 1.217 | 1.07E-01 | 1.994 | 2.31E-02 |
| A0A380DCB3 | Urocanate hydratase | hutU | 0.665 | 3.19E-02 | 1.387 | 1.31E-02 | 1.987 | 5.44E-02 |

| | | | | | | | | |
|---|---|---|---|---|---|---|---|---|
| A0A0U1MW41 | Glutamate--tRNA ligase | gltX | 0.211 | 3.84E-01 | 1.421 | 5.75E-02 | 1.984 | 4.20E-02 |
| A0A2S6DR81 | Insulinase family protein | | 0.711 | 4.91E-02 | 1.650 | 8.57E-03 | 1.982 | 4.95E-02 |
| A0A131JP41 | Glycine-glycine endopeptidase | lytM | 0.069 | 2.22E-01 | 1.735 | 3.14E-04 | 1.977 | 3.20E-02 |
| A0A380DB52 | Lipase family protein | | 0.489 | 2.81E-01 | 1.690 | 1.96E-02 | 1.976 | 1.42E-02 |
| A0A0E8HDY3 | Phage capsid protein | | 0.128 | 2.12E-01 | 1.345 | 2.09E-02 | 1.976 | 2.13E-02 |
| A0A1Q8DEE5 | Ornithine aminotransferase | rocD | 0.220 | 1.80E-02 | 1.696 | 1.01E-01 | 1.973 | 2.74E-02 |
| A0A1Q8DGJ5 | ABC transporter substrate-binding protein | | -0.052 | 7.41E-01 | 1.632 | 1.02E-02 | 1.964 | 3.85E-02 |
| A0A2X2KXJ1 | Signal transduction histidine kinase | arlS | -0.028 | 8.66E-01 | 1.253 | 6.45E-02 | 1.959 | 2.97E-02 |
| A0A0E8J2S9 | Lytic transglycosylase | sceD | 0.200 | 1.74E-01 | 1.565 | 4.27E-03 | 1.955 | 2.41E-02 |
| A0A380DFQ8 | 2-succinyl-5-enolpyruvyl-6-hydroxy-3-cyclohexene-1-carboxylate synthase | menD | -0.087 | 7.66E-01 | 1.411 | 3.15E-02 | 1.946 | 6.29E-03 |
| A0A1Q8DAU2 | Aspartate aminotransferase | | 0.194 | 2.68E-01 | 1.373 | 1.89E-02 | 1.942 | 4.16E-03 |
| A0A077VN69 | DNA polymerase III delta prime subunit | holB | 0.297 | 3.32E-02 | 2.239 | 3.14E-02 | 1.941 | 8.49E-02 |
| A0A133PUH1 | Uncharacterized protein | | 0.160 | 3.36E-01 | 1.393 | 3.22E-02 | 1.937 | 5.44E-03 |
| A0A380DQT3 | Chitinase B | | -0.045 | 8.54E-01 | 1.577 | 1.55E-02 | 1.936 | 1.52E-02 |
| A0A2W3ADB2 | Phage protein | | 0.290 | 1.82E-01 | 1.756 | 2.20E-02 | 1.927 | 2.58E-02 |
| A0A380DB49 | Hydrolase | | 0.126 | 3.31E-01 | 1.000 | 6.91E-03 | 1.919 | 9.40E-03 |
| A0A0D6HWT3 | NADH-dependent flavin oxidoreductase | | 0.154 | 5.73E-02 | 1.323 | 1.90E-02 | 1.911 | 6.21E-03 |
| A0A0Z3M065 | Carboxypeptidase, M20(D) family | yxeP | 0.312 | 1.50E-01 | 1.380 | 1.33E-02 | 1.907 | 1.99E-02 |
| A0A0U1MSV6 | Putative phage infection-like protein | | -0.169 | 2.98E-01 | 1.504 | 1.15E-02 | 1.905 | 3.77E-03 |
| A0A077UF09 | PTS galactitol transporter subunit IIB | gatB | 0.111 | 5.64E-01 | 1.075 | 8.11E-02 | 1.900 | 1.94E-02 |
| A0A380E2H3 | PhiETA ORF57-like protein | | 0.279 | 2.43E-01 | 0.835 | 9.58E-02 | 1.894 | 4.12E-02 |
| A0A2W3KJ87 | Putative seryl-tRNA synthetase | | 0.132 | 4.20E-01 | 1.330 | 5.23E-03 | 1.893 | 2.98E-02 |
| A0A2X2L540 | Uncharacterized protein | | 0.521 | 1.50E-01 | 1.196 | 2.61E-03 | 1.888 | 2.66E-03 |
| A0A0C6DV27 | Uncharacterized protein | | 0.551 | 1.42E-01 | 1.366 | 3.42E-02 | 1.885 | 1.66E-02 |
| A0A455WRS3 | Uncharacterized protein | | 0.083 | 6.10E-01 | 1.293 | 4.49E-02 | 1.876 | 1.89E-02 |
| A0A2I7Y5C4 | Uncharacterized protein | | 0.237 | 2.54E-01 | 1.440 | 3.82E-02 | 1.867 | 1.36E-02 |
| A0A380E732 | Phage nucleotide-binding protein | | 0.330 | 1.46E-01 | 0.635 | 1.11E-04 | 1.858 | 2.15E-02 |
| A0A1D4R878 | 2,3-cyclic-nucleotide 2-phosphodiesterase | | 0.009 | 9.51E-01 | 1.511 | 4.09E-03 | 1.858 | 3.37E-02 |
| A0A1Q8DE28 | Nuclease SbcCD subunit C | | 0.264 | 2.26E-01 | 1.356 | 2.92E-02 | 1.856 | 1.17E-02 |
| A0A0U1MTZ1 | Uncharacterized protein | | 0.324 | 1.54E-01 | 1.520 | 9.97E-03 | 1.855 | 3.64E-02 |
| A0A380E4T3 | NH(3)-dependent NAD(+) synthetase | nadE | 0.382 | 1.73E-01 | 1.395 | 5.85E-02 | 1.853 | 2.75E-02 |
| A0A0K2D732 | Phosphate actyltransferase (Fragment) O | pta | -0.129 | 5.05E-01 | 1.131 | 5.59E-02 | 1.850 | 9.49E-03 |
| A0A133QAU9 | CHAP domain protein | | 0.880 | 1.96E-01 | 3.101 | 1.55E-02 | 1.847 | 6.49E-02 |
| A0A380DCS7 | ABC transporter permease | macB | 0.216 | 4.33E-01 | 2.401 | 5.85E-03 | 1.847 | 2.39E-02 |
| A0A133PYW1 | Serine protease | | 0.028 | 5.63E-01 | 1.643 | 2.73E-02 | 1.845 | 4.68E-02 |
| A0A380EVH6 | Phage tail length tape-measure protein | | 0.058 | 8.53E-01 | 1.129 | 1.36E-01 | 1.844 | 4.37E-02 |
| A0A077UJK9 | Acid sugar phosphatase | nagD | 0.075 | 5.42E-01 | 1.184 | 4.14E-02 | 1.843 | 3.71E-03 |
| A0A447ZDS7 | Xanthine phosphoribosyltransferase | xpt | 0.413 | 4.60E-02 | 1.973 | 7.88E-03 | 1.841 | 1.05E-01 |
| A0A1D4TY18 | Phosphoenolpyruvate-protein phosphotransferase | ptsI | 0.663 | 1.85E-01 | 1.394 | 6.05E-03 | 1.836 | 2.40E-03 |
| A0A1Q8DD87 | GMP synthase [glutamine-hydrolyzing] | guaA | -0.106 | 5.06E-01 | 0.817 | 2.82E-01 | 1.832 | 1.34E-02 |
| A0A447ZBT2 | Pyruvate carboxylase | pyc | -0.140 | 3.11E-01 | 1.004 | 1.02E-02 | 1.831 | 1.96E-02 |
| A0A1D4XZR8 | Oxidoreductase of aldo/keto reductase family, subgroup 1 | yvgN | 0.120 | 3.39E-01 | 1.960 | 7.18E-03 | 1.830 | 5.71E-02 |
| A0A1Q8DE60 | Chaperone protein | dnaK | -0.068 | 6.64E-01 | 1.072 | 1.15E-01 | 1.827 | 1.32E-02 |
| A0A3S4Q816 | Phenylalanine--tRNA ligase beta subunit | pheT | 0.207 | 4.60E-01 | 1.423 | 3.46E-02 | 1.823 | 6.44E-02 |
| A0A1Q8DCZ3 | Type II pantothenate kinase | coaW | 0.513 | 1.51E-01 | 2.073 | 1.70E-01 | 1.809 | 2.40E-02 |
| A0A0U1MZR3 | Superoxide dismutase | sodA | 0.249 | 5.67E-02 | 1.424 | 5.45E-02 | 1.799 | 9.92E-03 |
| A0A3S4MWM7 | Triosephosphate isomerase | tpiA | 0.193 | 4.18E-02 | 1.294 | 3.54E-03 | 1.783 | 2.16E-02 |
| A0A1Q8DD82 | Peptidase M24 family protein | | 0.094 | 4.39E-01 | 1.325 | 2.65E-02 | 1.778 | 4.60E-02 |

| | | | | | | | | |
|---|---|---|---|---|---|---|---|---|
| A0A380E5H5 | Aldo/keto reductase family protein | yvgN | 0.147 | 8.88E-02 | 1.199 | 1.49E-03 | 1.777 | 2.78E-02 |
| A0A090N226 | Glyoxylate reductase / Glyoxylate reductase / Hydroxypyruvate reductase | | -0.091 | 1.79E-01 | 1.052 | 7.29E-03 | 1.769 | 2.60E-03 |
| A0A1D4UX46 | Phosphoribosylaminoimidazole-succinocarboxamide synthase | purC | 0.390 | 2.22E-01 | 2.269 | 3.15E-03 | 1.766 | 6.03E-02 |
| A0A380DYB2 | Transketolase | tkt | 0.196 | 4.16E-01 | 1.371 | 2.77E-02 | 1.764 | 1.45E-02 |
| A0A380DJU3 | Serine-aspartate repeat-containing protein E | sdrE | 0.286 | 4.32E-01 | 1.660 | 1.02E-01 | 1.763 | 2.87E-02 |
| G0XY25 | Clumping factor A (Fragment) | clfA | 0.345 | 8.59E-03 | 1.537 | 1.44E-01 | 1.756 | 5.17E-02 |
| A0A077W170 | Cytosolic protein | | 0.088 | 5.04E-02 | 1.507 | 1.90E-02 | 1.753 | 2.40E-02 |
| A0A2S6DJA2 | Pyruvate kinase (Fragment) | pyk | -0.136 | 5.37E-01 | 2.159 | 2.69E-02 | 1.735 | 3.24E-02 |
| Q200T2 | S-ribosylhomocysteine lyase (Fragment) | luxS | 0.194 | 5.08E-01 | 1.216 | 4.50E-02 | 1.733 | 8.50E-03 |
| A0A0D6GN51 | Aminotransferase, class I/II | hisC | 0.065 | 6.67E-02 | 1.131 | 3.64E-02 | 1.730 | 2.36E-03 |
| A0A0U1MIC3 | D-alanine--D-alanyl carrier protein ligase | dltA | 0.030 | 3.30E-01 | 1.553 | 5.72E-04 | 1.729 | 1.45E-04 |
| A0A2Y9TRY4 | Aspartyl/glutamyl-tRNA(Asn/Gln) amidotransferase subunit B | gatB | -0.022 | 8.20E-01 | 0.546 | 8.10E-02 | 1.729 | 6.14E-03 |
| A0A077UVM0 | Alkyl hydroperoxide reductase C | ahpC | 0.168 | 1.48E-01 | 1.306 | 2.14E-03 | 1.726 | 3.38E-02 |
| A0A0U1MGB8 | Putative thioredoxin or thiol-disulfide isomerase | ydbP | 0.323 | 1.13E-01 | 1.321 | 1.05E-02 | 1.725 | 6.81E-03 |
| A0A2S6D1M4 | Ribosome biogenesis GTPase Der (Fragment) | engA | 0.046 | 7.59E-01 | 1.060 | 2.16E-02 | 1.723 | 1.11E-02 |
| A0A380DC42 | Serine--tRNA ligase | serS | 0.033 | 7.25E-01 | 1.243 | 1.34E-02 | 1.718 | 2.30E-02 |
| A0A3S4Q2L4 | Leucine--tRNA ligase | leuS | -0.030 | 5.19E-01 | 1.013 | 1.69E-02 | 1.717 | 6.25E-03 |
| A0A380DHA1 | Cysteine desulfurase | sufS | 0.263 | 3.86E-02 | 1.272 | 1.34E-02 | 1.713 | 4.30E-02 |
| A0A2S6D5K7 | Ribosomal RNA small subunit methyltransferase A | rsmA | 0.484 | 4.87E-01 | 2.025 | 8.67E-03 | 1.711 | 5.82E-02 |
| A0A3S4L4S8 | FeS assembly protein | sufD | 0.320 | 1.22E-01 | 1.311 | 7.50E-03 | 1.696 | 5.11E-03 |
| A0A166PD83 | Phi PVL ORF 22-like protein | | -0.192 | 3.05E-01 | 1.333 | 3.25E-02 | 1.693 | 1.11E-02 |
| A0A166LGR2 | Abortive infection protein | | 0.373 | 2.40E-01 | 1.527 | 6.80E-03 | 1.690 | 4.57E-03 |
| A0A1Q8DB01 | Cysteine desulfurase | | 0.106 | 6.38E-01 | 1.059 | 6.39E-03 | 1.688 | 3.04E-02 |
| A0A3S4M760 | Adenylate cyclase | yjbK | -0.015 | 9.52E-01 | 1.131 | 1.40E-01 | 1.681 | 4.90E-02 |
| A0A068DWK6 | DUF4889 domain-containing protein | | 0.196 | 4.31E-01 | 1.296 | 4.98E-02 | 1.681 | 2.34E-02 |
| A0A2W3VG08 | Glyoxalase/bleomycin resistance protein/dioxygenase superfamily protein | | 0.280 | 5.10E-02 | 1.052 | 2.54E-02 | 1.675 | 1.43E-02 |
| A0A0D6HC73 | Beta-channel forming cytolysin | lukH | 0.146 | 1.86E-01 | 1.261 | 1.52E-02 | 1.673 | 2.96E-02 |
| A0A0D6HSN6 | Heme transporter | isdB | -0.060 | 3.92E-01 | 1.637 | 1.37E-03 | 1.672 | 1.30E-03 |
| A0A1Q8DD75 | Alanine dehydrogenase | | 0.076 | 5.07E-01 | 1.148 | 2.37E-03 | 1.665 | 1.33E-02 |
| A0A3S4NHS9 | UDP-N-acetylenolpyruvoylglucosamine reductase | murB | 0.073 | 5.15E-01 | 0.958 | 1.43E-01 | 1.661 | 2.77E-02 |
| A0A2X2JVD1 | Arginine permease | lysP | 0.168 | 3.33E-01 | 1.352 | 3.07E-02 | 1.658 | 1.49E-02 |
| A0A0D6HM31 | Putative staphylococcal protein | | 0.126 | 1.13E-01 | 1.026 | 7.50E-03 | 1.657 | 5.22E-02 |
| A0A0D6HLS2 | Carboxy-terminal processing proteinase | ctpA | 0.035 | 2.84E-01 | 1.245 | 6.58E-02 | 1.647 | 1.79E-02 |
| A0A077UGV9 | GNAT family acetyltransferase | ytmI | 0.310 | 2.27E-01 | 1.091 | 1.95E-01 | 1.637 | 3.89E-02 |
| A0A380DDQ2 | Glucosaminidase | gad | -0.049 | 7.92E-01 | 1.098 | 4.90E-02 | 1.634 | 2.99E-02 |
| A0A380DX42 | UDP-N-acetylglucosamine 1-carboxyvinyltransferase | murZ | 0.134 | 3.90E-01 | 1.225 | 6.40E-04 | 1.631 | 2.33E-03 |
| A0A0D3QA44 | Conserved domain protein, putative | | 0.018 | 7.41E-01 | 1.032 | 3.38E-03 | 1.629 | 8.26E-03 |
| A0A0E7MS50 | DUF86 domain-containing protein | yutE | 0.156 | 7.58E-02 | 1.263 | 2.15E-02 | 1.626 | 3.36E-03 |
| A0A077UK57 | Tagatose 1,6-diphosphate aldolase | lacD | 0.274 | 3.61E-02 | 1.205 | 3.02E-03 | 1.623 | 3.24E-02 |
| A0A0U1MQA4 | Uncharacterized protein | | 0.196 | 1.12E-03 | 1.361 | 2.37E-02 | 1.619 | 3.30E-02 |
| C1KB86 | ClfA (Fragment) | | 0.331 | 1.38E-01 | 1.584 | 1.01E-01 | 1.612 | 8.09E-03 |
| D2J7V3 | Uncharacterized protein | | 0.122 | 6.49E-01 | 1.894 | 2.29E-02 | 1.599 | 2.75E-02 |
| A0A266CWC8 | Pyrimidine-nucleoside phosphorylase | deoA | 0.012 | 8.38E-01 | 1.179 | 6.76E-03 | 1.597 | 1.78E-02 |
| A0A389Q533 | Ser-Asp rich fibrinogen/bone sialoprotein-binding protein | sdrE | 0.141 | 4.14E-01 | 0.873 | 6.97E-03 | 1.593 | 5.80E-02 |
| A0A1Q8DEE8 | Glucose-6-phosphate isomerase | pgi | 0.187 | 2.94E-01 | 1.346 | 8.28E-03 | 1.593 | 8.72E-03 |
| A0A2S6DMU3 | Transposase | | -0.037 | 7.01E-01 | 1.410 | 2.13E-03 | 1.592 | 1.89E-03 |
| A0A1D4Y5L3 | Iron(III) dicitrate-binding protein | | 0.450 | 6.86E-02 | 2.262 | 4.08E-02 | 1.586 | 3.82E-03 |
| P0C1R5 | Galactose-6-phosphate isomerase subunit | lacA | 0.354 | 5.71E-01 | 0.922 | 1.27E-02 | 1.575 | 1.00E-01 |

| | | | | | | | | |
|---|---|---|---|---|---|---|---|---|
| A0A0D3Q935 | NERD domain-containing protein | | -0.040 | 7.15E-01 | 1.157 | 3.07E-02 | 1.568 | 2.25E-02 |
| A0A380DJ28 | Hypoxanthine phosphoribosyltransferase | hpt | -0.070 | 6.74E-02 | 0.994 | 2.13E-01 | 1.563 | 3.22E-02 |
| A0A3S4NH17 | Ribonuclease HII | rnhB | 0.205 | 4.11E-01 | 1.024 | 3.54E-02 | 1.561 | 2.11E-02 |
| A0A0E8H648 | Gamma-glutamyltransferase | ggt | -0.045 | 7.08E-01 | 0.252 | 3.50E-03 | 1.555 | 1.17E-01 |
| A0A1Q8DG35 | Aspartate--tRNA ligase | aspS | 0.271 | 9.29E-02 | 0.977 | 8.26E-03 | 1.554 | 4.46E-03 |
| A0A1L1J3H3 | Phosphoesterase | | 0.128 | 4.34E-01 | 0.813 | 3.11E-02 | 1.551 | 2.23E-02 |
| A0A077VKJ3 | Formate acetyltransferase | pflB | -0.302 | 9.96E-02 | 1.161 | 1.41E-02 | 1.551 | 5.01E-03 |
| A0A390NZS7 | Protein | grpE | 0.025 | 8.67E-01 | 0.837 | 1.21E-01 | 1.546 | 9.70E-03 |
| A0A2S6DIX3 | Cell division protein FtsZ (Fragment) | ftsZ | 0.034 | 3.98E-01 | 1.530 | 6.07E-04 | 1.545 | 5.88E-02 |
| A0A133QBU7 | Response regulator ArlR | arlR | 0.206 | 3.99E-02 | 1.068 | 4.87E-02 | 1.541 | 1.31E-03 |
| A0A0U1MR18 | Pyrimidine-nucleoside phosphorylase | pdp | 0.113 | 1.28E-01 | 1.264 | 5.06E-02 | 1.540 | 4.34E-04 |
| A0A077UPS7 | Phosphonate ABC transporter phosphate-binding periplasmic component | phnD | 0.089 | 5.22E-01 | 0.978 | 4.08E-04 | 1.536 | 1.55E-04 |
| A0A0U1MPB8 | Universal stress protein | | 0.186 | 1.40E-01 | 1.135 | 2.55E-02 | 1.534 | 2.67E-02 |
| A0A380DFN1 | Dihydroxy-acid dehydratase | ilvD | -0.249 | 6.27E-02 | 0.718 | 2.38E-02 | 1.531 | 1.54E-02 |
| A0A068A9N4 | Nickel ABC transporter substrate-binding protein | nikA | -0.033 | 6.04E-01 | 1.290 | 2.53E-03 | 1.529 | 1.77E-02 |
| X5EED1 | Phosphatidylethanolamine-binding protein | | 0.170 | 1.66E-01 | 1.249 | 6.75E-02 | 1.517 | 1.96E-03 |
| A0A484W455 | NIF3 protein | | 0.054 | 6.39E-01 | 1.146 | 2.15E-02 | 1.512 | 4.66E-03 |
| A0A1Q8DDU5 | Fructose-bisphosphate aldolase class 1 | fda | 0.121 | 3.88E-01 | 1.331 | 4.23E-02 | 1.509 | 2.12E-02 |
| A0A133Q404 | Nuclease SbcCD subunit D | sbcD | 0.116 | 1.41E-01 | 1.261 | 6.74E-04 | 1.509 | 7.99E-03 |
| A0A1Q8DDM5 | Integrase | | -0.150 | 5.24E-01 | 0.830 | 5.35E-02 | 1.507 | 3.04E-02 |
| A0A0U1MFX4 | Pseudouridine-5'-phosphate glycosidase | psuG | -0.020 | 9.26E-01 | 0.911 | 1.07E-01 | 1.504 | 2.51E-02 |
| A0A3S4L205 | Superfamily II helicase and inactivated derivatives | | 0.066 | 4.61E-01 | 0.714 | 2.84E-02 | 1.503 | 9.55E-02 |
| A0A1Q8DGA4 | Uncharacterized protein | | -0.076 | 7.81E-01 | 0.496 | 5.44E-03 | 1.502 | 1.43E-02 |
| A0A0U1MFS8 | ABC transporter, quaternary amine uptake transporter family, substrate-binding protein | | 0.090 | 6.32E-01 | 0.717 | 8.70E-02 | 1.493 | 1.85E-02 |
| R4X5D3 | Staphylocoagulase | coa | 0.333 | 6.43E-01 | 1.534 | 1.64E-02 | 1.493 | 1.73E-02 |
| A0A1Q8DFA1 | Phosphoenolpyruvate-protein phosphotransferase | | 0.018 | 8.75E-01 | 0.930 | 1.72E-02 | 1.487 | 6.99E-03 |
| A0A0U1MT10 | Nickel ABC transporter, nickel/metallophore periplasmic binding protein | nikA | -0.087 | 3.09E-01 | 1.229 | 1.03E-03 | 1.487 | 6.99E-04 |
| W8U115 | Uncharacterized protein | | 0.366 | 4.55E-02 | 0.956 | 2.59E-02 | 1.485 | 8.81E-03 |
| A0A3S4N079 | Superfamily I DNA/RNA helicase protein | | -0.063 | 7.20E-01 | 0.776 | 5.40E-04 | 1.478 | 9.12E-02 |
| W8U3Q1 | Formimidoylglutamase | hutG | 0.086 | 1.71E-01 | 1.168 | 1.32E-02 | 1.472 | 6.72E-02 |
| A0A0Y9ZDD9 | Inosine-uridine preferring nucleoside hydrolase | rihA | -0.004 | 9.74E-01 | 0.982 | 4.25E-02 | 1.472 | 3.07E-02 |
| A0A380DFE1 | Amidophosphoribosyltransferase | purF | 0.107 | 6.25E-01 | 1.140 | 8.71E-02 | 1.470 | 4.92E-02 |
| A0A1Q8DFK8 | Serine--tRNA ligase | serS | 0.737 | 8.47E-02 | 1.364 | 2.28E-01 | 1.466 | 2.51E-02 |
| A0A0D6GYJ2 | Alanine dehydrogenase | ald1 | 0.029 | 6.48E-01 | 0.961 | 4.95E-03 | 1.465 | 2.91E-03 |
| A0A0U1MWH5 | Putative exported protein | yycI | -0.070 | 3.35E-01 | 0.954 | 9.10E-03 | 1.464 | 7.67E-02 |
| A0A0U1MDD7 | L-serine dehydratase, alpha subunit | sdhA | 0.090 | 1.59E-01 | 1.023 | 3.74E-03 | 1.461 | 2.41E-02 |
| A0A0U1MC94 | S-ribosylhomocysteine lyase | luxS | 0.162 | 2.45E-01 | 1.227 | 2.92E-02 | 1.458 | 2.18E-02 |
| A0A0E8I376 | N-acetylmuramoyl-L-alanine amidase | | 0.027 | 4.48E-01 | 1.565 | 4.78E-03 | 1.457 | 1.59E-02 |
| A0A447ZDA6 | 50S ribosomal protein L11 | rplK | 0.182 | 3.71E-01 | 1.036 | 1.99E-02 | 1.457 | 2.34E-02 |
| A0A0D3Q467 | Type VII secretion protein | esaA | -0.047 | 6.64E-01 | 1.214 | 7.97E-03 | 1.455 | 1.27E-02 |
| A0A380E0T6 | Phosphoribosylformylglycinamidine synthase subunit | purQ | 0.155 | 2.13E-01 | 1.195 | 2.46E-02 | 1.454 | 2.65E-02 |
| A0A0D6HJV9 | Alpha-amylase | malA | 0.015 | 6.77E-01 | 0.945 | 5.68E-02 | 1.454 | 3.61E-02 |
| A0A2S6D350 | DUF669 domain-containing protein | | 0.054 | 6.70E-01 | 0.737 | 7.32E-02 | 1.453 | 3.03E-02 |
| A0A2L1K4G2 | Serine-aspartate repeat-containing protein D | sdrD | 0.013 | 9.23E-01 | 1.252 | 2.39E-02 | 1.448 | 4.14E-02 |
| P11175 | 6-phospho-beta-galactosidase | lacG | 0.100 | 1.14E-01 | 0.951 | 2.17E-02 | 1.442 | 3.13E-02 |
| A0A0Z4ZGY7 | Glycerol ester hydrolase Lipase | geh | 0.125 | 5.40E-01 | 1.437 | 9.94E-03 | 1.441 | 1.81E-03 |
| A0A0U1MK94 | Nuclease SbcCD subunit C | sbcC | -0.036 | 5.98E-01 | 1.290 | 2.75E-02 | 1.441 | 5.32E-02 |
| A0A2X2KBT1 | Pyridoxine biosynthesis amidotransferase | pdxT | 1.184 | 1.78E-02 | 2.002 | 1.48E-01 | 1.440 | 8.27E-03 |

| | | | | | | | | |
|---|---|---|---|---|---|---|---|---|
| A0A133PT89 | FMN-dependent NADH-azoreductase | azoR | 0.228 | 2.13E-02 | 0.743 | 4.69E-03 | 1.435 | 5.74E-02 |
| A0A0Y9RX90 | Dehydrogenase | ycjS | 0.155 | 1.79E-01 | 1.248 | 7.56E-02 | 1.433 | 3.91E-03 |
| A0A1Q8DEA4 | Probable glycine dehydrogenase (decarboxylating) subunit 2 | gcvPB | 0.647 | 9.09E-02 | 1.499 | 2.12E-01 | 1.433 | 1.78E-02 |
| A0A133PTF8 | Formate dehydrogenase OS=Staphylococcus aureus | | 0.097 | 2.63E-01 | 1.064 | 1.14E-02 | 1.423 | 3.96E-02 |
| A0A2X2KXS1 | Transcription termination factor | rho | 0.290 | 2.11E-01 | 0.999 | 7.85E-02 | 1.417 | 1.79E-02 |
| A0A380DUE7 | Serine hydroxymethyltransferase | glyA | 1.851 | 1.01E-02 | 1.478 | 5.26E-02 | 1.416 | 1.75E-02 |
| A0A077UJU0 | CsbD-like family protein | | 0.189 | 2.74E-01 | 1.259 | 1.61E-02 | 1.416 | 9.02E-03 |
| A0A2X2JWJ7 | Glutamine synthetase | glnA | -0.152 | 2.16E-01 | 0.821 | 1.58E-01 | 1.412 | 2.49E-02 |
| A0A0D3Q3Y4 | Iron-sulfur cluster repair protein | scdA | -0.051 | 6.25E-01 | 1.122 | 5.00E-03 | 1.410 | 3.96E-03 |
| A0A133Q9L3 | 7,8-dihydroneopterin aldolase | | 0.105 | 4.38E-01 | 1.193 | 1.05E-02 | 1.409 | 1.08E-02 |
| A0A1Q8DF50 | Ribosome-recycling factor | frr | 0.130 | 4.52E-01 | 0.752 | 1.27E-03 | 1.409 | 8.07E-02 |
| A0A484W167 | Surface anchored protein | isdC | -0.077 | 5.93E-01 | 1.505 | 2.65E-02 | 1.404 | 9.11E-03 |
| A0A2S6D8Z2 | Beta-ketoacyl-ACP reductase (Fragment) | | -0.193 | 5.64E-01 | 0.882 | 2.42E-01 | 1.403 | 2.96E-02 |
| A0A380EF94 | Clumping factor ClfB, fibrinogen binding protein | clfB | -0.115 | 2.96E-02 | 0.901 | 2.52E-03 | 1.402 | 4.73E-02 |
| A0A380E2R7 | Serine protease, DegP/HtrA, do-like protein | degP | 0.278 | 2.41E-02 | 1.164 | 1.22E-01 | 1.401 | 8.69E-04 |
| A0A380DQ17 | Thioesterase superfamily protein | | 0.284 | 8.96E-02 | 1.043 | 1.02E-02 | 1.399 | 4.99E-03 |
| A0A0C6E3Z3 | Phage minor capsid protein | | 0.128 | 4.02E-01 | 0.860 | 2.65E-02 | 1.397 | 2.14E-02 |
| A0A0U1MNL2 | Protein GrpE | grpE | 0.426 | 1.92E-01 | 1.398 | 1.53E-01 | 1.394 | 8.77E-03 |
| P0A017 | Dihydrofolate reductase | folA | 0.102 | 2.98E-01 | 1.028 | 2.59E-02 | 1.390 | 1.10E-02 |
| A0A077UL73 | ATP-dependent helicase/deoxyribonuclease subunit B | addB | 0.102 | 3.64E-01 | 0.883 | 1.01E-02 | 1.389 | 8.09E-03 |
| A0A090LZA4 | 3-deoxy-7-phosphoheptulonate synthase | aroX | 0.006 | 8.68E-01 | 1.182 | 1.90E-02 | 1.388 | 2.88E-02 |
| P0C1U7 | N-acetylmuramoyl-L-alanine amidase | sle1 | 0.414 | 2.10E-01 | 1.512 | 3.40E-02 | 1.383 | 1.15E-04 |
| A0A0Z0AGQ2 | Porphobilinogen deaminase | hemC | 0.224 | 9.23E-02 | 1.316 | 6.75E-02 | 1.383 | 3.06E-03 |
| A0A0U1MKI1 | Coenzyme A disulfide reductase | cdr | 0.026 | 8.56E-01 | 0.817 | 1.41E-02 | 1.377 | 4.41E-03 |
| A0A380ETU7 | Phosphoglucosamine mutase | glmM | -0.126 | 3.26E-01 | 0.981 | 3.11E-02 | 1.377 | 8.63E-03 |
| A3F6G7 | Clumping factor A (Fragment) | clfA | 0.168 | 2.35E-01 | 1.223 | 7.15E-04 | 1.374 | 1.59E-02 |
| W8UWZ2 | Thiol reductase thioredoxin | | 0.095 | 1.28E-01 | 1.118 | 3.26E-02 | 1.374 | 3.08E-02 |
| A0A1Q8DCE6 | Dihydrolipoamide dehydrogenase | | 0.824 | 1.14E-01 | 2.281 | 4.02E-03 | 1.371 | 3.96E-03 |
| A0A2S6DAG5 | Phosphopentomutase (Fragment) | | -0.001 | 9.92E-01 | 1.141 | 5.56E-03 | 1.369 | 4.84E-02 |
| A0A0U1MUK5 | Ornithine carbamoyltransferase | argF | 0.063 | 6.32E-01 | 0.922 | 4.86E-02 | 1.368 | 1.48E-02 |
| A0A077UYG7 | UPF0741 protein BN1321_150034 | | 0.054 | 7.33E-01 | 1.216 | 1.98E-02 | 1.367 | 3.29E-02 |
| A0A0D3QAU1 | Fructose-bisphosphate aldolase class 1 | fda | 0.075 | 2.25E-01 | 1.087 | 1.01E-02 | 1.366 | 1.82E-02 |
| A0A0Z0YL02 | DNA-3-methyladenine glycosidase | alkA | 0.006 | 9.13E-01 | 1.229 | 3.89E-02 | 1.366 | 9.49E-03 |
| A0A380DT36 | Dihydrolipoamide acetyltransferase component of pyruvate dehydrogenase complex | bmfBB | -0.113 | 7.29E-01 | 0.988 | 3.19E-02 | 1.364 | 9.51E-03 |
| A0A380EQJ2 | Phi PV83 orf 20-like protein | | -0.016 | 8.43E-01 | 1.042 | 2.24E-02 | 1.364 | 2.09E-02 |
| A0A0D6WDP8 | Biotin carboxyl carrier protein of acetyl-CoA carboxylase | accB | 0.186 | 3.41E-02 | 1.287 | 2.20E-02 | 1.361 | 6.66E-03 |
| X5DSE7 | 6,7-dimethyl-8-ribityllumazine synthase | ribH | 0.037 | 6.78E-01 | 1.196 | 2.20E-02 | 1.355 | 1.90E-02 |
| A0A0D3QAJ7 | Uncharacterized protein | | 0.100 | 4.31E-01 | 1.111 | 1.03E-02 | 1.348 | 8.40E-03 |
| A0A0U1MN71 | LPXTG-motif cell wall anchor domain protein | | 0.107 | 2.30E-01 | 1.540 | 5.73E-03 | 1.346 | 1.46E-02 |
| A0A0C6EL68 | Portal protein, phage associated | | 0.197 | 3.26E-01 | 0.641 | 7.62E-02 | 1.346 | 1.43E-02 |
| A0A2C9TXT2 | Nucleoside diphosphate kinase | ndk | 0.446 | 4.11E-02 | 1.175 | 1.46E-02 | 1.346 | 2.39E-02 |
| A0A0D6GE99 | LLM class flavin-dependent oxidoreductase | limB | -0.041 | 4.58E-01 | 1.049 | 7.29E-02 | 1.346 | 9.30E-04 |
| A0A0D6GIX1 | Isochorismatase | yecD | -0.086 | 7.26E-01 | 0.852 | 4.51E-02 | 1.345 | 5.05E-02 |
| A0A133QCI5 | Oligoendopeptidase F | | 0.045 | 4.60E-01 | 0.889 | 3.71E-04 | 1.345 | 2.23E-02 |
| A0A0A0R1B0 | Uncharacterized protein | | 0.080 | 6.88E-01 | 1.146 | 4.14E-02 | 1.345 | 2.18E-02 |
| A0A1Q8DCA8 | Enolase | eno | -0.068 | 7.36E-01 | 0.635 | 9.86E-02 | 1.345 | 2.38E-02 |
| P0C1S4 | Phosphoenolpyruvate carboxykinase (ATP) | pckA | 0.034 | 6.23E-02 | 0.929 | 6.35E-02 | 1.344 | 3.97E-02 |
| A0A077UI71 | Heme oxygenase (staphylobilin-producing) | isdG | -0.238 | 1.49E-01 | 1.313 | 5.58E-03 | 1.338 | 4.46E-03 |

| | | | | | | | | |
|---|---|---|---|---|---|---|---|---|
| A0A0U1MYM0 | 3-oxoacyl-[acyl-carrier-protein] synthase 2 | fabF | 0.091 | 1.30E-01 | 1.055 | 1.19E-03 | 1.336 | 8.75E-04 |
| A0A0D6HWN0 | Signal peptidase I | lepB | 0.135 | 3.12E-01 | 1.085 | 8.16E-03 | 1.334 | 3.45E-03 |
| A0A133PUJ0 | PTS system, Lactose/Cellobiose specific IIB subunit | | -0.635 | 2.18E-01 | 0.608 | 5.23E-01 | 1.325 | 7.04E-03 |
| A0A2C9THQ3 | Uncharacterized protein (Fragment) | | -0.069 | 4.52E-01 | 0.906 | 7.03E-02 | 1.323 | 3.63E-03 |
| A0A0U1MVP2 | Putative PTS system EIIBC component | ybbF | -0.145 | 4.82E-01 | 1.162 | 2.15E-01 | 1.320 | 2.09E-02 |
| A0A380EHK6 | Nitric oxide reductase activation protein | norD | -0.195 | 3.16E-01 | 0.738 | 2.92E-01 | 1.318 | 2.50E-02 |
| A0A0U1MPL3 | UDP-N-acetylmuramate--L-alanine ligase | murC | 0.601 | 1.96E-01 | 1.546 | 4.17E-02 | 1.314 | 4.38E-02 |
| A0A077V3X7 | DoxX family protein | | -0.073 | 5.79E-01 | 0.848 | 4.90E-03 | 1.312 | 2.57E-02 |
| A0A380DFJ6 | Pyruvate dehydrogenase E1 component subunit alpha | pdhA | -0.426 | 9.49E-02 | 0.334 | 2.88E-02 | 1.312 | 8.43E-02 |
| Q00626 | Beta-lactamase | | 0.068 | 3.26E-01 | 0.910 | 1.94E-02 | 1.307 | 1.05E-02 |
| A0A1Q8DGG9 | Carbamate kinase | | -0.175 | 2.62E-01 | 0.758 | 1.39E-01 | 1.304 | 2.89E-02 |
| X5DZR3 | Initiation-control protein | yabA | 0.135 | 1.42E-01 | 1.253 | 3.06E-03 | 1.303 | 1.16E-02 |
| A0A133PTB9 | Putative chlorophyll synthesis pathway protein BchC | bchC | 0.098 | 5.10E-01 | 0.991 | 5.87E-02 | 1.299 | 4.50E-02 |
| A0A484VWQ0 | Glycerol phosphate lipoteichoic acid synthase | itaS | -0.112 | 4.25E-01 | 1.178 | 9.56E-02 | 1.293 | 3.19E-04 |
| A0A077UKY6 | Putative membrane peptidase, contains TPR repeat domain | gluP | 0.180 | 1.86E-01 | 1.222 | 2.16E-02 | 1.292 | 7.93E-03 |
| A0A0U1MQS9 | Thiamine-phosphate synthase | thiE | 0.132 | 9.29E-02 | 1.090 | 1.15E-02 | 1.291 | 1.28E-02 |
| A0A1Q8DH17 | Exopolyphosphatase | | 0.405 | 1.76E-01 | 1.633 | 2.68E-02 | 1.287 | 1.74E-01 |
| A0A391F3W4 | Sensor histidine kinase | | -0.060 | 7.23E-01 | 0.973 | 3.98E-02 | 1.285 | 2.01E-02 |
| A0A2I7Y8W5 | Membrane anchored protein | | 0.092 | 1.07E-01 | 1.291 | 7.72E-03 | 1.279 | 3.49E-02 |
| A0A2S6DF63 | MSCRAMM family adhesin SdrD | sdrD | -0.206 | 4.87E-01 | 0.971 | 4.65E-02 | 1.279 | 6.53E-02 |
| A0A1Q8DEY2 | Radical SAM/CxCxxxxC motif protein YfkAB | | 0.319 | 2.43E-01 | 1.364 | 1.84E-02 | 1.277 | 2.70E-02 |
| A0A0D6FS79 | Flavohemoprotein (Hemoglobin-like protein) (Flavohemoglobin) (Nitric oxide dioxygenase) | hmp | 0.026 | 6.96E-01 | 1.018 | 1.66E-02 | 1.264 | 8.07E-03 |
| A0A133PVF3 | Thiol peroxidase | tpx | 0.173 | 2.35E-01 | 0.910 | 9.62E-03 | 1.262 | 5.24E-02 |
| A0A447ZC96 | Regulatory protein | spxA | 0.327 | 1.17E-01 | 1.335 | 1.68E-02 | 1.262 | 3.03E-02 |
| A0A380DR70 | Aconitate hydratase | citB | 0.039 | 3.67E-01 | 0.872 | 2.69E-02 | 1.260 | 1.07E-02 |
| A0A167H2Q7 | Putative glycerophosphoryl diester phosphodiesterase | glpQ | 0.067 | 6.76E-01 | 1.076 | 2.62E-02 | 1.258 | 2.50E-02 |
| A0A133Q3H1 | Isochorismatase family protein | | 0.051 | 1.92E-01 | 0.905 | 1.40E-02 | 1.257 | 9.95E-03 |
| A0A0U1MTC0 | ABC transporter family protein | | 0.002 | 9.49E-01 | 0.998 | 5.04E-02 | 1.256 | 1.63E-03 |
| A0A077VK48 | Fumarate hydratase class II | citG | 0.018 | 7.85E-01 | 0.853 | 2.88E-02 | 1.255 | 1.29E-02 |
| A0A380DSG5 | Uncharacterized protein | | 0.113 | 4.43E-01 | 0.771 | 2.77E-02 | 1.251 | 1.08E-02 |
| A0A2X2L3P1 | Thioesterase superfamily protein | entH | 0.117 | 4.50E-01 | 0.855 | 6.36E-02 | 1.250 | 2.32E-02 |
| A0A0U1MXB3 | Phosphoglycerate kinase | pgk | 0.188 | 1.11E-01 | 1.051 | 2.71E-02 | 1.248 | 4.50E-03 |
| A0A0D6G2Q2 | DASS family divalent anion:sodium (Na+) symporter | ybhI | 0.014 | 9.26E-01 | 1.038 | 4.76E-02 | 1.246 | 9.46E-03 |
| A0A3S4LCT9 | FtsW/RodA/SpoVE family cell division protein | ftsW | 0.165 | 3.09E-01 | 1.162 | 6.13E-03 | 1.244 | 2.11E-02 |
| A0A2S6D4K7 | Branched-chain-amino-acid aminotransferase | ilvE | 0.127 | 1.41E-01 | 1.017 | 8.89E-03 | 1.240 | 4.56E-02 |
| A0A2X2KYW6 | Ribonuclease D | rnd | -0.065 | 6.36E-01 | 0.956 | 2.97E-02 | 1.238 | 5.09E-02 |
| X5EEW0 | DUF2187 domain-containing protein | ykvS | 0.065 | 6.60E-01 | 1.173 | 3.63E-02 | 1.235 | 3.93E-02 |
| A0A077UN13 | Protein from nitrogen regulatory protein P-II (GLNB) family | yaaQ | 0.135 | 3.37E-01 | 0.881 | 3.66E-02 | 1.230 | 1.02E-02 |
| A0A0E8JHP5 | Aminotransferase class V-fold PLP-dependent enzyme | | -0.157 | 1.59E-01 | 0.665 | 6.30E-02 | 1.227 | 2.11E-03 |
| A0A1Q8DC93 | Phosphoglycerate kinase | pgk | 0.370 | 4.17E-01 | 0.674 | 8.92E-02 | 1.226 | 4.30E-02 |
| A0A2S6DLA2 | L-threonine dehydratase catabolic TdcB | | -0.009 | 9.71E-01 | 0.798 | 7.09E-02 | 1.223 | 3.25E-02 |
| A0A1Q8DDB2 | Alkaline-shock protein | | 0.412 | 1.29E-01 | 0.817 | 1.06E-01 | 1.220 | 6.97E-04 |
| A0A077W4L8 | ATP synthase epsilon chain | atpC | -0.005 | 9.64E-01 | 1.024 | 1.90E-02 | 1.220 | 9.51E-03 |
| A0A380DD13 | Aldehyde dehydrogenase | aldA2 | 0.270 | 3.56E-01 | 1.084 | 7.39E-02 | 1.218 | 4.22E-02 |
| A0A447Z9M1 | 3'-5' exoribonuclease YhaM | cbf1 | -0.221 | 2.90E-01 | 1.002 | 7.30E-03 | 1.216 | 1.02E-03 |
| A0A0D6GB61 | GMP synthase [glutamine-hydrolyzing] | guaA | 0.060 | 2.39E-01 | 0.963 | 4.40E-02 | 1.213 | 3.25E-03 |
| A0A0U1MMK5 | Uncharacterized protein | | -0.173 | 2.30E-01 | 0.751 | 1.18E-01 | 1.212 | 4.12E-02 |
| A0A380ETY6 | ATP-dependent DNA helicase | pcrA | -0.706 | 1.99E-04 | 0.760 | 4.10E-02 | 1.208 | 5.01E-02 |

| | | | | | | | | |
|---|---|---|---|---|---|---|---|---|
| A0A2S6DIU8 | Phosphopyruvate hydratase (Fragment) | eno | -0.142 | 4.48E-01 | 0.701 | 6.07E-02 | 1.208 | 2.70E-02 |
| A0A3S4Q1U7 | Fumarylacetoacetate hydrolase family protein | | 0.199 | 2.20E-01 | 1.122 | 2.67E-02 | 1.206 | 7.64E-02 |
| A0A2Y9TMW3 | Serine-aspartate repeat-containing protein C | sdrC | -0.070 | 2.34E-01 | 1.010 | 5.85E-03 | 1.200 | 4.68E-02 |
| W8UY26 | Catalase | katA | 0.219 | 6.66E-02 | 1.061 | 2.05E-02 | 1.197 | 6.69E-02 |
| A0A380EKC5 | Aspartyl/glutamyl-tRNA(Asn/Gln) amidotransferase subunit C | gatC | 0.020 | 7.99E-01 | 1.067 | 4.04E-02 | 1.195 | 5.89E-02 |
| A0A0U1MGP8 | Uncharacterized protein | | 0.288 | 4.39E-01 | 1.872 | 8.17E-02 | 1.191 | 1.28E-02 |
| A0A0K7KQU2 | Nitrite reductase | nasE | 0.023 | 8.99E-01 | 1.402 | 3.38E-03 | 1.189 | 4.99E-03 |
| A0A0H2AKG4 | Hypothetical phage-related protein | | 0.227 | 1.81E-01 | 0.795 | 1.40E-02 | 1.188 | 2.86E-02 |
| A0A0D6HL69 | Ribonuclease Z | rnz | 0.025 | 7.56E-01 | 1.124 | 5.52E-02 | 1.187 | 5.66E-03 |
| A0A454V6M3 | DUF2187 domain-containing protein O | | 0.010 | 8.95E-01 | 1.180 | 1.11E-02 | 1.186 | 1.05E-02 |
| A0A3S4L8M8 | Catabolite control protein A | ccpA | 0.210 | 1.52E-01 | 1.196 | 4.30E-03 | 1.180 | 2.72E-03 |
| Q93P61 | PTS system glucose-specific EIIA component | crr | -0.023 | 6.22E-01 | 0.856 | 1.00E-02 | 1.178 | 1.57E-03 |
| A0A133Q314 | S4 domain protein YaaA | yaaA | 0.267 | 1.07E-01 | 1.020 | 3.20E-02 | 1.175 | 1.65E-02 |
| A0A1Q8DBY3 | 3-oxoacyl-[acyl-carrier-protein] synthase 3 | fabH | 0.198 | 5.38E-02 | 0.871 | 1.57E-02 | 1.175 | 1.45E-02 |
| A0A077UGI5 | Salicylate hydroxylase | nagX | 0.094 | 6.02E-01 | 1.659 | 2.00E-02 | 1.174 | 5.05E-02 |
| P0C1P8 | Glutamate-1-semialdehyde 2,1-aminomutase 1 | hemL1 | 0.032 | 3.83E-01 | 0.802 | 5.17E-03 | 1.173 | 1.46E-02 |
| A0A2S6D0K7 | ThiJ/PfpI family protein | yhbO | 0.221 | 2.41E-02 | 0.943 | 5.14E-02 | 1.171 | 4.28E-02 |
| W8UX71 | Component of the preprotein translocase | yrbF | -0.361 | 7.10E-02 | 0.387 | 1.30E-01 | 1.170 | 7.54E-03 |
| A0A0D6GIE2 | Gfo/Idh/MocA family oxidoreductase | ycjS | 0.055 | 3.27E-01 | 0.952 | 5.55E-02 | 1.169 | 1.59E-02 |
| A0A2S6D0M1 | Acyl-CoA hydrolase | | 0.101 | 2.72E-01 | 0.780 | 5.68E-04 | 1.168 | 2.47E-02 |
| A0A229LY40 | DegV family protein | | -0.007 | 7.61E-01 | 0.844 | 3.19E-03 | 1.167 | 3.63E-03 |
| X5DW67 | Thiamin pyrophosphokinase | thiN | 0.105 | 1.19E-01 | 0.866 | 7.03E-02 | 1.167 | 1.10E-03 |
| A0A1Q8DEX5 | Glutamyl-tRNA(Gln) amidotransferase subunit | gatA | 0.065 | 1.10E-01 | 0.706 | 2.45E-04 | 1.164 | 8.02E-03 |
| P0A014 | 10 kDa chaperonin | groS | 0.004 | 9.63E-01 | 1.227 | 2.59E-02 | 1.160 | 4.07E-02 |
| A0A1Q8DG93 | DNA primase | | -0.250 | 2.08E-01 | 0.560 | 8.76E-02 | 1.160 | 3.56E-02 |
| A0A1Q8DFA4 | Phosphoribosylformylglycinamidine cyclo-ligase | purM | 0.036 | 7.09E-01 | 0.723 | 1.93E-02 | 1.160 | 1.16E-02 |
| A0A380ECQ0 | Metal-dependent hydrolase with rhodanese-homology domain (RHOD) | | 0.090 | 5.30E-01 | 0.755 | 2.08E-02 | 1.158 | 4.46E-02 |
| A0A0B4ND04 | L-lactate permease | lctP | -0.235 | 2.66E-01 | 0.566 | 1.00E-01 | 1.154 | 2.00E-02 |
| A0A2X2V5W4 | Protein GrpE | grpE | 0.089 | 2.10E-01 | 0.987 | 1.95E-02 | 1.152 | 2.79E-02 |
| A0A077W0Q6 | L-lactate dehydrogenase | ldh2 | -0.093 | 3.23E-01 | 0.878 | 1.59E-02 | 1.151 | 1.89E-02 |
| A0A0D6GG32 | Ribokinase | rbsK | 0.019 | 6.64E-01 | 0.621 | 7.30E-03 | 1.151 | 3.08E-02 |
| A0A2S6D9Q6 | Cysteine--tRNA ligase (Fragment) | | -0.002 | 9.89E-01 | -0.256 | 6.32E-01 | 1.149 | 1.56E-02 |
| A0A0D6GJ33 | Alpha-keto acid decarboxylase family protein | ipdC | 0.081 | 1.21E-01 | 0.969 | 1.62E-02 | 1.148 | 9.93E-04 |
| X5EMW4 | Cof-like hydrolase | | 0.387 | 2.14E-01 | 0.856 | 3.95E-02 | 1.146 | 1.85E-02 |
| A0A077ULM5 | L-2-haloalkanoic acid dehalogenase | hdl IVa | 0.056 | 1.02E-01 | 0.763 | 2.85E-03 | 1.145 | 7.73E-03 |
| D7URR5 | Cell shape-determining protein MreC | mreC | 0.167 | 4.16E-02 | 0.829 | 1.78E-03 | 1.142 | 1.88E-03 |
| A0A2X2UHP5 | Alpha-acetolactate decarboxylase | aldC | 0.138 | 1.74E-01 | 0.926 | 9.70E-02 | 1.139 | 9.76E-03 |
| A0A2S6DIS8 | Branched chain amino acid aminotransferase (Fragment) | | 0.301 | 5.55E-01 | 1.117 | 5.52E-02 | 1.137 | 3.87E-02 |
| A0A2X2JS79 | Clumping factor | clfA | -0.009 | 9.89E-01 | 1.394 | 2.23E-02 | 1.134 | 1.59E-02 |
| A0A0U1MNG8 | 6-phosphogluconate dehydrogenase, decarboxylating | gnd | -0.042 | 7.73E-01 | 0.994 | 6.14E-02 | 1.133 | 2.62E-02 |
| A0A455WXX8 | N-acetyltransferase | | -0.012 | 9.33E-01 | 1.048 | 2.96E-02 | 1.120 | 2.03E-02 |
| A0A162FHJ7 | Periplasmic binding protein | btuF | -0.438 | 4.65E-02 | 0.463 | 3.88E-01 | 1.119 | 1.44E-02 |
| A0A2C6MUA4 | Multidrug transporter | | 0.075 | 2.19E-01 | 0.714 | 9.35E-02 | 1.119 | 2.95E-02 |
| W8TZY6 | 2-oxoisovalerate dehydrogenase | pdhB | 0.046 | 3.64E-01 | 0.828 | 2.36E-02 | 1.118 | 6.11E-03 |
| A0A077VZL3 | Uncharacterized protein | | 0.022 | 7.80E-01 | 0.532 | 3.68E-02 | 1.117 | 6.04E-03 |
| A0A133PTY0 | Elastin-binding protein EbpS | ebpS | 0.026 | 6.79E-01 | 0.727 | 5.43E-02 | 1.112 | 1.67E-02 |
| A0A1D4DUI2 | Heme oxygenase (staphylobilin-producing) | isdI | 0.047 | 7.71E-01 | 1.170 | 2.91E-02 | 1.109 | 2.45E-02 |
| A0A0U1MJ30 | Phosphoesterase | | -0.027 | 8.26E-01 | 0.726 | 3.18E-02 | 1.103 | 7.58E-02 |

| | | | | | | | | |
|---|---|---|---|---|---|---|---|---|
| A0A2X2M4U4 | Glutamyl-tRNA reductase | hemA | 0.161 | 4.24E-01 | 0.458 | 2.25E-01 | 1.100 | 3.26E-02 |
| A0A380DFN4 | Transaldolase | tal | 0.150 | 2.80E-01 | 1.044 | 2.04E-02 | 1.098 | 1.74E-02 |
| A0A2S6D5M7 | 33 kDa chaperonin | hslO | 0.074 | 2.21E-01 | 0.846 | 1.03E-02 | 1.098 | 1.74E-02 |
| A0A0U1MUZ8 | Putative cytosolic protein | | 0.211 | 2.94E-01 | 1.465 | 9.65E-03 | 1.094 | 1.77E-01 |
| Q9RL69 | FmtB protein | mrp | -0.009 | 8.92E-01 | 0.871 | 3.32E-02 | 1.092 | 2.98E-02 |
| A0A0Y9XYF3 | Uncharacterized protein | | 0.184 | 2.67E-01 | 1.038 | 5.23E-03 | 1.091 | 6.34E-02 |
| A0A068A8X3 | Probable glycine dehydrogenase (decarboxylating) subunit 1 | gcvPA | 0.153 | 3.42E-02 | 0.833 | 2.20E-02 | 1.088 | 1.05E-02 |
| A0A077UI06 | Putative thiamine pyrophosphokinase | thiN | -0.159 | 3.63E-01 | 0.481 | 2.82E-01 | 1.085 | 3.52E-02 |
| A0A077UNB2 | Transcriptional regulator | pfoR | 0.477 | 8.17E-02 | 1.599 | 6.61E-03 | 1.083 | 4.50E-02 |
| A0A0U1MG43 | Uncharacterized protein | | 0.030 | 7.33E-01 | 0.554 | 1.95E-01 | 1.076 | 3.48E-02 |
| A0A115D577 | Orotidine 5'-phosphate decarboxylase | pyrF | -0.013 | 7.97E-01 | 0.926 | 3.07E-03 | 1.074 | 1.42E-02 |
| A0A0U1MGW3 | Protein DltD | dltD | -0.036 | 7.92E-01 | 0.656 | 4.33E-02 | 1.072 | 2.57E-02 |
| A0A077UJH0 | DNA-binding response regulator | bvgA | 0.082 | 6.51E-01 | 0.810 | 4.48E-02 | 1.069 | 4.11E-02 |
| A0A133Q075 | Uncharacterized protein | | 0.143 | 2.14E-01 | 0.953 | 4.73E-02 | 1.063 | 2.81E-02 |
| A0A1D4MLN0 | Potassium efflux system KefA protein / Small-conductance mechanosensitive channel | kefA | -0.043 | 6.82E-01 | 0.647 | 3.56E-02 | 1.060 | 2.07E-02 |
| A0A1K9AE69 | Adhesin | sasF | 0.132 | 3.36E-01 | 1.096 | 2.39E-02 | 1.056 | 1.17E-02 |
| A0A0D3QB03 | Membrane protein | | -0.109 | 4.22E-01 | 1.220 | 2.59E-02 | 1.052 | 8.34E-03 |
| A0A0D3Q949 | UPF0435 protein EP54_01915 | | 0.176 | 6.44E-02 | 1.194 | 1.17E-01 | 1.047 | 4.47E-02 |
| Q5TKS0 | 3-oxoacyl-[acyl-carrier-protein] synthase 2 (Fragment) | fabF | -0.070 | 7.09E-02 | 0.877 | 1.40E-02 | 1.042 | 1.31E-02 |
| W8UVN7 | 50S ribosomal protein L30 | rpmD | 0.145 | 1.71E-01 | 0.831 | 5.87E-03 | 1.039 | 2.05E-02 |
| A0A390WHA6 | Haloacid dehalogenase-like hydrolase | | 0.140 | 3.20E-01 | 0.431 | 7.83E-02 | 1.036 | 4.21E-02 |
| W8U1E8 | Pyridoxal 5'-phosphate synthase subunit | pdxT | 0.104 | 2.03E-01 | 0.736 | 3.83E-02 | 1.035 | 6.51E-03 |
| A0A0D6H3Z2 | Molybdopterin molybdenumtransferase | moeA | 0.000 | 9.96E-01 | 0.627 | 3.31E-03 | 1.033 | 9.37E-04 |
| A0A0D6GJB8 | Aldehyde dehydrogenase | aldA | 0.094 | 3.63E-01 | 0.687 | 1.88E-02 | 1.033 | 7.42E-03 |
| A0A0D3Q4H2 | DNA mismatch repair protein MutT | mutT | -0.178 | 1.33E-01 | 0.653 | 1.30E-02 | 1.030 | 2.10E-02 |
| A0A133Q3G0 | Carbamate kinase | | 0.001 | 9.81E-01 | 0.806 | 3.90E-03 | 1.028 | 1.20E-03 |
| A0A380DVR3 | Endoribonuclease | ybeY | 0.150 | 6.59E-02 | 0.730 | 4.59E-03 | 1.024 | 7.37E-02 |
| A0A2K0IFD7 | Siderophore biosynthesis protein | sbnF | -0.026 | 8.14E-01 | 1.227 | 3.04E-02 | 1.020 | 1.69E-02 |
| A0A133Q4D8 | Uncharacterized protein | | -0.077 | 2.88E-01 | 0.852 | 5.82E-04 | 1.017 | 3.16E-02 |
| A0A1D4KYI9 | Formate acetyltransferase | pflB | -0.056 | 4.45E-01 | 0.733 | 3.10E-02 | 1.017 | 4.41E-03 |
| H3K0K1 | Glutathione peroxidase | bsaA | 0.049 | 6.49E-01 | 0.759 | 2.66E-02 | 1.016 | 5.64E-02 |
| A0A0Z0HU04 | Ornithine aminotransferase | rocD2 | 0.010 | 8.56E-01 | 0.742 | 7.86E-04 | 1.016 | 1.39E-03 |
| P0A040 | Glutamine synthetase | glnA | 0.092 | 1.80E-03 | 0.949 | 5.77E-02 | 1.015 | 3.96E-05 |
| W8TUD5 | Translation initiation factor IF-1 | infA | 0.380 | 1.53E-01 | 1.007 | 2.16E-02 | 1.011 | 2.57E-02 |
| W8U006 | Phosphoribosylformylglycinamidine synthase subunit | purS | 0.071 | 4.99E-02 | 0.753 | 2.86E-02 | 1.005 | 1.13E-03 |
| A0A0D6HN74 | Transketolase | tkt | 0.026 | 6.86E-01 | 0.757 | 2.92E-02 | 1.005 | 3.52E-02 |
| A0A1Q8DEL3 | Isoleucine--tRNA ligase | ileS | 0.171 | 5.09E-01 | 0.765 | 6.91E-02 | 1.004 | 1.51E-02 |
| A0A1D4RKA7 | NADH dehydrogenase | | 1.744 | 5.08E-02 | 2.400 | 2.29E-02 | 1.002 | 2.11E-01 |
| A0A133Q2X4 | Lactonase Drp35 | | 0.280 | 2.08E-01 | 1.139 | 2.42E-02 | 0.989 | 4.15E-02 |
| A0A0U1MRF3 | Bifunctional hydroxy-methylpyrimidine kinase and hydroxy-phosphomethylpyrimidine kinase | thiD | 0.049 | 1.51E-01 | 1.070 | 7.01E-02 | 0.984 | 2.99E-02 |
| A0A380ERQ2 | Glyceraldehyde-3-phosphate dehydrogenase | gap | -0.022 | 4.20E-01 | 1.167 | 6.01E-02 | 0.954 | 1.12E-03 |
| A0A447ZJ85 | Superoxide dismutase | sodM | -0.139 | 2.29E-01 | 1.028 | 1.20E-01 | 0.951 | 3.43E-02 |
| A0A2S6D259 | ABC transporter ATP-binding protein | lolD | 0.078 | 5.89E-01 | 1.663 | 5.15E-03 | 0.946 | 7.35E-02 |
| A0A2S6D1C1 | 50S ribosomal protein L3 (Fragment) | | 0.152 | 3.45E-01 | 1.375 | 2.79E-02 | 0.846 | 1.37E-01 |
| A0A1Q8DBG9 | Uncharacterized protein | | 0.172 | 4.01E-01 | 1.830 | 4.55E-02 | 0.842 | 4.22E-02 |
| A0A133Q5N2 | Periplasmic binding protein | | -0.163 | 1.79E-01 | 1.199 | 3.06E-02 | 0.820 | 3.93E-02 |
| A0A380DQ33 | Iron aquisition yersiniabactin synthesis enzyme (Irp2) | mbtB | 2.136 | 2.63E-02 | 2.756 | 4.61E-02 | 0.701 | 2.67E-01 |
| A0A077UTV3 | Histidine-containing phosphocarrier protein of the phosphotransferase system (PTS) (HPr protein) | ptsH | 0.203 | 8.91E-02 | 1.139 | 1.17E-01 | 0.616 | 1.65E-02 |

| | | | | | | | | |
|---|---|---|---|---|---|---|---|---|
| A0A3S4NHL6 | Peptidyl-prolyl cis-trans isomerase | | 0.489 | 2.12E-01 | 1.646 | 1.10E-02 | 0.344 | 1.68E-01 |
| A0A0C5BK91 | Uncharacterized protein | | 0.637 | 1.13E-01 | 1.582 | 3.63E-02 | 0.253 | 1.60E-01 |
| A0A0U1MUY7 | tRNA modification GTPase | mnmE | 1.191 | 2.55E-01 | 2.192 | 3.09E-02 | 0.109 | 9.07E-01 |
| A0A380DS39 | RND multidrug efflux transporter Acriflavin resistance protein | swrC | 0.260 | 4.55E-01 | 1.049 | 2.29E-02 | 0.076 | 6.03E-01 |
| A0A133PVC0 | HTH-type transcriptional regulator | rot | -0.128 | 2.26E-01 | 1.094 | 4.64E-02 | -0.011 | 9.79E-01 |
| A0A133QB85 | Purine nucleoside phosphorylase DeoD-type | deoD | 0.215 | 3.44E-01 | 1.101 | 9.02E-02 | -0.233 | 9.69E-03 |
| A0A380DPN8 | Phenylalanine--tRNA ligase beta subunit | pheT | 1.286 | 2.32E-02 | 1.948 | 6.99E-03 | -0.342 | 2.61E-01 |
| A0A077UYJ9 | Putative staphylococcal protein | | 0.103 | 1.40E-01 | -1.362 | 1.09E-01 | -0.415 | 8.63E-04 |
| Q8GN53 | Holliday junction resolvase | recU | -0.029 | 2.66E-01 | -1.103 | 4.91E-02 | -0.450 | 1.07E-01 |
| A0A077VZB2 | Phosphate-specific transport system accessory protein | phoU | 0.599 | 1.03E-01 | -2.220 | 3.38E-02 | -0.494 | 7.62E-01 |
| A0A077ULD7 | Single-stranded DNA-binding protein | ssb | 0.029 | 8.88E-01 | -1.010 | 3.51E-02 | -0.596 | 1.15E-01 |
| A0A077UIR2 | General stress protein-like protein | | -0.176 | 3.41E-01 | -1.432 | 2.39E-02 | -0.815 | 4.01E-02 |
| W8UXA3 | Acetyl-CoA carboxylase biotin carboxylase subunit | accC | -0.043 | 4.56E-01 | -1.344 | 7.48E-03 | -0.880 | 8.93E-03 |
| B9UDT1 | Elongation factor Tu (Fragment) | tuf | -1.146 | 1.47E-02 | -1.234 | 6.30E-02 | -0.893 | 1.08E-01 |
| A0A145KBX7 | Excisionase | | 0.187 | 1.33E-01 | -1.166 | 4.13E-03 | -0.980 | 1.05E-02 |
| A0A0D6GWD3 | Putative lipoprotein | | -0.045 | 2.00E-01 | -1.018 | 2.42E-02 | -0.997 | 1.33E-01 |
| A0A0D6H3X1 | Molybdopterin biosynthesis MoeB protein | moeB | 0.121 | 1.95E-01 | -0.782 | 5.23E-02 | -1.003 | 5.18E-03 |
| A0A133Q1I6 | o-succinylbenzoate synthase | menC | 0.016 | 7.58E-01 | -0.696 | 3.90E-02 | -1.004 | 1.51E-03 |
| A0A447ZU22 | Dipeptidyl aminopeptidase/acylaminoacyl-peptidase | | -0.253 | 3.24E-01 | -0.502 | 3.83E-01 | -1.005 | 3.44E-02 |
| A0A455X0Y7 | Transcriptional regulator | | -0.052 | 8.46E-02 | -0.134 | 2.77E-02 | -1.006 | 2.06E-01 |
| X5E2H8 | 1,4-dihydroxy-2-naphthoyl-CoA synthase | menB | 0.060 | 3.05E-01 | -1.002 | 3.44E-02 | -1.012 | 6.36E-02 |
| A0A077ULC5 | 50S ribosomal protein L27 | rpmA | 0.009 | 7.64E-01 | -0.401 | 2.72E-01 | -1.014 | 1.04E-03 |
| A0A133PUA6 | Uncharacterized protein | | -0.232 | 2.14E-01 | -0.650 | 7.22E-02 | -1.021 | 4.92E-02 |
| A0A0D3QAJ9 | Probable malate:quinone oxidoreductase | mqo | -0.339 | 2.51E-02 | -1.108 | 5.01E-02 | -1.021 | 2.51E-03 |
| A0A0D6GRX1 | DUF4176 domain-containing protein | esaC | -0.072 | 1.76E-01 | -0.253 | 8.78E-02 | -1.025 | 1.04E-02 |
| A0A0D3Q8L6 | Tetratricopeptide domain-containing protein | yrrB | -0.292 | 2.19E-01 | -0.913 | 7.23E-03 | -1.027 | 1.16E-02 |
| A0A0U1MMS5 | Helicase loader | dnaI | -0.032 | 2.56E-01 | -0.574 | 1.51E-02 | -1.035 | 8.77E-03 |
| A0A0D3Q886 | Acetoin utilization protein | acuA | -0.066 | 1.92E-01 | -0.773 | 5.46E-03 | -1.036 | 2.30E-02 |
| A0A380DGW5 | Allophanate hydrolase subunit 2 | | -0.058 | 7.77E-03 | -0.768 | 2.61E-01 | -1.042 | 5.46E-02 |
| W8TSG6 | CDP-glycerol glycerophosphotransferase family protein | tagF | -0.148 | 1.34E-01 | -0.835 | 3.89E-03 | -1.047 | 1.43E-02 |
| A0A0D6HNK2 | Anthranilate synthase component I | trpE | 0.034 | 8.06E-01 | -0.799 | 5.67E-02 | -1.048 | 2.83E-02 |
| A0A077UG96 | Energy-coupling factor transporter ATP-binding protein | ecfA1 | -0.240 | 2.11E-01 | -0.997 | 4.85E-02 | -1.048 | 5.72E-02 |
| A0A2P7D9K6 | Protein phosphatase 2C domain-containing protein | | -0.009 | 7.51E-01 | -0.787 | 1.11E-02 | -1.052 | 4.46E-02 |
| A0A380EFH8 | NAD-specific glutamate dehydrogenase | gluD | -0.296 | 1.23E-01 | -0.462 | 1.20E-01 | -1.063 | 1.39E-02 |
| X5DRM5 | Chaperone involved in Fe-S cluster assembly | sufA | 0.100 | 1.18E-01 | -0.451 | 1.02E-02 | -1.063 | 7.07E-02 |
| A0A0B4NAE1 | UvrABC system protein A | uvrA | -0.008 | 7.48E-01 | -0.680 | 2.26E-02 | -1.068 | 1.41E-01 |
| A0A2C9TK40 | Glycosyl transferase family 1 | | -0.061 | 1.91E-01 | -0.657 | 1.67E-02 | -1.069 | 2.02E-01 |
| A0A0U1MWQ2 | Uncharacterized protein | | 0.124 | 4.64E-01 | -0.716 | 2.04E-02 | -1.069 | 2.21E-01 |
| A0A380EAW2 | Phage repressor protein | | -0.035 | 5.23E-01 | -0.783 | 8.86E-02 | -1.070 | 3.54E-03 |
| X5DZ51 | Ribosomal silencing factor RsfS | ybeB | -0.107 | 2.32E-01 | -0.252 | 5.34E-01 | -1.071 | 4.59E-02 |
| Q9KWZ5 | Uncharacterized protein ORF242 | ORF242 | -0.070 | 1.98E-01 | -1.185 | 1.39E-02 | -1.077 | 8.42E-02 |
| A0A1D5ACE0 | 3-hydroxy-3-methylglutaryl-coenzyme A reductase | | -0.475 | 1.20E-02 | -1.279 | 1.83E-01 | -1.077 | 1.31E-01 |
| A0A133PZE9 | Cytosolic protein | | -0.172 | 1.91E-01 | -1.161 | 1.05E-01 | -1.078 | 1.05E-02 |
| A0A0H2B2F3 | Uncharacterized protein | | 0.224 | 3.25E-01 | -0.901 | 2.61E-01 | -1.084 | 2.50E-02 |
| A0A133PY85 | Uncharacterized protein | | 0.072 | 3.63E-01 | -0.654 | 2.87E-02 | -1.088 | 3.11E-02 |
| A0A0U1MTR3 | Uncharacterized protein O | | -0.092 | 3.08E-01 | -0.855 | 1.93E-01 | -1.104 | 3.65E-02 |
| B3VKN2 | RNAIII-activating protein | traP | -0.143 | 6.55E-02 | -0.789 | 4.81E-02 | -1.109 | 1.64E-03 |
| A0A380EL30 | 50S ribosomal protein L31 type B | rpmE2 | 0.040 | 7.84E-01 | -1.205 | 1.53E-02 | -1.110 | 1.75E-02 |
| A0A2S6DJJ4 | Uridine kinase | udk | -0.018 | 8.97E-01 | -1.110 | 5.43E-03 | -1.111 | 2.34E-01 |

| | | | | | | | | |
|---|---|---|---|---|---|---|---|---|
| A0A133Q6U1 | Cell division inhibitor | | -0.047 | 3.45E-02 | -1.284 | 1.79E-02 | -1.116 | 9.24E-03 |
| A0A0U1MJP8 | Zinc metalloprotease | rseP | -0.013 | 9.10E-01 | -0.873 | 9.62E-02 | -1.117 | 3.93E-02 |
| A0A0U1MXE0 | Uncharacterized protein | | 0.240 | 3.16E-01 | -1.150 | 3.61E-02 | -1.122 | 7.44E-02 |
| A0A0U1MW88 | Uncharacterized protein | | -0.167 | 1.35E-01 | -1.254 | 1.33E-02 | -1.122 | 1.32E-02 |
| A0A077UIB1 | Spermidine/putrescine import ATP-binding protein | potA | -0.020 | 7.40E-01 | -1.469 | 1.40E-02 | -1.125 | 5.32E-02 |
| A0A077VZD0 | 4-phosphopantetheinyl transferase | psf-1 | 0.157 | 8.07E-02 | -0.679 | 1.68E-01 | -1.127 | 9.59E-03 |
| A0A2X2N3E8 | Thioredoxin-like oxidoreductase | | 0.019 | 6.20E-01 | -0.901 | 3.44E-03 | -1.128 | 9.61E-04 |
| A0A133PY07 | Metallo-beta-lactamase domain protein | | 0.056 | 4.68E-01 | -0.967 | 3.23E-02 | -1.136 | 6.14E-03 |
| A0A090N244 | GAF domain-containing protein | | -0.144 | 1.07E-01 | -0.741 | 3.11E-03 | -1.142 | 1.44E-03 |
| A0A0C2HXZ5 | Uncharacterized protein | | 0.054 | 7.67E-01 | -1.688 | 2.91E-01 | -1.143 | 4.41E-02 |
| A0A077UHQ2 | Endonuclease III domain protein | | 0.010 | 9.17E-01 | -0.418 | 4.76E-02 | -1.151 | 4.46E-02 |
| A0A0D3Q757 | Tryptophan synthase beta chain | trpB | -0.039 | 7.87E-01 | -0.862 | 5.60E-02 | -1.152 | 1.76E-02 |
| A0A0D6HS19 | DNA topoisomerase 1 | topA | 0.008 | 8.43E-01 | -0.460 | 2.77E-04 | -1.159 | 1.01E-01 |
| A0A133PUJ7 | ROK family protein | | -0.024 | 8.63E-01 | -1.413 | 1.60E-02 | -1.160 | 5.73E-02 |
| A0A0E7VRL0 | 3-isopropylmalate dehydrogenase | leuB | 0.062 | 7.48E-01 | -0.742 | 2.78E-01 | -1.161 | 2.96E-02 |
| A0A133Q0B1 | Glycerol kinase | glpK | 0.008 | 9.41E-01 | -1.149 | 4.58E-02 | -1.163 | 6.68E-03 |
| A0A077UQ52 | Amino acid ABC transporter, ATP-binding protein | tcyC | -0.093 | 3.10E-01 | -1.633 | 4.79E-02 | -1.165 | 3.45E-02 |
| A0A0D6GLZ1 | Phage tail protein | | 0.086 | 5.53E-01 | -0.809 | 2.26E-02 | -1.171 | 8.98E-02 |
| A0A447ZDT5 | Ribosome-binding ATPase YchF | engD | -0.189 | 3.15E-01 | -1.186 | 1.23E-01 | -1.172 | 3.69E-02 |
| A0A090LZQ0 | 50S RNA-binding protein | yqeI | 0.119 | 9.11E-02 | -0.692 | 1.37E-02 | -1.172 | 2.01E-03 |
| A0A0D6HQW1 | Proline--tRNA ligase | proS | 0.062 | 1.40E-02 | -1.182 | 2.97E-02 | -1.174 | 8.47E-03 |
| A0A169GTH9 | Glutamate-1-semialdehyde 2,1-aminomutase | gsaB | -0.030 | 1.70E-01 | -1.026 | 3.34E-02 | -1.175 | 1.67E-01 |
| A0A0D6W6I7 | NETI motif-containing protein | | -0.157 | 2.87E-01 | -0.824 | 1.45E-01 | -1.179 | 4.70E-03 |
| A0A0U1MM46 | DNA polymerase III, delta subunit | holA | -0.011 | 6.22E-01 | -1.038 | 5.02E-02 | -1.183 | 1.36E-03 |
| A0A2C9TRW4 | Iron-sulfur cluster carrier protein | apbC | -0.046 | 6.06E-01 | -1.139 | 1.08E-02 | -1.185 | 6.03E-02 |
| A0A0E7X8W5 | D-3-phosphoglycerate dehydrogenase | serA | 0.058 | 3.04E-01 | -0.554 | 2.89E-02 | -1.186 | 8.18E-02 |
| A0A077V0A1 | Thymidine kinase | tdk | 0.032 | 6.48E-01 | -0.437 | 8.67E-02 | -1.188 | 1.49E-02 |
| A0A2Y9TLN0 | 50S ribosomal protein L9 | rplI | 0.050 | 4.37E-01 | -0.351 | 3.42E-02 | -1.194 | 3.18E-02 |
| A0A162FEW9 | CBS domain-containing protein | | -0.143 | 6.78E-02 | -1.271 | 4.61E-02 | -1.194 | 4.75E-04 |
| Q53687 | FemC, factor involved in methicillin resistance / Glutamine synthetase repressor | glnR | 0.087 | 3.08E-01 | -0.579 | 2.84E-02 | -1.195 | 3.60E-02 |
| A0A0U1MHC9 | Putative cytosol aminopeptidase | pepA | -0.001 | 9.85E-01 | -0.609 | 4.29E-02 | -1.199 | 7.61E-05 |
| A0A133PZR4 | Putative ribosomal protein L7Ae-like | | -0.128 | 4.36E-01 | -0.726 | 8.70E-03 | -1.207 | 5.29E-02 |
| A0A3S4LA18 | SsrA-binding protein | smpB | 0.094 | 5.66E-01 | -1.073 | 8.76E-02 | -1.214 | 1.21E-03 |
| A0A0D3QA10 | Alanine racemase | sfaC | -0.182 | 3.81E-01 | -0.827 | 1.30E-02 | -1.220 | 1.04E-01 |
| A0A077V1B7 | Release factor glutamine methyltransferase | prmC | -0.263 | 3.58E-01 | -2.087 | 3.03E-02 | -1.221 | 4.18E-02 |
| A0A0D6HLK7 | UDP-N-acetylglucosamine--N-acetylmuramyl-(pentapeptide) pyrophosphoryl-undecaprenol N-acetylglucosamine transferase | murG | -0.199 | 1.09E-01 | -1.358 | 3.24E-02 | -1.222 | 3.65E-03 |
| A0A133PZF9 | O-acetylserine dependent cystathionine beta-synthase | mccA | 0.415 | 3.22E-02 | -0.696 | 7.68E-02 | -1.225 | 5.12E-03 |
| A0A0D3Q4U5 | Chromosome partitioning protein | parB | 0.035 | 6.41E-01 | -1.363 | 4.50E-02 | -1.241 | 2.32E-02 |
| A0A2W2TEC8 | Smooth muscle caldesmon | | -0.055 | 4.24E-02 | -0.068 | 3.65E-01 | -1.242 | 1.84E-02 |
| A0A077EW58 | Cassette chromosome recombinase B | hin | -0.070 | 4.71E-02 | -0.483 | 1.65E-02 | -1.243 | 9.00E-02 |
| A0A077UL08 | Transcriptional regulator, MarR family | | -0.124 | 1.21E-01 | -0.761 | 2.55E-02 | -1.249 | 4.30E-02 |
| A0A0D6GUN6 | Diapolycopene oxygenase | crtP | -0.051 | 4.54E-01 | -1.068 | 1.54E-02 | -1.250 | 5.61E-02 |
| A0A068DTZ1 | Autolysis response regulater | lytR | -0.052 | 6.03E-01 | -1.754 | 2.21E-03 | -1.253 | 4.59E-02 |
| A0A0U1MSI8 | Lipoprotein | | -0.212 | 2.49E-02 | -1.344 | 7.83E-03 | -1.254 | 1.31E-03 |
| A0A0U1MJX3 | Bifunctional purine biosynthesis protein | purH | 0.832 | 4.21E-02 | -1.307 | 4.55E-01 | -1.264 | 1.68E-02 |
| A0A0D6HSN8 | Small ribosomal subunit biogenesis GTPase RsgA | rsgA/engC | 0.172 | 7.81E-02 | -0.817 | 2.63E-02 | -1.266 | 4.08E-02 |
| A0A0D6HMR0 | Anthranilate phosphoribosyltransferase | trpD | 0.083 | 5.51E-01 | -1.142 | 1.39E-02 | -1.267 | 4.66E-02 |
| A0A455WGT1 | D-alanine--D-alanine ligase | ddl | -0.109 | 4.03E-02 | -0.748 | 5.27E-02 | -1.269 | 1.40E-01 |

| | | | | | | | | |
|---|---|---|---|---|---|---|---|---|
| A0A0D6HC09 | Inactive protein of metal-dependent protease family, putative molecular chaperone | tsaB/ydiC | 0.032 | 5.61E-01 | -1.634 | 3.33E-02 | -1.270 | 1.83E-02 |
| A0A380DLR8 | UPF0753 protein NCTC5664_00256 | | -0.067 | 5.76E-01 | -0.764 | 2.29E-02 | -1.270 | 1.34E-02 |
| A0A0E7RBY4 | Sulfite reductase [NADPH] flavoprotein alpha-component | cysJ | 0.040 | 3.95E-01 | -0.831 | 3.65E-02 | -1.280 | 7.38E-03 |
| A0A380E4N9 | PVL orf 52-like protein | | -0.019 | 8.27E-01 | -0.752 | 1.30E-01 | -1.282 | 3.61E-02 |
| W8U6G6 | 2'3' and 3'5' cyclic nucleotide monophosphates phosphodiesterase involved in biofilm formation | pdeB | 0.006 | 8.46E-01 | -1.397 | 3.17E-02 | -1.284 | 1.51E-02 |
| A0A133PU93 | Transcription termination factor | rho | -0.026 | 4.91E-01 | -1.104 | 3.28E-02 | -1.285 | 6.32E-02 |
| W8U368 | GTP pyrophosphokinase | ywaC | 0.008 | 7.62E-01 | -0.953 | 2.71E-03 | -1.291 | 6.84E-02 |
| W8U174 | Flavin reductase | ywqN | 0.095 | 9.13E-02 | -0.989 | 2.48E-02 | -1.292 | 4.28E-03 |
| A0A077ULL5 | LPXTG surface protein | sdrD | 0.095 | 7.31E-01 | -0.253 | 1.31E-01 | -1.298 | 2.09E-02 |
| W8U0X1 | Protein NrdI | nrdI | 0.014 | 8.51E-01 | -1.289 | 2.66E-02 | -1.299 | 2.10E-02 |
| A0A133PWU8 | Toprim domain protein | | 0.075 | 9.59E-02 | -0.846 | 3.36E-02 | -1.309 | 1.85E-01 |
| A0A2X2K0J5 | GntR family transcriptional regulator | gabR | -0.089 | 2.87E-01 | -0.734 | 2.08E-02 | -1.311 | 6.86E-03 |
| A0A1Q8DC81 | Alpha,alpha-phosphotrehalase | | -0.127 | 2.16E-01 | -1.577 | 1.07E-01 | -1.312 | 3.43E-03 |
| A0A163QRA1 | D-aminoacyl-tRNA deacylase | dtd | -0.103 | 4.41E-01 | -0.966 | 2.51E-02 | -1.316 | 3.85E-02 |
| A0A1D4ISG8 | Probable dual-specificity RNA methyltransferase | rlmN | -0.029 | 6.59E-01 | -1.058 | 2.06E-02 | -1.320 | 2.50E-02 |
| A0A380EM85 | 50S ribosomal protein L2 | rplB | -0.034 | 3.07E-01 | -0.464 | 1.35E-01 | -1.322 | 2.31E-02 |
| W8TRD5 | 50S ribosomal protein L24 | rplX | 0.035 | 6.15E-01 | -0.552 | 1.46E-01 | -1.323 | 4.63E-03 |
| A0A380DL24 | 3-ketoacyl-CoA thiolase Acetyl-CoA acetyltransferase | | -0.090 | 4.95E-01 | -1.160 | 5.06E-03 | -1.327 | 1.57E-02 |
| A0A0H2F458 | Recombinase | recT | 0.065 | 3.85E-01 | -1.545 | 1.40E-02 | -1.330 | 1.53E-02 |
| A0A0U1MUD9 | Epoxyqueuosine reductase | queH | -0.178 | 3.32E-02 | -1.800 | 6.37E-02 | -1.343 | 2.86E-02 |
| A0A133QA80 | LysR substrate binding domain protein | cidC | -0.069 | 4.60E-01 | -0.516 | 7.39E-02 | -1.347 | 1.98E-02 |
| A0A2C9TIP4 | Acyl-CoA--6-aminopenicillanic acid acyltransferase | | 0.029 | 6.81E-01 | -0.867 | 1.39E-03 | -1.348 | 7.34E-02 |
| A0A380ETQ4 | Aminopeptidase | pepS | 0.084 | 6.45E-01 | -1.646 | 4.63E-02 | -1.349 | 1.06E-01 |
| A0A0H2BKJ8 | Ribosome-binding ATPase | ychF | -0.011 | 6.30E-01 | -1.228 | 2.57E-02 | -1.351 | 3.05E-02 |
| A0A0D6HQ65 | tRNA-2-methylthio-N(6)-dimethylallyladenosine synthase | miaB | -0.234 | 1.83E-01 | -1.371 | 5.42E-02 | -1.352 | 5.90E-04 |
| A0A2X3RV25 | D-3-phosphoglycerate dehydrogenase | ghrB | 0.031 | 4.86E-01 | -0.608 | 5.23E-02 | -1.353 | 3.07E-02 |
| W8UW49 | Redox-sensing transcriptional repressor | rex | -0.091 | 2.32E-01 | -0.399 | 4.89E-02 | -1.355 | 8.10E-02 |
| W8UU09 | Phosphosugar-binding transcriptional regulator, RpiR family | ybbH | -0.015 | 8.05E-01 | -2.449 | 1.32E-01 | -1.357 | 5.05E-03 |
| A0A380E0L2 | Ribosome maturation factor | rimP | -0.099 | 4.69E-01 | -0.434 | 2.56E-01 | -1.358 | 1.98E-02 |
| A0A0U1MLP5 | Demethylmenaquinone methyltransferase | menH | -0.247 | 2.76E-01 | -1.158 | 6.47E-02 | -1.360 | 2.07E-02 |
| A0A0D6HH30 | Aminotransferase | iscS | -0.094 | 3.28E-01 | -1.609 | 2.70E-02 | -1.360 | 4.71E-03 |
| A0A0C2HX83 | Signal peptidase II | yaaT | -0.186 | 2.01E-01 | -1.283 | 3.27E-02 | -1.362 | 4.03E-02 |
| A0A068DYF6 | ATP-dependent helicase | dnaQ | -0.037 | 4.43E-01 | -0.796 | 2.03E-02 | -1.363 | 6.53E-02 |
| A0A2W2TKZ5 | Autolysin | atl | -0.190 | 1.51E-01 | -1.071 | 1.27E-02 | -1.365 | 4.39E-03 |
| A0A133Q660 | TagF domain protein | | 0.021 | 6.06E-01 | -1.441 | 3.84E-02 | -1.369 | 1.24E-02 |
| W8TUE6 | 50S ribosomal protein L13 | rplM | 0.012 | 9.01E-01 | -0.298 | 5.26E-02 | -1.371 | 1.89E-02 |
| A0A380DTU0 | DNA binding protein | | 0.149 | 7.41E-02 | -0.821 | 3.50E-03 | -1.378 | 3.78E-02 |
| W8U5R6 | Pseudouridine synthase | rluB | -0.009 | 8.79E-01 | -0.542 | 8.95E-03 | -1.380 | 3.28E-02 |
| A0A0U1MEI3 | Ribosomal RNA small subunit methyltransferase A | ksgA | -0.183 | 2.25E-01 | -0.586 | 4.28E-02 | -1.381 | 4.02E-02 |
| A0A077ULQ3 | Lipid A export ATP-binding/permease protein | msbA | -0.222 | 8.89E-02 | -1.279 | 2.27E-03 | -1.383 | 3.02E-02 |
| A0A2X2KXU1 | 3'-5' exonuclease | dinG | 0.022 | 6.49E-01 | -0.985 | 1.27E-02 | -1.383 | 2.83E-04 |
| A0A0Y9VKF8 | Carbohydrate kinase | iolC | -0.065 | 6.68E-01 | -1.594 | 8.19E-02 | -1.385 | 3.98E-03 |
| A0A0U1MKC9 | Anthranilate phosphoribosyltransferase | trpD | 0.136 | 4.66E-01 | -1.056 | 3.12E-02 | -1.386 | 1.10E-01 |
| A0A2I7Y5L8 | UPF0398 protein BTN44_05320 | | 0.016 | 7.79E-01 | -1.831 | 4.83E-03 | -1.387 | 5.62E-02 |
| A0A133QD18 | Uncharacterized protein | | -0.299 | 2.53E-01 | -0.936 | 2.74E-04 | -1.388 | 1.32E-01 |
| A0A077UHW3 | D-alanyl-D-alanine carboxypeptidase / Penicillin binding protein PBP4 | dacA | -0.230 | 8.42E-02 | -1.051 | 2.46E-02 | -1.393 | 8.69E-02 |
| A0A0D6HHP6 | Probable tRNA sulfurtransferase | thiI | 0.062 | 1.94E-01 | -1.594 | 4.68E-02 | -1.397 | 1.23E-02 |

| | | | | | | | | |
|---|---|---|---|---|---|---|---|---|
| Q936D1 | Glutaredoxin | nrdH | -0.279 | 7.21E-02 | -1.460 | 2.22E-02 | -1.402 | 2.41E-02 |
| A0A133Q440 | HAD hydrolase family | | 0.053 | 3.63E-01 | -1.080 | 1.77E-02 | -1.407 | 1.99E-02 |
| A0A0G2LUY7 | RNA-binding protein | yhgF | -0.002 | 9.03E-01 | -1.052 | 2.37E-02 | -1.409 | 1.81E-04 |
| O07318 | Outer surface binding 70kD protein (Fragment) | | -0.015 | 9.17E-01 | -0.784 | 1.76E-01 | -1.411 | 3.48E-02 |
| A0A0D6HQH3 | tRNA pseudouridine synthase B | truB | -0.061 | 1.33E-01 | -0.620 | 9.61E-02 | -1.413 | 3.51E-02 |
| A0A077UU26 | 30S ribosomal protein S3 | rpsC | -0.047 | 2.29E-01 | -1.389 | 3.16E-02 | -1.414 | 2.17E-02 |
| A0A1D5ACH6 | 30S ribosomal protein S5 | rpsE | -0.026 | 7.49E-01 | -0.840 | 6.75E-02 | -1.426 | 1.47E-02 |
| A0A0U1MJA7 | Carbamoyl-phosphate synthase small chain | pyrAA | -0.073 | 4.80E-01 | -0.721 | 2.55E-01 | -1.431 | 2.92E-02 |
| A0A1Q8DE65 | GTPase | era | 1.658 | 3.20E-02 | 0.689 | 1.60E-01 | -1.433 | 2.10E-01 |
| A0A122ZUC1 | Lysophospholipase Monoglyceride lipase putative | ytpA | -0.093 | 1.37E-01 | -1.152 | 2.01E-02 | -1.440 | 9.05E-02 |
| W8U661 | Protein VraC | vraC | 0.241 | 2.41E-01 | -0.549 | 1.82E-01 | -1.440 | 2.19E-02 |
| A0A0D6HDR3 | Lipoprotein | | -0.204 | 2.20E-01 | -0.862 | 9.05E-03 | -1.442 | 1.08E-01 |
| A0A0Z1DTC1 | Ribonuclease HIII | rnhC | 0.058 | 3.34E-01 | -0.290 | 2.38E-02 | -1.443 | 7.49E-02 |
| A0A0D6GVG9 | Methylated-DNA--protein-cysteine methyltransferase | ogt | -0.044 | 6.20E-01 | -0.695 | 3.78E-02 | -1.454 | 5.40E-02 |
| A0A3A5LQI3 | Toxin-antitoxin system, toxin component, Txe/YoeB family | | -0.029 | 6.13E-01 | -0.832 | 4.35E-03 | -1.461 | 3.33E-03 |
| A0A077ULZ0 | tRNA-dihydrouridine synthase | dus | -0.185 | 9.92E-02 | -0.963 | 1.74E-02 | -1.479 | 6.42E-02 |
| Q9KIN5 | Undecaprenyl-diphosphatase | uppP | 0.084 | 1.87E-01 | -0.905 | 1.19E-01 | -1.484 | 1.83E-02 |
| A0A077UGA7 | 50S ribosomal protein L15 | rplO | -0.097 | 2.59E-01 | -0.642 | 5.66E-02 | -1.490 | 2.94E-03 |
| A0A0K7NB37 | Ribose-phosphate pyrophosphokinase | prs | 0.006 | 9.06E-01 | -1.784 | 1.23E-03 | -1.502 | 1.37E-03 |
| A0A2X2K0Z0 | Transcription accessory protein (S1 RNA-binding domain) | | 0.657 | 3.57E-01 | 0.256 | 5.14E-01 | -1.502 | 2.74E-02 |
| A0A2X2MT76 | Phage integrase family protein | xerC | -0.080 | 1.42E-01 | -0.942 | 3.86E-02 | -1.509 | 2.11E-02 |
| W8U5N5 | Exodeoxyribonuclease 7 large subunit | xseA | -0.030 | 4.03E-01 | -1.131 | 1.01E-02 | -1.510 | 1.28E-02 |
| A0A2X2JWY3 | Colicin V production protein | | 0.070 | 2.64E-01 | -0.817 | 3.72E-03 | -1.517 | 7.08E-03 |
| A0A0S2J5U9 | Peptide methionine sulfoxide reductase regulator | msrR | -0.149 | 1.08E-01 | -1.658 | 1.07E-02 | -1.521 | 9.25E-03 |
| A0A1D4W0N2 | UPF0178 protein SAMEA1708664_00914 | | -0.047 | 6.30E-01 | -1.130 | 2.51E-03 | -1.524 | 3.77E-03 |
| A0A266CYE9 | Cytosolic protein | | 0.097 | 9.02E-02 | -0.426 | 2.18E-03 | -1.532 | 3.87E-02 |
| A0A2X2KC90 | Accessory regulator | | -0.139 | 1.97E-01 | -0.614 | 4.12E-02 | -1.532 | 5.30E-02 |
| A0A0E7Y8C1 | ACT domain protein | pheA | 0.025 | 7.12E-01 | -0.730 | 3.09E-02 | -1.534 | 4.23E-02 |
| A0A077W0Z7 | Exported protein | | -0.223 | 1.86E-02 | -1.199 | 3.00E-02 | -1.535 | 7.60E-02 |
| A0A2C9TL67 | NADH-dependent flavin oxidoreductase | namA | -0.007 | 9.38E-01 | -1.509 | 2.19E-02 | -1.536 | 7.54E-02 |
| A0A0D6HNM9 | Indole-3-glycerol phosphate synthase | trpC | 0.097 | 5.31E-01 | -0.939 | 7.20E-02 | -1.544 | 8.92E-03 |
| A0A2Y1WCJ6 | Methyltransferase domain protein | rsmC | 0.016 | 6.57E-01 | -0.789 | 1.01E-02 | -1.552 | 1.54E-02 |
| A0A484W8N6 | 50S ribosomal protein L3 | rplC | 0.144 | 5.46E-01 | -0.577 | 1.27E-01 | -1.567 | 2.12E-02 |
| A0A389PUB6 | Putative antitoxin | yezG | -0.005 | 9.80E-01 | -2.490 | 1.97E-01 | -1.570 | 2.60E-02 |
| W8TTZ1 | Glycerol uptake operon antiterminator regulatory protein | glpP | -0.011 | 8.11E-01 | -0.806 | 2.57E-03 | -1.571 | 5.10E-02 |
| A0A0E7VD74 | Uroporphyrin-III methyltransferase | nasF | -0.103 | 2.08E-01 | -1.782 | 1.17E-03 | -1.577 | 9.76E-04 |
| A0A1Q8DG16 | Bifunctional folylpolyglutamate synthase/dihydrofolate synthase | | 0.082 | 6.70E-01 | -0.580 | 1.54E-01 | -1.577 | 2.86E-02 |
| A0A133Q5C8 | Uncharacterized protein | | -0.179 | 1.22E-01 | -1.240 | 5.60E-02 | -1.578 | 4.56E-02 |
| A0A133QCN8 | Putative phosphoesterase | | 0.064 | 5.59E-01 | -0.814 | 2.11E-01 | -1.581 | 3.53E-02 |
| A0A077UX11 | Adenylate cyclase family protein | yjbK | -0.272 | 4.73E-01 | -1.754 | 1.19E-02 | -1.585 | 1.73E-01 |
| A0A133Q7E3 | Uncharacterized protein | | 0.142 | 2.08E-01 | -2.276 | 2.72E-02 | -1.590 | 5.00E-02 |
| U3U9Y8 | Partitioning protein | | -0.209 | 2.02E-01 | -0.384 | 3.50E-02 | -1.593 | 8.45E-02 |
| A0A0D6HMU8 | Nitric oxide reductase activation protein | norD | -0.028 | 1.40E-02 | -1.964 | 2.47E-02 | -1.594 | 2.84E-03 |
| A0A077UVB6 | 50S ribosomal protein L19 | rplS | -0.055 | 3.47E-01 | -0.873 | 3.22E-03 | -1.596 | 1.08E-02 |
| A0A0U1MGQ8 | Uncharacterized protein | | -0.026 | 8.35E-01 | -1.636 | 1.74E-02 | -1.603 | 9.74E-03 |
| W8UVI1 | Autolysin | atl | -0.195 | 2.37E-01 | -1.605 | 2.47E-02 | -1.603 | 8.04E-02 |
| A0A2I7Y514 | Transcriptional repressor CcpN | ccpN | -0.071 | 5.83E-01 | -2.108 | 1.40E-02 | -1.606 | 3.31E-02 |
| A0A3S4Q6L9 | Acetyltransferase | | -0.083 | 4.66E-01 | -1.516 | 2.16E-02 | -1.609 | 6.18E-02 |

| | | | | | | | | |
|---|---|---|---|---|---|---|---|---|
| A0A0U1MF66 | Manganese ABC transporter (ATP-binding protein) | mntB | 0.112 | 5.00E-01 | -1.913 | 1.39E-02 | -1.610 | 1.58E-02 |
| A0A077VLD5 | 30S ribosomal protein S17 | rpsQ | 0.022 | 7.28E-01 | -0.573 | 3.47E-02 | -1.616 | 6.12E-02 |
| A0A133QD78 | Mannose-6-phosphate isomerase | | -0.262 | 4.11E-01 | -1.638 | 6.62E-03 | -1.618 | 7.04E-03 |
| A0A077UXQ0 | UTP--glucose-1-phosphate uridylyltransferase | gtaB | -0.035 | 4.75E-01 | -1.608 | 2.24E-03 | -1.620 | 1.19E-02 |
| A0A2W3ID47 | Acetyl-coenzyme A carboxylase carboxyl transferase subunit beta | accD | 0.002 | 9.70E-01 | -0.768 | 5.27E-02 | -1.625 | 1.46E-02 |
| A0A2X2JYL6 | GMP synthase (Glutamine-hydrolyzing) | guaA | -0.471 | 1.30E-01 | -1.745 | 5.27E-03 | -1.630 | 1.03E-01 |
| A0A0D6HMW3 | CCA-adding enzyme | papS | -0.141 | 4.81E-02 | -2.140 | 1.62E-01 | -1.632 | 9.28E-03 |
| A0A133QCQ2 | Choline ABC transporter, ATP-binding protein OpuBA | | -0.036 | 6.15E-01 | -1.776 | 9.71E-02 | -1.637 | 1.81E-02 |
| X5E067 | Acetate kinase | ackA | -0.173 | 1.02E-01 | -0.801 | 2.25E-01 | -1.640 | 7.13E-03 |
| O05155 | Glycerol-3-phosphate cytidylyltransferase | tagD | -0.022 | 7.55E-01 | -1.018 | 1.49E-02 | -1.642 | 4.21E-02 |
| A0A077UGV8 | 50S ribosomal protein L36 | rpmJ | -0.085 | 5.21E-01 | -0.729 | 7.06E-02 | -1.659 | 2.78E-02 |
| A0A2S6D518 | tRNA-specific 2-thiouridylase | mnmA | 0.062 | 7.80E-02 | -2.033 | 3.11E-04 | -1.660 | 5.53E-02 |
| K0A6J6 | FemA (Fragment) | femA | -0.073 | 2.51E-01 | -1.816 | 1.17E-01 | -1.667 | 3.92E-02 |
| A0A077VK60 | ABC transporter ATP-binding protein | ecsA | -0.377 | 3.22E-01 | -1.986 | 1.19E-01 | -1.668 | 2.86E-02 |
| A0A077UHD1 | Fic/DOC family protein | | 0.082 | 5.04E-01 | -1.000 | 2.01E-02 | -1.677 | 3.97E-03 |
| X5EHA8 | Uncharacterized protein | | -0.131 | 3.79E-01 | -1.049 | 1.43E-02 | -1.684 | 1.72E-02 |
| A0A0H2K0A0 | 30S ribosomal protein S10 | rpsJ | -0.064 | 2.60E-01 | -1.460 | 7.06E-02 | -1.685 | 6.84E-03 |
| A0A2S6D8D3 | Asp-tRNA(Asn)/Glu-tRNA(Gln) amidotransferase GatCAB subunit B (Fragment) | | -0.447 | 1.74E-02 | -1.727 | 3.86E-02 | -1.687 | 3.17E-03 |
| A0A2X3PLR9 | LSU m5C1962 methyltransferase | rlmI | -0.117 | 6.27E-01 | -1.020 | 4.21E-02 | -1.688 | 8.42E-02 |
| A0A0D6GN06 | DNA-binding protein | | 0.082 | 7.41E-01 | -1.722 | 5.25E-02 | -1.692 | 3.17E-02 |
| A0A068A921 | Nitrate reductase | narH | -0.101 | 1.53E-01 | -1.273 | 1.47E-02 | -1.693 | 7.19E-03 |
| W8TRB5 | Probable cell division protein | whiA | 0.082 | 4.65E-01 | -1.055 | 1.06E-02 | -1.694 | 2.23E-03 |
| W8TUC9 | 30S ribosomal protein S5 | rpsE | -0.018 | 3.25E-01 | -1.067 | 6.58E-02 | -1.696 | 1.04E-02 |
| A0A077V2C0 | NAD(FAD)-utilizing dehydrogenase | | -0.045 | 2.27E-01 | -0.966 | 1.11E-02 | -1.697 | 7.45E-03 |
| W8UX38 | Glyceraldehyde-3-phosphate dehydrogenase | gapA2 | 0.055 | 3.66E-01 | -1.407 | 3.92E-02 | -1.702 | 3.66E-03 |
| W8U3W0 | 50S ribosomal protein L3 | rplC | -0.129 | 8.03E-02 | -0.623 | 1.74E-01 | -1.713 | 2.13E-02 |
| A0A380EJB9 | Glycerol-3-phosphate dehydrogenase | glpD | -0.037 | 4.49E-01 | -0.880 | 7.00E-02 | -1.714 | 7.98E-03 |
| A0A133Q094 | ACT domain protein | | 0.042 | 1.72E-01 | -1.612 | 4.42E-02 | -1.718 | 5.55E-02 |
| A0A0D3Q6G5 | Cell division protein | zapA | 0.082 | 3.63E-01 | -1.560 | 4.30E-03 | -1.721 | 7.41E-03 |
| A0A0D3QB96 | PadR family transcriptional regulator | | 0.021 | 7.66E-01 | -0.650 | 5.37E-02 | -1.722 | 4.23E-02 |
| A0A0U1N0K2 | Riboflavin biosynthesis protein | ribC | 0.042 | 5.14E-01 | -1.203 | 2.29E-02 | -1.725 | 8.32E-02 |
| A0A2C9TQD3 | Bifunctional (P)ppGpp synthetase/guanosine-3',5'-bis(Diphosphate) 3'-pyrophosphohydrolase | | -0.046 | 3.80E-01 | -1.870 | 9.53E-03 | -1.728 | 4.18E-03 |
| A0A068DTK5 | Alpha-helical coiled-coil protein | srpF | 0.220 | 2.76E-01 | -1.530 | 8.85E-03 | -1.729 | 8.40E-03 |
| P0C5Z9 | UPF0413 protein (Fragment) | | 0.052 | 2.44E-01 | -1.439 | 4.35E-02 | -1.730 | 3.86E-02 |
| A0A390RWB3 | Cys-tRNA(Pro)/Cys-tRNA(Cys) deacylase | ybaK | -0.071 | 1.89E-01 | -0.843 | 1.51E-03 | -1.733 | 2.32E-02 |
| A0A0Z3MW82 | D-isomer specific 2-hydroxyacid dehydrogenase family protein O | ldhD | -0.031 | 3.56E-01 | -1.969 | 7.80E-02 | -1.735 | 2.08E-02 |
| A0A0D6GQW4 | Transcription factor | fapR | -0.045 | 2.18E-01 | -1.455 | 2.37E-02 | -1.736 | 3.18E-02 |
| A0A090LWE8 | Bacterial globin-like protein | yjbI | -0.182 | 4.59E-01 | -1.807 | 2.79E-02 | -1.742 | 3.32E-02 |
| X5EP39 | 2-amino-3-ketobutyrate CoA ligase | | -0.219 | 2.43E-01 | -0.697 | 8.39E-02 | -1.742 | 4.44E-02 |
| A0A2S6DKI3 | 30S ribosomal protein S7 | rpsG | -0.355 | 1.88E-01 | -1.222 | 6.30E-02 | -1.744 | 1.02E-02 |
| A0A2I7Y9K2 | Threonylcarbamoyl-AMP synthase | | 0.063 | 5.07E-01 | -1.715 | 1.05E-02 | -1.758 | 1.16E-02 |
| A0A0D6HV28 | Aminotransferase | patA | -0.088 | 2.88E-01 | -1.826 | 8.13E-02 | -1.763 | 3.47E-03 |
| A0A2X2UMS7 | Glycosyl transferase | | -0.083 | 1.53E-01 | -1.049 | 1.06E-02 | -1.764 | 1.38E-02 |
| Q8KYA6 | Glyoxalase-like domain protein | | -0.133 | 4.04E-02 | -1.666 | 6.21E-02 | -1.764 | 3.06E-03 |
| A0A0D6HKV2 | DNA primase | dnaG | -0.083 | 2.10E-01 | -1.673 | 3.24E-02 | -1.767 | 7.81E-04 |
| A0A0D6GQD4 | Mannose-6-phosphate isomerase | pmi/manA | -0.195 | 1.96E-01 | -1.000 | 7.13E-04 | -1.769 | 5.55E-02 |
| A0A133Q037 | DNA polymerase III PolC-type | polC | -0.060 | 2.50E-01 | -2.019 | 1.87E-02 | -1.772 | 3.09E-03 |
| A0A2S6D2I5 | Betaine aldehyde dehydrogenase | betB | -0.024 | 7.20E-01 | -1.870 | 9.21E-03 | -1.774 | 1.06E-02 |

| | | | | | | | | |
|---|---|---|---|---|---|---|---|---|
| P68863 | DNA replication and repair protein | recF | -0.024 | 2.79E-01 | -1.937 | 4.32E-02 | -1.777 | 1.08E-02 |
| O05337 | Putative pyruvate, phosphate dikinase regulatory protein | | 0.249 | 1.34E-01 | -1.884 | 4.40E-02 | -1.787 | 8.57E-02 |
| A0A133Q5T3 | ABC transporter, ATP-binding protein | | -0.033 | 3.84E-01 | -2.341 | 4.49E-02 | -1.795 | 1.25E-02 |
| A0A166Q2N9 | Ribosome biogenesis GTPase A | rbgA | -0.113 | 8.37E-02 | -1.293 | 2.25E-02 | -1.798 | 7.01E-04 |
| A0A0E7DVG7 | ABC transporter | yjjK | 0.111 | 1.08E-01 | -1.977 | 9.78E-04 | -1.808 | 5.03E-04 |
| A0A077VHU6 | 30S ribosomal protein S7 | rpsG | -0.010 | 8.72E-01 | -0.999 | 4.32E-02 | -1.812 | 2.61E-03 |
| A0A133Q127 | HTH-type transcriptional regulator | | 0.018 | 7.27E-01 | -0.925 | 1.94E-02 | -1.817 | 9.17E-02 |
| A0A0D6H5T3 | tRNA pseudouridine synthase A | truA | 0.004 | 9.71E-01 | -1.410 | 2.48E-02 | -1.829 | 5.93E-03 |
| A0A0D6HTX7 | Class I SAM-dependent rRNA methyltransferase | rlmI | -0.118 | 1.95E-01 | -1.607 | 1.11E-02 | -1.833 | 1.67E-03 |
| A0A2W3L6J0 | CobB/CobQ-like glutamine amidotransferase domain-containing protein | | -0.306 | 2.08E-01 | -1.702 | 5.52E-02 | -1.834 | 1.05E-02 |
| A0A2W2JFP2 | DNA-binding protein | | 0.014 | 9.27E-01 | -0.850 | 9.19E-02 | -1.836 | 1.92E-02 |
| A0A133PYC3 | 30S ribosomal protein S19 | rpsS | -0.079 | 4.37E-01 | -0.973 | 1.52E-01 | -1.838 | 6.74E-03 |
| A0A077UNE0 | 50S ribosomal protein L1 | rplA | -0.226 | 7.11E-02 | -0.899 | 1.42E-01 | -1.838 | 2.44E-02 |
| W8TVK2 | 30S ribosomal protein S4 | rpsD | -0.096 | 2.70E-02 | -0.966 | 4.98E-02 | -1.841 | 4.21E-02 |
| A0A077UDU1 | 30S ribosomal protein S2 | rpsB | -0.022 | 7.35E-01 | -0.913 | 3.48E-02 | -1.845 | 4.08E-02 |
| A0A077UKF9 | 50S ribosomal protein L22 | rplV | -0.056 | 3.39E-01 | -1.167 | 1.32E-01 | -1.853 | 1.67E-02 |
| A0A0U1MVF7 | Putative thiamine pyrophosphate-containing protein | ydaP | 0.043 | 4.32E-01 | -1.856 | 4.37E-02 | -1.877 | 1.26E-02 |
| A0A229LYM8 | MarR family transcriptional regulator | | 0.014 | 6.90E-01 | -0.729 | 7.96E-02 | -1.878 | 1.01E-03 |
| A0A0D6HT57 | Glycerophosphodiester phosphodiesterase | ugpQ | 0.060 | 5.28E-01 | -1.311 | 2.13E-02 | -1.880 | 6.93E-03 |
| A0A1D4UAJ8 | Ribonuclease HI | rnhA | -0.137 | 4.88E-01 | -1.483 | 2.39E-02 | -1.895 | 5.80E-05 |
| A0A2S6D5H7 | Protein map | | -0.223 | 3.74E-01 | -2.161 | 9.30E-02 | -1.898 | 4.18E-02 |
| A0A0D3Q9P6 | ATP-dependent Clp protease ATP-binding subunit | clpB | -0.028 | 6.87E-01 | -1.673 | 3.04E-03 | -1.900 | 8.57E-02 |
| A0A0D6H5T5 | DNA topoisomerase 3 | topB | -0.078 | 2.35E-01 | -1.204 | 2.61E-02 | -1.902 | 1.95E-02 |
| A0A1D4SF96 | Acetyltransferase (GNAT) family protein | | -0.008 | 8.37E-01 | -1.436 | 8.09E-02 | -1.905 | 3.81E-04 |
| A0A122NGZ2 | O-methyltransferase family protein | | -0.091 | 3.05E-01 | -2.340 | 1.05E-02 | -1.911 | 1.49E-03 |
| A0A2S6D714 | Asparagine--tRNA ligase (Fragment) | asnC | 0.752 | 1.99E-01 | -1.926 | 9.08E-03 | -1.914 | 2.28E-02 |
| A0A455JRS7 | TIGR01741 family protein | | -0.158 | 2.83E-02 | -1.952 | 4.33E-02 | -1.915 | 4.10E-02 |
| W8TRC7 | 50S ribosomal protein L4 | rplD | -0.255 | 1.93E-02 | -1.038 | 1.61E-01 | -1.920 | 2.09E-02 |
| A0A444VF20 | Uncharacterized protein | | -0.082 | 3.28E-01 | -1.200 | 2.15E-02 | -1.921 | 7.17E-02 |
| A0A077UMN7 | Hemolysin-like protein containing CBS domains | | -0.240 | 2.64E-02 | -2.116 | 2.97E-02 | -1.924 | 3.67E-02 |
| A0A0D6GSR0 | tRNA dimethylallyltransferase | miaA | 0.107 | 1.72E-01 | -1.123 | 3.17E-03 | -1.924 | 7.46E-02 |
| A0A133Q104 | Pantothenate synthetase | panC | -0.021 | 7.93E-01 | -1.783 | 2.81E-02 | -1.925 | 4.18E-02 |
| A0A380EFY1 | Bifunctional purine biosynthesis protein | purH | 0.135 | 5.21E-02 | -2.175 | 6.19E-02 | -1.925 | 4.53E-02 |
| A0A1D4TM68 | Dehydrogenase | gdhII | 0.336 | 1.50E-01 | -1.430 | 1.29E-02 | -1.930 | 9.02E-02 |
| A0A0D6HX24 | Pseudouridine synthase | yjbO | -0.024 | 9.29E-01 | -1.342 | 4.65E-03 | -1.930 | 1.05E-02 |
| A0A0D3Q6F6 | Extracellular fibrinogen-binding protein | fib | -0.079 | 6.21E-01 | -1.635 | 1.02E-02 | -1.938 | 5.49E-02 |
| W8UT65 | Peptidyl-tRNA hydrolase | pth | 0.054 | 5.16E-01 | -1.519 | 7.99E-03 | -1.942 | 5.86E-02 |
| A0A0H2C2Z4 | Bacterial protein of uncharacterized function (DUF951) | | -0.144 | 2.44E-01 | -1.292 | 3.87E-03 | -1.947 | 2.81E-03 |
| A0A0U1MWX3 | Putative hemin import ATP-binding protein | hrtA | -0.040 | 5.19E-01 | -2.251 | 2.65E-02 | -1.948 | 2.88E-02 |
| A0A1D4PWR6 | Aldehyde dehydrogenase | | 0.109 | 6.43E-03 | -2.238 | 1.80E-02 | -1.951 | 1.53E-03 |
| A0A2X2KSQ0 | 30S ribosomal protein S10 | rpsJ | -0.173 | 2.55E-01 | -1.931 | 7.20E-02 | -1.953 | 1.25E-02 |
| A0A0D3Q8K8 | Acetyl-coenzyme A carboxylase carboxyl transferase subunit alpha | accA | -0.075 | 1.89E-01 | -1.148 | 3.38E-02 | -1.970 | 1.14E-02 |
| A0A0U1MFE0 | Putative acetyl-CoA C-acetyltransferase | yhfS | -0.115 | 2.31E-01 | -1.661 | 6.24E-03 | -1.974 | 7.61E-03 |
| A0A1D4Q6A4 | Urease accessory protein | ureD | 0.009 | 7.21E-01 | -1.211 | 1.27E-02 | -1.975 | 4.86E-02 |
| A0A133Q045 | Translation initiation factor IF-2 | infB | -0.056 | 2.96E-02 | -2.081 | 4.02E-02 | -1.979 | 2.13E-03 |
| A0A168QXT6 | DEAD-box ATP-dependent RNA helicase | cshA | -0.037 | 5.73E-02 | -1.808 | 7.33E-03 | -1.980 | 3.39E-05 |
| W8UY96 | Ribonuclease J | rnjB | -0.230 | 1.72E-01 | -1.262 | 7.10E-03 | -1.986 | 1.09E-03 |
| A0A077UPQ3 | Uncharacterized protein | | -0.212 | 9.12E-02 | -2.382 | 1.60E-01 | -1.988 | 1.49E-02 |

| | | | | | | | | |
|---|---|---|---|---|---|---|---|---|
| A0A077V4G0 | 50S ribosomal protein L16 | rplP | 0.087 | 3.01E-01 | -0.942 | 2.89E-03 | -1.991 | 2.21E-02 |
| A0A0D6HG79 | Helicase | dnaB | -0.017 | 7.85E-01 | -2.128 | 9.88E-04 | -1.999 | 1.19E-02 |
| A0A133PU41 | Asparagine--tRNA ligase | asnS | 0.044 | 1.67E-01 | -2.074 | 9.44E-02 | -2.005 | 3.54E-02 |
| A0A077URJ8 | 50S ribosomal protein L28 | rpmB | -0.139 | 1.62E-01 | -1.145 | 5.30E-02 | -2.014 | 5.24E-04 |
| A0A1D4L9H6 | Fe-S oxidoreductase | | -0.047 | 4.85E-01 | -1.418 | 3.47E-03 | -2.014 | 1.04E-03 |
| A0A077UJF1 | Iron-sulfur cluster biosynthesis family protein | | -0.219 | 2.07E-01 | -3.327 | 1.28E-01 | -2.020 | 6.31E-03 |
| A0A3S4LIS0 | Transcription factor | | -0.118 | 2.54E-01 | -0.567 | 1.39E-02 | -2.020 | 5.72E-02 |
| A0A0D3QB56 | UDP-N-acetylglucosamine--peptide N-acetylglucosaminyltransferase stabilizing protein | gtfB | -0.188 | 3.97E-02 | -1.958 | 6.12E-02 | -2.022 | 2.30E-02 |
| A0A133Q1I5 | Nucleoside triphosphatase YtkD | ytkD | -0.049 | 2.94E-01 | -1.361 | 6.38E-05 | -2.028 | 4.26E-02 |
| A0A0U1MNV9 | GTPase involved in ribosome 30S assembly | yqeH | 0.060 | 5.46E-01 | -1.029 | 6.99E-03 | -2.029 | 3.73E-04 |
| A0A447ZRP4 | Anthranilate phosphoribosyltransferase | trpD2 | -0.183 | 5.16E-01 | -1.729 | 1.38E-01 | -2.031 | 3.56E-02 |
| A0A077UI74 | Peptide deformylase | def | -0.066 | 2.51E-02 | -1.245 | 4.37E-02 | -2.036 | 2.78E-02 |
| A0A077UM51 | ABC transporter ATP-binding protein | ykoD | -0.027 | 2.32E-01 | -1.513 | 7.66E-04 | -2.037 | 1.51E-01 |
| A0A0D6GGT8 | ABC transporter ATP-binding protein | oppF | 0.044 | 7.35E-01 | -2.045 | 2.58E-03 | -2.039 | 2.54E-03 |
| G5ELA5 | Pyridine nucleotide-disulfide oxidoreductase | | 0.056 | 2.19E-01 | -2.267 | 1.87E-02 | -2.042 | 4.18E-02 |
| A0A0D6H7H2 | ATP-dependent RecD-like DNA helicase | recD | -0.036 | 6.12E-01 | -1.759 | 6.55E-02 | -2.043 | 4.36E-02 |
| Q9KWZ9 | Pseudouridine synthase | ORF231 | 0.022 | 7.39E-01 | -1.734 | 1.44E-02 | -2.045 | 3.58E-02 |
| W8TU12 | DNA-binding protein | ylxR | -0.191 | 1.22E-01 | -0.906 | 5.94E-02 | -2.059 | 2.73E-03 |
| H9BRP0 | Putative phosphoserine phosphatase | rsbU | -0.049 | 3.85E-02 | -1.911 | 4.97E-02 | -2.064 | 5.25E-02 |
| A0A077UG91 | 50S ribosomal protein L17 | rplQ | -0.139 | 1.52E-02 | -1.259 | 1.54E-01 | -2.064 | 4.05E-03 |
| A0A1Q8DCS8 | DNA-directed RNA polymerase subunit beta | rpoB | -0.575 | 3.15E-01 | -1.643 | 1.05E-02 | -2.064 | 6.89E-02 |
| A0A077UJD8 | UDP-N-acetylmuramoylpentapeptide-glycine glycyltransferase | femA | 0.336 | 4.22E-01 | -0.913 | 1.35E-01 | -2.069 | 4.47E-02 |
| A0A133Q678 | Ferredoxin--NADP reductase | | 0.106 | 2.80E-01 | -1.892 | 1.85E-03 | -2.071 | 1.16E-03 |
| A0A133Q0Y4 | Dehydrosqualene synthase | | 0.003 | 9.76E-01 | -1.814 | 1.22E-02 | -2.072 | 3.36E-03 |
| O85142 | ArsR family transcriptional regulator | rzcA | 0.068 | 1.77E-01 | -1.114 | 1.23E-01 | -2.072 | 1.99E-02 |
| W8TP29 | Ribosomal RNA small subunit methyltransferase I | rsmI | 0.048 | 5.64E-01 | -1.852 | 3.86E-02 | -2.074 | 2.14E-02 |
| A0A0U1ML69 | DNA polymerase III PolC-type | polC | 0.117 | 3.52E-01 | -1.711 | 1.55E-02 | -2.074 | 5.80E-02 |
| A0A380E2U2 | Isocitrate dehydrogenase | citC | -0.223 | 4.27E-02 | -1.215 | 1.95E-01 | -2.077 | 1.62E-01 |
| W8TUB4 | 50S ribosomal protein L23 | rplW | -0.033 | 3.87E-01 | -0.991 | 5.26E-02 | -2.094 | 2.46E-02 |
| A0A0U1MMK6 | ATP-dependent Clp protease ATP-binding subunit | clpX | 0.003 | 9.58E-01 | -1.594 | 9.04E-02 | -2.103 | 4.92E-02 |
| A0A077UHS1 | 30S ribosomal protein S21 | rpsU | -0.179 | 1.91E-01 | -1.144 | 9.54E-02 | -2.105 | 4.33E-03 |
| Q93EP0 | 50S ribosomal protein L21 | rplU | -0.133 | 1.88E-01 | -1.425 | 5.68E-02 | -2.115 | 2.69E-04 |
| A0A0U1MNU9 | Uncharacterized protein | yqeM | -0.005 | 9.66E-01 | -0.567 | 4.59E-02 | -2.118 | 8.68E-02 |
| W8U2K6 | Replicative DNA helicase | dnaC | -0.053 | 3.57E-01 | -2.182 | 5.65E-03 | -2.119 | 1.74E-03 |
| P0C1U9 | DNA topoisomerase 4 subunit A | parC | -0.033 | 5.06E-01 | -2.023 | 2.38E-02 | -2.128 | 1.35E-02 |
| A0A0G2LSY7 | Translation initiation factor IF-3 | infC | 0.008 | 9.04E-01 | -0.838 | 7.74E-03 | -2.129 | 3.10E-03 |
| A0A2X3Q6M1 | Carboxylesterase | est | -0.131 | 5.26E-02 | -1.334 | 7.62E-02 | -2.137 | 8.29E-03 |
| A0A1Q8DEB6 | D-alanyl-lipoteichoic acid biosynthesis protein DltD | dltD | -0.895 | 2.11E-02 | -2.044 | 1.14E-01 | -2.138 | 1.70E-03 |
| A0A0U1MP34 | DNA-dependent ATPase | rarA | -0.023 | 6.23E-01 | -2.087 | 3.24E-02 | -2.145 | 7.24E-03 |
| A0A133PW28 | Uncharacterized protein | | 0.031 | 4.84E-01 | -0.559 | 4.86E-03 | -2.151 | 6.75E-04 |
| A0A0Z0K9X2 | Nitrite reductase [NAD(P)H] large subunit | nasD | -0.207 | 2.89E-01 | -1.594 | 1.22E-01 | -2.155 | 9.11E-03 |
| A0A2S6D906 | Multidrug efflux protein (Fragment) | | -0.063 | 7.34E-01 | -0.912 | 3.51E-02 | -2.156 | 1.87E-03 |
| A0A0D6H8Z4 | EVE domain protein | | -0.055 | 8.94E-03 | -2.645 | 6.51E-02 | -2.159 | 2.71E-02 |
| A0A0U1MX93 | Tyrosine-protein phosphatase | capC | -0.073 | 6.22E-01 | -1.632 | 1.53E-02 | -2.159 | 1.53E-03 |
| A0A2C9TUV1 | Tyrosine recombinase | xerC | -0.032 | 7.54E-01 | -1.264 | 2.12E-02 | -2.159 | 2.65E-02 |
| A0A077V2E3 | Regulatory protein | R | 0.033 | 3.37E-01 | -1.197 | 1.99E-02 | -2.161 | 1.90E-02 |
| W8TRU5 | mRNA interferase | ydcE | 0.083 | 2.13E-01 | -0.853 | 4.31E-02 | -2.164 | 6.44E-02 |
| A0A077W129 | 30S ribosomal protein S13 | rpsM | -0.276 | 2.66E-01 | -2.034 | 4.07E-02 | -2.170 | 1.93E-02 |

| | | | | | | | | |
|---|---|---|---|---|---|---|---|---|
| A0A0Y9VMJ0 | Uncharacterized protein | | 0.063 | 5.13E-01 | -1.017 | 1.57E-01 | -2.175 | 1.04E-02 |
| A0A0U1MED5 | Putative 4-diphosphocytidyl-2-C-methyl-D-erythritol kinase | ispE | -0.015 | 8.54E-01 | -1.903 | 3.76E-02 | -2.177 | 4.45E-03 |
| A0A133Q5H6 | Hydrolase, alpha/beta domain protein | | -0.047 | 2.53E-01 | -1.451 | 4.04E-02 | -2.183 | 4.74E-03 |
| W8TRE0 | 50S ribosomal protein L18 | rplR | -0.002 | 9.54E-01 | -1.407 | 4.87E-02 | -2.188 | 2.67E-02 |
| A0A0D6HBI8 | [Ribosomal protein S18]-alanine N-acetyltransferase | rimI | -0.026 | 7.52E-01 | -2.092 | 4.50E-03 | -2.191 | 5.93E-02 |
| A0A090LVU9 | Signal recognition particle protein | ffh | -0.076 | 5.85E-01 | -2.058 | 2.88E-02 | -2.192 | 2.60E-02 |
| A0A0U1MLI2 | DnaD domain protein | | -0.100 | 4.69E-01 | -2.173 | 1.02E-02 | -2.193 | 2.90E-02 |
| W8TUC3 | 50S ribosomal protein L5 | rplE | -0.210 | 1.02E-01 | -2.115 | 2.53E-02 | -2.198 | 5.05E-04 |
| A0A0Y9UBB6 | Long-chain-fatty-acid--CoA ligase | fcbA2 | -0.216 | 2.54E-01 | -2.017 | 1.42E-01 | -2.199 | 2.09E-02 |
| A0A090LSV1 | Uncharacterized protein | | 0.062 | 3.22E-01 | -1.992 | 6.00E-02 | -2.211 | 5.81E-03 |
| A0A077UGW7 | 30S ribosomal protein S14 type Z | rpsN | 0.069 | 4.83E-01 | -1.506 | 1.79E-02 | -2.220 | 6.93E-03 |
| A0A077VK84 | Hydrogen peroxide-inducible genes activator | oxyR | -0.010 | 8.48E-01 | -1.182 | 5.85E-03 | -2.224 | 2.25E-02 |
| A0A380EW28 | Succinate-semialdehyde dehydrogenase (NADP+) | gabD | -0.394 | 8.60E-02 | -2.432 | 1.51E-02 | -2.228 | 8.56E-02 |
| A0A0D3Q6J0 | RNA-binding domain protein | ylmH | -0.005 | 9.41E-01 | -1.200 | 2.34E-02 | -2.234 | 1.25E-02 |
| A0A0G2LTJ5 | Shikimate kinase | aroK | 0.131 | 7.01E-02 | -1.847 | 6.53E-02 | -2.242 | 1.02E-02 |
| A0A2I7Y7E4 | Protein translocase subunit | secA | -0.019 | 6.43E-01 | -2.373 | 2.22E-02 | -2.243 | 6.32E-03 |
| A0A380EFG1 | Phosphomethylpyrimidine kinase | pdxK | 0.103 | 4.15E-01 | -2.124 | 3.87E-02 | -2.245 | 4.67E-02 |
| W8TRF0 | Allophanate hydrolase | | -0.135 | 7.05E-02 | -2.035 | 5.53E-02 | -2.260 | 1.77E-02 |
| A0A0U1MN82 | DUF309 domain-containing protein | | 0.055 | 7.17E-01 | -1.390 | 1.12E-01 | -2.261 | 1.86E-02 |
| A0A0S3E242 | Probable DNA-directed RNA polymerase subunit delta | rpoE | 0.095 | 1.97E-01 | -3.400 | 2.01E-02 | -2.261 | 2.96E-02 |
| W8TNU8 | Bifunctional hydroxymethylpyrimidine kinase/phosphomethylpyrimidine kinase | pdxK | -0.106 | 5.61E-02 | -2.552 | 4.82E-02 | -2.262 | 1.67E-02 |
| A0A077UUA0 | 50S ribosomal protein L14 | rplN | -0.125 | 6.34E-02 | -1.182 | 6.96E-02 | -2.265 | 2.65E-02 |
| X5DSR5 | Holliday junction ATP-dependent DNA helicase | ruvB | 0.022 | 3.71E-01 | -2.075 | 2.02E-02 | -2.277 | 4.50E-02 |
| A0A133PUQ6 | ABC transporter, ATP-binding protein | | -0.115 | 1.21E-01 | -1.659 | 7.01E-02 | -2.283 | 4.32E-03 |
| A0A0U1MN78 | Uncharacterized protein | | -0.625 | 2.26E-01 | -1.112 | 4.48E-02 | -2.283 | 5.64E-02 |
| A0A380DGQ7 | Veg protein | | 0.145 | 1.17E-02 | -0.960 | 7.63E-02 | -2.287 | 1.87E-02 |
| A0A0D6G0M0 | Transcription-repair-coupling factor | mfd | -0.077 | 3.66E-01 | -2.742 | 6.58E-04 | -2.292 | 1.36E-03 |
| W8TN11 | FAD-dependent oxidoreductase | yutJ | -0.014 | 6.68E-01 | -2.198 | 4.68E-02 | -2.294 | 5.94E-03 |
| A0A077V0F0 | Protein RecA | recA | -0.065 | 4.42E-01 | -2.715 | 1.76E-02 | -2.302 | 2.89E-03 |
| A0A0D6HK08 | Putative pyruvate, phosphate dikinase regulatory protein | yqfL | 0.028 | 7.30E-01 | -2.340 | 2.55E-02 | -2.305 | 2.73E-02 |
| A0A0Y9XT13 | Dihydroorotate dehydrogenase (quinone) | pyrD | -0.100 | 1.54E-01 | -2.664 | 3.57E-02 | -2.306 | 2.00E-02 |
| A0A380DKV5 | GTP-binding and nucleic acid-binding protein | ychF | -0.135 | 3.29E-01 | -2.262 | 1.32E-02 | -2.308 | 1.39E-02 |
| A0A0U1MMX6 | Glycerophosphoryl diester phosphodiesterase family protein | | 0.062 | 3.48E-01 | -1.468 | 4.60E-02 | -2.322 | 1.91E-02 |
| A0A068A649 | tRNA (guanine-N(7)-)-methyltransferase | trmB | -0.033 | 7.39E-01 | -1.254 | 1.89E-02 | -2.328 | 4.81E-03 |
| A0A1Q8DGG5 | Putative N-acetylmannosamine-6-phosphate 2-epimerase | nanE | -0.111 | 3.89E-01 | -0.972 | 4.66E-02 | -2.335 | 4.94E-02 |
| A0A0E7MV68 | Deoxyribodipyrimidine photo-lyase | phrB | 0.030 | 6.35E-01 | -0.958 | 3.12E-02 | -2.346 | 1.26E-01 |
| P45555 | Chaperone protein | dnaJ | 0.000 | 9.99E-01 | -2.635 | 6.08E-02 | -2.348 | 4.24E-03 |
| A0A2W3M2W5 | Cytosolic protein | | -0.012 | 8.87E-01 | -1.066 | 9.63E-03 | -2.361 | 1.28E-03 |
| A0A2S6DIE4 | Enoyl-[acyl-carrier-protein] reductase [NADPH] | fabI | -0.037 | 5.66E-01 | -2.334 | 4.83E-02 | -2.369 | 8.34E-03 |
| A0A077UZP7 | Transcriptional repressor | nrdR | -0.030 | 6.75E-01 | -2.269 | 2.51E-04 | -2.371 | 4.15E-02 |
| A0A380DR43 | SCP family extracellular protein | | -0.246 | 2.59E-01 | -2.200 | 7.48E-02 | -2.372 | 1.40E-02 |
| A0A1L0UI21 | Protoporphyrinogen oxidase | hemY | 0.180 | 1.18E-01 | -3.022 | 8.02E-02 | -2.373 | 3.91E-03 |
| P0A0A8 | Aminoacyltransferase | femB | 0.001 | 9.82E-01 | -1.825 | 3.22E-02 | -2.378 | 2.87E-02 |
| W8UXC3 | 30S ribosomal protein S20 | rpsT | 0.092 | 1.11E-01 | -1.227 | 4.30E-02 | -2.388 | 3.39E-04 |
| A0A380E3U6 | Pyruvate kinase | pykA | -0.469 | 3.54E-01 | -1.745 | 3.79E-02 | -2.393 | 2.48E-02 |
| A0A0D6H3B7 | KR domain-containing protein | yghA | -0.014 | 2.99E-01 | -2.189 | 1.19E-01 | -2.399 | 1.23E-02 |
| Q93PV8 | Translation elongation factor Tu (Fragment) | tuf | -0.011 | 9.16E-01 | -1.138 | 1.59E-02 | -2.408 | 6.87E-02 |
| A0A077VYH1 | Uncharacterized protein | | -0.258 | 3.60E-01 | -1.035 | 4.21E-02 | -2.411 | 3.76E-03 |

| | | | | | | | | |
|---|---|---|---|---|---|---|---|---|
| A0A2X2JYE9 | Capsular polysaccharide synthesis enzyme Cap5G UDP-N-acetylglucosamine 2-epimerase | capG | 0.013 | 8.20E-01 | -2.397 | 5.65E-02 | -2.415 | 2.49E-02 |
| A0A077UYM9 | DNA repair protein | radA | -0.096 | 2.00E-01 | -2.246 | 1.25E-01 | -2.419 | 4.79E-02 |
| A0A1D4Q3C3 | Gluconate operon transcriptional repressor | ydfH | -0.167 | 3.48E-01 | -1.846 | 7.30E-02 | -2.431 | 1.50E-02 |
| A0A2S6D3H4 | Octopine dehydrogenase | odh | 0.004 | 9.31E-01 | -2.101 | 2.38E-02 | -2.432 | 9.68E-03 |
| A0A2S6DIV4 | 50S ribosomal protein L6 (Fragment) | | 0.081 | 1.88E-01 | -1.173 | 8.16E-04 | -2.432 | 3.85E-02 |
| A0A0U1MDZ2 | GNAT family acetyltransferase | | 0.187 | 4.61E-01 | -2.631 | 4.89E-02 | -2.435 | 1.40E-02 |
| A0A0U1MJQ2 | Putative glycosyl transferases | tagE | -0.078 | 5.11E-01 | -1.407 | 6.76E-03 | -2.436 | 1.48E-02 |
| Q9LAB5 | Immunodominant staphylococcal antigen B | isaB | -0.082 | 3.68E-02 | -1.645 | 6.62E-02 | -2.437 | 1.09E-02 |
| W8U126 | DUF47 domain-containing protein | ykaA | -0.081 | 1.38E-01 | -1.847 | 4.06E-02 | -2.456 | 4.28E-02 |
| A0A380DG06 | ATP-dependent RecD-like DNA helicase | recD | 0.143 | 3.63E-01 | -2.180 | 6.22E-04 | -2.457 | 2.75E-02 |
| A0A3S4MW48 | Putative plasmid replication-associated protein | soj | -0.069 | 5.73E-01 | -0.413 | 2.05E-02 | -2.468 | 3.29E-04 |
| A0A077UKD6 | 30S ribosomal protein S11 | rpsK | -0.002 | 9.65E-01 | -1.361 | 1.28E-03 | -2.472 | 2.61E-04 |
| A0A133QCN6 | Putative glycerate kinase | | -0.151 | 2.54E-01 | -2.188 | 6.95E-02 | -2.473 | 1.30E-02 |
| A0A077UKZ2 | Esterase/lipase | mlhB | 0.040 | 2.66E-01 | -2.853 | 1.78E-02 | -2.474 | 4.10E-02 |
| A0A380DE26 | Putative radical SAM family protein | miaB | -0.049 | 6.86E-01 | -1.478 | 1.20E-03 | -2.480 | 8.12E-02 |
| A0A447ZS66 | Bacillithiol system oxidoreductase, YphP/YqiW family | | -0.103 | 3.82E-01 | -1.703 | 6.33E-02 | -2.485 | 1.88E-03 |
| W8TSV3 | Probable GTP-binding protein | engB | 0.035 | 6.40E-01 | -1.352 | 1.16E-03 | -2.485 | 2.82E-02 |
| A0A380ENP7 | 30S ribosomal protein S11 | rpsK | -0.219 | 3.86E-01 | -1.614 | 1.07E-01 | -2.497 | 1.19E-02 |
| Q9RL83 | SepS16A | | -0.005 | 9.05E-01 | -1.701 | 4.25E-03 | -2.499 | 3.61E-02 |
| A0A3S4N1T4 | TrmH family tRNA/rRNA methyltransferase YacO | yacO | -0.114 | 1.54E-02 | -1.577 | 2.91E-02 | -2.503 | 5.24E-02 |
| W8UBI1 | Ribosomal RNA small subunit methyltransferase H | mraW | 0.080 | 4.09E-01 | -1.442 | 6.81E-02 | -2.507 | 4.13E-02 |
| A0A133PVD0 | DNA translocase FtsK | sftA | -0.003 | 9.56E-01 | -2.553 | 6.49E-04 | -2.513 | 2.00E-04 |
| A0A077VGK5 | Aluminum resistance protein | | 0.026 | 7.12E-01 | -1.168 | 2.70E-02 | -2.513 | 1.97E-03 |
| A0A0D6HN81 | 5'-3' exonuclease | ypcP | 0.050 | 2.26E-01 | -3.240 | 6.36E-02 | -2.515 | 4.54E-02 |
| A0A077VZA6 | Conserved virulence factor B | cvfB | -0.121 | 5.78E-02 | -1.293 | 9.80E-02 | -2.521 | 2.61E-02 |
| A0A166KDV7 | LysR family HTH-type transcriptional regulator | cynR | -0.157 | 4.13E-02 | -1.340 | 3.83E-02 | -2.524 | 1.83E-02 |
| A0A077UK83 | Adapter protein | mecA | 0.072 | 2.21E-01 | -1.941 | 7.26E-02 | -2.525 | 4.96E-02 |
| A0A077W1W4 | Teichoic acids export ATP-binding protein | tagH | -0.083 | 4.68E-01 | -2.154 | 3.12E-03 | -2.529 | 3.00E-03 |
| A0A1D5A9I0 | DNA polymerase III beta subunit | dnaN | 1.870 | 5.95E-02 | 0.215 | 5.65E-01 | -2.536 | 4.73E-02 |
| A0A133PWN9 | Uncharacterized protein | | 0.043 | 6.38E-01 | -1.724 | 9.87E-02 | -2.544 | 1.53E-02 |
| W8UX97 | Uridine kinase | udk | -0.056 | 2.73E-01 | -2.379 | 2.89E-02 | -2.545 | 3.95E-02 |
| A0A077VMP6 | 50S ribosomal protein L20 | rplT | -0.082 | 2.73E-01 | -0.967 | 3.69E-02 | -2.545 | 6.69E-04 |
| A0A0E7XUD2 | 3-beta hydroxysteroid dehydrogenase | | 0.039 | 2.81E-01 | -2.300 | 3.48E-02 | -2.545 | 7.11E-03 |
| A0A2C9TMI7 | UPF0176 protein D1G01_02095 | | 0.102 | 4.59E-01 | -1.980 | 3.10E-02 | -2.557 | 5.46E-02 |
| A0A380ES29 | M protein trans-acting positive regulator (MGA) PRD domain | | -0.145 | 3.64E-01 | -1.799 | 2.00E-02 | -2.558 | 2.63E-02 |
| A0A0Y8XVL1 | Ribulose-5-phosphate reductase | tarJ | 0.117 | 1.03E-01 | -2.204 | 6.43E-03 | -2.566 | 4.98E-02 |
| A0A2S6D5J1 | 30S ribosomal protein S7 (Fragment) | | -0.179 | 1.49E-01 | -1.602 | 3.07E-02 | -2.569 | 2.74E-02 |
| A0A0T7MWK2 | Ribonuclease P protein component | rnpA | -0.594 | 1.34E-01 | -2.240 | 1.86E-02 | -2.589 | 1.20E-01 |
| A0A0D6HDD9 | Protoporphyrinogen oxidase | hemY | -0.010 | 7.67E-01 | -2.722 | 3.78E-02 | -2.599 | 2.30E-03 |
| A0A0U1MJ63 | Methylenetetrahydrofolate--tRNA-(uracil-5-)-methyltransferase | trmFO | -0.021 | 5.84E-02 | -2.527 | 2.69E-02 | -2.604 | 1.86E-03 |
| A0A0D6HUA0 | Glycosyl transferase family 1 | tagE | -0.083 | 2.29E-01 | -1.911 | 5.79E-02 | -2.610 | 2.44E-02 |
| A0A0U1MH16 | Choline ABC transporter (ATP-binding protein) | opuBA | 0.115 | 3.78E-01 | -2.696 | 1.08E-03 | -2.613 | 1.18E-01 |
| A0A0E7NNB9 | Nitroreductase | | -0.044 | 3.38E-01 | -1.789 | 1.01E-02 | -2.620 | 1.23E-02 |
| A0A455X0X3 | Uncharacterized protein | | -0.130 | 2.47E-01 | -2.010 | 5.70E-02 | -2.625 | 3.12E-03 |
| D2JA15 | 6-phosphogluconate dehydrogenase, decarboxylating | | 0.015 | 8.64E-01 | -1.046 | 1.30E-02 | -2.629 | 1.82E-02 |
| A0A0D6HXC4 | ATP-binding cassette domain-containing protein | oppF | 0.053 | 6.55E-01 | -2.772 | 1.57E-02 | -2.637 | 2.80E-02 |
| A0A380EFX7 | Phosphoribosylaminoimidazolesuccinocarboxamide synthase | purK | -0.219 | 2.72E-01 | -2.454 | 1.22E-02 | -2.643 | 6.60E-03 |
| A0A133Q8Z9 | 50S ribosomal protein L25 | rplY | 0.037 | 5.66E-01 | -2.399 | 5.20E-02 | -2.645 | 2.00E-02 |

| | | | | | | | | |
|---|---|---|---|---|---|---|---|---|
| A0A1Q8DFG7 | Fatty acid-binding protein DegV | degV | -0.402 | 9.94E-02 | -2.545 | 2.57E-02 | -2.650 | 7.88E-03 |
| A0A0E7DLE1 | Phosphatidylglycerol--prolipoprotein diacylglyceryl transferase | lgt | -0.218 | 9.40E-02 | -2.179 | 8.20E-03 | -2.650 | 6.80E-03 |
| A0A0U1MTR7 | Short-chain dehydrogenase/reductase family oxidoreductase | | 0.002 | 9.72E-01 | -2.105 | 1.04E-01 | -2.650 | 7.86E-03 |
| W8TRX1 | Polyribonucleotide nucleotidyltransferase | yabR | -0.045 | 5.18E-01 | -1.361 | 6.89E-02 | -2.652 | 1.82E-02 |
| A0A380ER18 | Chromosome partition protein | smc | -0.108 | 5.16E-01 | -2.795 | 3.10E-02 | -2.659 | 2.21E-02 |
| A0A133PZQ5 | 30S ribosomal protein S12 | rpsL | -0.193 | 4.37E-02 | -2.428 | 4.84E-02 | -2.679 | 3.06E-04 |
| A0A0U1MG17 | Ribonuclease M5 | rnmV | 0.108 | 7.99E-02 | -1.854 | 6.88E-02 | -2.691 | 1.38E-04 |
| A0A077UL47 | 50S ribosomal protein L35 | rpmI | 0.042 | 8.78E-01 | -1.513 | 9.15E-02 | -2.692 | 7.51E-03 |
| A0A0D6HYA6 | Ribonuclease R | rnr | -0.014 | 7.31E-01 | -1.828 | 4.82E-02 | -2.696 | 5.39E-06 |
| A0A0D6HJL9 | Elongation factor 4 | lepA | 0.057 | 4.10E-01 | -2.466 | 5.16E-02 | -2.705 | 1.98E-02 |
| A0A2S6DIY3 | Carbamoyl phosphate synthase small subunit (Fragment) | | -0.172 | 4.87E-01 | -1.869 | 4.16E-02 | -2.710 | 1.77E-02 |
| A0A0D6H514 | DEAD-box ATP-dependent RNA helicase | cshB | 0.001 | 9.74E-01 | -1.533 | 6.72E-04 | -2.722 | 5.86E-03 |
| A0A0D6HFE2 | 2-succinylbenzoate--CoA ligase | menE | 0.196 | 1.76E-01 | -2.215 | 6.38E-02 | -2.723 | 2.55E-02 |
| A0A0D6HVX5 | Phosphoribosylformylglycinamidine synthase subunit | purL | 0.096 | 7.81E-03 | -2.794 | 3.71E-02 | -2.724 | 1.42E-02 |
| A0A0D6HQ28 | DNA mismatch repair protein | mutL | -0.075 | 1.52E-01 | -2.327 | 4.09E-02 | -2.738 | 2.61E-04 |
| A0A0D6HL28 | 5-formyltetrahydrofolate cyclo-ligase | yqgN | -0.089 | 3.67E-01 | -2.285 | 1.01E-02 | -2.739 | 5.16E-02 |
| A0A0D6W8U6 | Acetyltransferase | rimL | -0.114 | 4.31E-01 | -1.765 | 3.99E-04 | -2.748 | 2.66E-02 |
| A0A0D6HDA2 | DUF1727 domain-containing protein | | -0.027 | 7.90E-01 | -2.369 | 2.35E-03 | -2.755 | 1.40E-02 |
| A0A0U1MZJ5 | Uncharacterized protein | | -0.087 | 2.97E-01 | -2.426 | 1.99E-02 | -2.777 | 1.25E-02 |
| H9BRQ8 | Phenol-soluble modulin PSM-alpha-4 | psm alpha-4 | -0.071 | 8.46E-01 | -1.556 | 1.33E-02 | -2.785 | 9.26E-03 |
| Q9X665 | ABC transporter ATP-binding protein | fhuC | -0.279 | 5.83E-02 | -2.634 | 6.60E-02 | -2.785 | 2.20E-02 |
| A0A077UHB7 | Biotin synthase | bioB | -0.019 | 9.49E-01 | -2.473 | 3.56E-02 | -2.796 | 5.23E-02 |
| A0A0U1MEZ3 | Transcriptional regulator, BadM/Rrf2 family | | -0.228 | 9.34E-02 | -2.582 | 6.36E-02 | -2.800 | 4.41E-02 |
| P0C1S1 | DNA mismatch repair protein | mutS | -0.028 | 4.92E-01 | -2.326 | 5.17E-02 | -2.802 | 9.36E-04 |
| A0A0D6H048 | 8-amino-7-oxononanoate synthase | | -0.014 | 9.66E-01 | -2.566 | 1.79E-02 | -2.804 | 2.94E-02 |
| X5DZV5 | Uncharacterized protein | | 0.337 | 1.27E-01 | -2.078 | 1.31E-02 | -2.817 | 2.93E-02 |
| Q936F1 | Uncharacterized HTH-type transcriptional regulator | | 0.001 | 9.75E-01 | -2.347 | 3.12E-02 | -2.830 | 4.18E-02 |
| A0A1Q8DB85 | Pseudouridine synthase | | 0.234 | 1.41E-01 | -2.027 | 9.42E-02 | -2.831 | 1.29E-02 |
| V5YQH8 | Response regulator | vicR | 0.012 | 9.32E-01 | -2.923 | 1.72E-02 | -2.835 | 1.98E-03 |
| A0A0G2LPK9 | Transcriptional regulator CtsR | ctsR | 0.080 | 2.63E-02 | -2.845 | 6.39E-02 | -2.840 | 7.79E-03 |
| I7A6D3 | DNA topoisomerase IV subunit B (Fragment) | parE | -0.433 | 1.01E-01 | -2.823 | 3.04E-03 | -2.841 | 3.23E-02 |
| A0A380DJ37 | PurR: transcription regulator associated with purine metabolism | purR | 0.007 | 8.75E-01 | -2.687 | 1.01E-02 | -2.844 | 2.37E-03 |
| A0A133PWZ6 | Uncharacterized protein | | 0.145 | 6.18E-02 | -0.359 | 3.17E-03 | -2.852 | 2.29E-02 |
| A0A077UL97 | UPF0473 protein ERS140147_00959 | | 0.180 | 2.79E-01 | -3.047 | 2.34E-02 | -2.855 | 4.27E-04 |
| A0A133Q698 | Oxidoreductase | | 0.040 | 2.62E-02 | -2.962 | 3.38E-02 | -2.872 | 1.74E-02 |
| A0A0D6HLE6 | Segregation and condensation protein B | scpB | 0.057 | 4.10E-01 | -2.734 | 2.37E-02 | -2.889 | 9.88E-03 |
| A0A380EQM1 | Ribonuclease R | rnr | -0.027 | 8.45E-01 | -1.786 | 5.60E-03 | -2.892 | 4.63E-02 |
| A0A0D6HS77 | Pseudouridine synthase | rluD | -0.035 | 5.11E-01 | -2.525 | 2.39E-02 | -2.896 | 1.47E-02 |
| A0A133PYK7 | DeoR family transcriptional regulator | | -0.204 | 8.95E-02 | -2.267 | 9.26E-02 | -2.903 | 2.14E-02 |
| A0A0U1MQP6 | Putative ABC transporter (ATP-binding protein) | ydiF | -0.015 | 7.68E-01 | -2.862 | 3.53E-02 | -2.907 | 7.37E-04 |
| A0A447ZWY8 | Molybdenum cofactor biosynthesis protein B | moaB | -0.097 | 4.59E-01 | -1.112 | 1.21E-02 | -2.910 | 5.48E-03 |
| A0A068A2G4 | LysR family transcriptional regulator | gltC | -0.034 | 8.20E-01 | -1.950 | 9.07E-03 | -2.915 | 9.27E-03 |
| A0A0U1MJN3 | Endonuclease MutS2 | mutSB | -0.070 | 1.38E-01 | -2.821 | 3.44E-02 | -2.916 | 1.98E-02 |
| A0A2S6DGR5 | FAD-dependent oxidoreductase | | 0.040 | 3.32E-01 | -2.371 | 6.89E-02 | -2.924 | 3.12E-02 |
| A0A380EQV1 | Uncharacterized protein | | 0.028 | 2.96E-01 | -2.909 | 8.76E-02 | -2.933 | 4.67E-02 |
| A0A0D6HBJ2 | DNA polymerase III polC-type | polC | 0.183 | 2.00E-01 | -2.349 | 1.23E-02 | -2.935 | 5.70E-02 |
| A0A0U1MV96 | Uncharacterized protein O | | -0.024 | 7.37E-01 | -1.711 | 1.01E-03 | -2.943 | 2.05E-02 |
| C9E0D4 | Factor essential for methicillin resistance (Fragment) | femB | -0.324 | 9.14E-02 | -3.046 | 9.55E-02 | -2.952 | 3.87E-03 |

| | | | | | | | | |
|---|---|---|---|---|---|---|---|---|
| A0A2W2T506 | GTPase | obg | 0.031 | 5.77E-01 | -3.016 | 4.52E-02 | -2.953 | 2.00E-04 |
| A0A0U1MY33 | Phosphoribosylformylglycinamidine synthase subunit | purL | 0.141 | 1.74E-01 | -4.399 | 4.11E-02 | -2.972 | 8.63E-02 |
| A0A077US01 | ABC transporter | | 0.019 | 6.84E-01 | -2.334 | 2.49E-02 | -2.978 | 5.14E-04 |
| A0A133Q1H9 | Uncharacterized protein | | -0.336 | 1.10E-01 | -1.506 | 2.75E-02 | -3.013 | 5.55E-04 |
| A0A0U1ME45 | Secretory antigen SsaA | ssaA2 | -0.036 | 7.00E-01 | -2.533 | 6.08E-02 | -3.017 | 3.48E-02 |
| W8TUP6 | 3-hydroxyacyl-[acyl-carrier-protein] dehydratase | fabZ | -0.212 | 1.66E-01 | -0.647 | 2.12E-01 | -3.022 | 7.30E-03 |
| A0A0D6HQT9 | ATP-dependent DNA helicase | recG | -0.019 | 4.99E-01 | -2.692 | 3.52E-02 | -3.029 | 1.98E-03 |
| A0A380DJ06 | Acid phosphatase | hel | -0.126 | 4.40E-01 | -1.950 | 5.41E-02 | -3.032 | 2.31E-03 |
| P0A0A5 | Aminoacyltransferase | femA | 0.019 | 7.48E-01 | -2.136 | 6.07E-02 | -3.033 | 1.32E-03 |
| A0A090LZ30 | Putative 3-methyladenine DNA glycosylase | | -0.111 | 2.58E-01 | -3.236 | 2.90E-02 | -3.034 | 3.89E-02 |
| A0A2X2KAD1 | Lipoyl synthase | lipA | -0.180 | 3.05E-01 | -1.157 | 3.97E-03 | -3.040 | 4.08E-02 |
| A0A133Q4H9 | Inositol monophosphatase family protein | | -0.108 | 6.41E-01 | -3.649 | 7.79E-02 | -3.055 | 1.79E-03 |
| A0A3S4Q242 | Peptide chain release factor 1 | prfA | -0.353 | 3.31E-01 | -1.804 | 3.45E-02 | -3.069 | 8.07E-03 |
| A0A1L1JA81 | S-adenosylmethionine:tRNA ribosyltransferase-isomerase | queA | -0.407 | 3.25E-01 | -2.287 | 1.22E-02 | -3.074 | 2.59E-02 |
| A0A447ZVX3 | Protein SprT-like | | 0.052 | 4.73E-01 | -1.873 | 3.93E-02 | -3.075 | 6.04E-03 |
| A0A0U1MWH0 | Uncharacterized protein | | -0.324 | 1.00E-01 | -1.353 | 1.47E-02 | -3.079 | 7.34E-03 |
| A0A090N1X6 | DNA repair protein | recO | 0.049 | 8.49E-02 | -3.125 | 7.59E-04 | -3.079 | 4.56E-02 |
| W8TQE6 | 16S rRNA (Guanine(966)-N(2))-methyltransferase | rsmD | 0.023 | 5.19E-01 | -2.100 | 2.49E-02 | -3.085 | 1.33E-02 |
| P21222 | 30 kDa neutral phosphatase (Fragment) | | -0.100 | 1.63E-02 | -1.387 | 4.41E-02 | -3.132 | 2.25E-05 |
| A0A1D3V4E8 | 6-carboxyhexanoate--CoA ligase | bioW | 0.111 | 7.59E-01 | -2.610 | 6.96E-02 | -3.139 | 2.63E-02 |
| A0A3S4LDE9 | Heme chaperone HemW | hemN | -0.077 | 5.33E-01 | -1.835 | 4.82E-03 | -3.144 | 3.21E-03 |
| A0A0H2I3D5 | AraC family regulatory protein | marA | 0.050 | 5.86E-01 | -2.924 | 2.66E-02 | -3.154 | 7.55E-03 |
| A0A222UAN2 | Protein RecA (Fragment) | recA | -0.496 | 1.14E-01 | -3.743 | 7.59E-02 | -3.160 | 1.64E-02 |
| A0A0D6G7M1 | Uncharacterized protein | | 0.181 | 3.84E-01 | -2.971 | 4.25E-02 | -3.162 | 1.72E-02 |
| A0A077V135 | Peptide deformylase | def1 | -0.091 | 3.69E-01 | -1.415 | 3.87E-03 | -3.163 | 1.33E-01 |
| A0A2C9TLS8 | Coenzyme A biosynthesis bifunctional protein CoaBC | | 0.000 | 9.97E-01 | -3.437 | 1.12E-02 | -3.172 | 4.22E-03 |
| A0A2X2KU00 | DNA topoisomerase 4 subunit B | parE | -0.012 | 7.14E-01 | -2.765 | 3.85E-02 | -3.174 | 1.32E-02 |
| A0A0E7XZB8 | tRNA modification GTPase | mnmE | -0.102 | 1.46E-01 | -2.006 | 4.55E-02 | -3.183 | 2.44E-02 |
| A0A133Q0M2 | Isoprenyl transferase | | -0.045 | 4.75E-01 | -2.220 | 5.22E-03 | -3.206 | 9.24E-03 |
| X5E161 | Heat-inducible transcription repressor | hrcA | 0.102 | 2.35E-01 | -2.699 | 1.21E-02 | -3.237 | 8.00E-04 |
| A0A0D6HN86 | Adenosylmethionine-8-amino-7-oxononanoate aminotransferase | bioA | -0.012 | 9.64E-01 | -2.736 | 2.77E-02 | -3.248 | 2.12E-02 |
| A0A0D6GS23 | Esterase family protein | | -0.192 | 1.48E-01 | -3.104 | 8.06E-03 | -3.254 | 3.51E-02 |
| A0A0D3Q6B8 | Peptide chain release factor 3 | prfC | 0.075 | 3.36E-01 | -3.057 | 2.53E-02 | -3.257 | 2.32E-02 |
| A0A077UZF4 | 30S ribosomal protein S18 | rpsR | -0.099 | 3.11E-01 | -2.124 | 5.42E-02 | -3.269 | 9.05E-03 |
| A0A0D6HND3 | SWIM zinc finger domain protein | | 0.002 | 9.79E-01 | -3.436 | 4.63E-02 | -3.274 | 3.65E-02 |
| A0A077VJU6 | 30S ribosomal protein S15 | rpsO | 0.165 | 6.86E-02 | -1.473 | 1.65E-03 | -3.275 | 1.26E-02 |
| A0A0U1MU00 | Esterase/lipase | mlhB | 0.214 | 6.02E-02 | -4.206 | 1.47E-01 | -3.294 | 1.39E-03 |
| A0A389Q0L6 | Putative lipoprotein | | 0.139 | 5.20E-01 | -3.219 | 5.86E-03 | -3.312 | 7.25E-03 |
| C4TF80 | Uncharacterized protein | | -0.294 | 2.58E-01 | -1.258 | 7.01E-02 | -3.330 | 7.60E-03 |
| A0A0U1MI46 | Putative lipoprotein | | -0.702 | 1.73E-01 | -2.995 | 2.60E-02 | -3.333 | 1.96E-02 |
| A0A2S6DE36 | 30S ribosomal protein S7 (Fragment) | | -0.156 | 9.40E-02 | -2.531 | 3.22E-03 | -3.343 | 1.08E-02 |
| A0A389VCH8 | Ketol-acid reductoisomerase (NADP(+)) | ilvC | -0.069 | 6.00E-01 | -3.152 | 3.37E-02 | -3.345 | 5.39E-03 |
| A0A2S6DU15 | MurR/RpiR family transcriptional regulator | | -0.024 | 5.52E-01 | -2.654 | 3.81E-04 | -3.358 | 1.04E-03 |
| A0A077W1R3 | Antitoxin MazE | ydcD | 0.241 | 1.90E-01 | -3.842 | 8.88E-02 | -3.373 | 2.12E-03 |
| A0A2C9TH70 | Ribonuclease III (Fragment) | | 0.149 | 1.82E-01 | -0.957 | 2.75E-02 | -3.387 | 4.99E-04 |
| A0A2I7Y5R0 | UPF0346 protein BTN44_05455 | | 0.148 | 1.72E-01 | -1.678 | 9.02E-02 | -3.388 | 2.96E-02 |
| A0A133Q5S2 | Uncharacterized protein | | 0.010 | 8.60E-01 | -2.127 | 8.65E-02 | -3.405 | 4.63E-02 |
| A0A0D6HZA6 | Carboxylesterase | est | -0.062 | 6.03E-01 | -3.552 | 3.43E-02 | -3.416 | 1.48E-03 |

| | | | | | | | | |
|---|---|---|---|---|---|---|---|---|
| A0A2S6DTY7 | Capsular biosynthesis protein | | -0.194 | 2.03E-01 | -3.291 | 5.38E-02 | -3.423 | 2.11E-02 |
| W8TVX7 | GTPase Era | bex | -0.155 | 9.83E-02 | -3.003 | 5.20E-02 | -3.439 | 2.61E-03 |
| W8TVT7 | Collagenase-like protease | yhbU | 0.048 | 4.13E-01 | -3.846 | 5.83E-02 | -3.444 | 3.27E-02 |
| A0A0D6GUP8 | Alpha/beta hydrolase | mlhB | 0.088 | 1.49E-01 | -3.418 | 6.13E-03 | -3.486 | 1.59E-03 |
| P0A0I7 | Accessory gene regulator protein A | agrA | -0.038 | 1.27E-01 | -2.308 | 4.22E-04 | -3.521 | 2.21E-02 |
| A0A0D6H808 | DUF3885 domain-containing protein | | -0.102 | 1.46E-01 | -2.976 | 6.05E-02 | -3.562 | 2.27E-02 |
| W8U450 | DUF2750 domain-containing protein | | -0.084 | 1.32E-01 | -3.419 | 7.69E-02 | -3.620 | 1.72E-02 |
| A0A133Q130 | Oxygen-dependent choline dehydrogenase | betA | -0.190 | 2.84E-01 | -3.681 | 5.86E-02 | -3.702 | 1.01E-02 |
| A0A1D4J866 | Sensor histidine kinase | nreB | -0.147 | 4.55E-02 | -3.900 | 1.83E-02 | -3.741 | 2.35E-02 |
| A0A2S6D886 | 50S ribosomal protein L25/general stress protein Ctc (Fragment) | | 0.087 | 3.99E-01 | -2.851 | 6.43E-02 | -3.750 | 3.50E-02 |
| A0A380DZS3 | CopZ | copZ | -0.065 | 3.32E-01 | -2.752 | 7.16E-03 | -3.817 | 1.35E-02 |
| A0A0U1MM88 | Peptidase, U32 family | yrrN | 0.009 | 7.35E-01 | -3.684 | 5.07E-02 | -3.825 | 5.63E-03 |
| A0A380EE99 | Dihydropteroate synthase | folP | -0.009 | 9.02E-01 | -3.564 | 4.71E-02 | -3.863 | 1.32E-02 |
| A0A077VHR5 | Phosphomevalonate kinase | mvaK2 | -0.064 | 1.48E-01 | -3.563 | 3.11E-02 | -3.882 | 3.02E-02 |
| A0A077ULY5 | Ornithine cyclodeaminase / Siderophore staphylobactin biosynthesis protein | sbnB | -0.087 | 6.16E-01 | -3.335 | 1.44E-02 | -3.893 | 1.66E-02 |
| A0A1L0ZJS8 | D-aminoacyl-tRNA deacylase | dtd | -0.318 | 9.20E-02 | -3.305 | 3.46E-02 | -3.901 | 2.13E-02 |
| A0A0D6GXG5 | ABC transporter ATP-binding protein | ybbL | 0.061 | 5.74E-02 | -4.168 | 4.99E-02 | -3.923 | 1.91E-02 |
| A0A0D6GZF2 | ATP-dependent dethiobiotin synthetase | bioD | -0.028 | 9.40E-01 | -4.008 | 1.35E-02 | -3.945 | 4.03E-02 |
| A0A077VMC3 | General stress protein 13 | yugI | -0.152 | 2.97E-01 | -3.151 | 4.70E-03 | -4.049 | 1.59E-01 |
| A0A2X2MDC9 | UDP-GlcNAc 2-epimerase | mnaA | -0.385 | 4.40E-01 | -2.479 | 1.56E-03 | -4.062 | 6.14E-02 |
| A0A0D6HM65 | Aldo/keto reductase | yhdN | -0.199 | 2.40E-01 | -4.999 | 1.33E-01 | -4.285 | 7.74E-04 |
| A0A1D4IEJ6 | Anthranilate synthase component II | pabA | 0.012 | 9.03E-01 | -4.156 | 6.16E-02 | -4.303 | 6.64E-04 |
| A0A0D6HS97 | Glyoxalase | catE | 0.034 | 5.70E-01 | -4.058 | 8.26E-02 | -4.450 | 2.95E-02 |
| A0A133Q6E7 | Heme response regulator HssR | hssR | -0.098 | 1.62E-01 | -2.961 | 1.26E-02 | -5.132 | 1.10E-01 |
| A0A0U1MBW0 | GCN5-related N-acetyltransferase (GNAT) | | 0.075 | 2.18E-01 | -2.332 | 2.02E-03 | -5.187 | 7.89E-04 |
| A0A0U1MFR2 | tRNA1(Val) (Adenine(37)-N6)-methyltransferase | trmN | -0.001 | 9.87E-01 | -4.325 | 8.08E-03 | -5.261 | 7.16E-02 |
| A0A0E8GB80 | Tetratricopeptide repeat protein | | 0.009 | 6.14E-01 | -4.609 | 6.18E-02 | -5.544 | 2.31E-02 |
| A0A2X2KNH5 | Phage protein | | -0.207 | 6.54E-02 | -4.136 | 8.89E-02 | -6.099 | 4.69E-02 |

LogFC, average log2 fold change (n=2). Heat map applied to LogFC values - shading in red indicate upregulation and blue indicate downregulation of protein abundance levels compared to control samples. Statistically significant values shown in black font ($p < 0.05$) and insignificant values ($p > 0.05$) in grey font as determined by *t*-test. Proteins with no gene names assigned have been left blank.
